# BRIDGE: Backbone Representation for Inverse-folding-Driven Generalizable RNA Encoding

**DOI:** 10.64898/2026.11.23.768775

**Authors:** Jinung Song, Junsu Ko, Juyong Lee, Woong-Hee Shin

## Abstract

RNA recognizes small molecules and proteins through their three-dimensional structures, but sparse functional annotations limit task-specific learning of transferable structural features. We present BRIDGE, an inverse-folding encoder pretrained on 2,501 RNA-chain records from experimental structures. Each nucleotide is represented by its phosphorus atom and five ribose atoms (P, C1′, C2′, C3′, C4′, O4′). Base coordinates are not encoder inputs. From these atoms, the encoder is trained to predict nucleotide identity, solvent-accessible surface area (SASA) and base geometry. With matched readouts, frozen BRIDGE representations reach a three-model ensemble pooled average precision (AP) of 0.627 for external ligand-binding site prediction and 0.647 for RNA–protein interface prediction, compared with 0.503 and 0.516 for frozen, randomly initialized encoders. Under the evaluated full-adaptation recipes, pretrained initialization reaches AP 0.703 for ligand binding and 0.641 for protein interfaces, versus 0.635 and 0.623 for training from scratch. Sequence-only pretraining, without SASA or base-geometry targets, also transfers to both tasks. Whether the base-geometry objectives add further value depends on the task and adaptation mode. In decoy ranking and apo/holo evaluations, transfer depends on the readout and the test set, and small target panels and downstream training overlap in the apo/holo evaluation limit broader conclusions. These results support inverse-folding pretraining as a route to transferable RNA structural representations that do not require observed base coordinates at inference.

## Introduction

RNA recognizes small molecules, proteins and other molecular partners through its three-dimensional structure. Riboswitches change conformation when they bind metabolites [1, 2], ribosomal RNA forms the binding sites of many antibiotics [3, 4], and HIV-1 TAR RNA rearranges its structure when it binds arginine, the residue at the center of its recognition by the Tat protein [5]. Structured RNAs also contain pockets that can bind drug-like small molecules [6]. Benchmark sets of annotated RNA binding sites, however, contain only a few hundred chains [7]. Deep models trained on these labels alone therefore have few examples from which to learn the structural features that underlie recognition, and representations learned from unlabeled RNA structures could complement such task-specific training.

Inverse folding can learn structural features without functional labels, because the native sequence serves as a target. In proteins, ESM-IF1 and Protein-MPNN learn structure-conditioned sequence preferences [8, 9], and their representations transfer beyond sequence design. IF-SitePred and DiscoTope-3.0 use ESM-IF1 features to predict ligand-binding residues and B-cell epitopes [10, 11]. ThermoMPNN transfers ProteinMPNN representations to mutation-induced stability prediction, and SPURS addresses the same task by combining structural and sequence representations through supervised adaptation [12, 13]. These studies use pretrained representations both as fixed features and within adapted models, so we evaluate both frozen and task-adapted encoders.

RNA inverse-folding models have been developed mainly for sequence design. They include the data-efficient RDesign, the aptamer-oriented RhoDesign and the diffusion-based RiboDiffusion [14, 15, 16]. gRNAde conditions design on one or more RNA conformations, and NA-MPNN models nucleic-acid sequence preferences within a unified biopolymer graph [17, 18]. RDesign and gRNAde train on thousands of RNA structures. RiboSphere also learns RNA structure representations, by coordinate reconstruction rather than inverse folding, and combines them with sequence, secondary-structure and ligand features to predict RNA–ligand interactions at the pair level [19]. It therefore needs a query ligand, whereas binding-site prediction locates binding residues without one.

Here, we ask whether RNA inverse-folding representations transfer to molecular recognition. We introduce BRIDGE (Backbone Representation for Inverse-folding-Driven Generalizable RNA Encoding), an encoder pretrained by inverse folding with auxiliary geometric supervision (Figure 1A). For each nucleotide, BRIDGE takes the phosphorus atom, five ribose atoms and a deterministic sugar-derived virtual nitrogen as input and produces a per-nucleotide structural representation. Native base atoms are excluded because they would reveal which nucleotide is being reconstructed, so base geometry enters only as a pre-training target. We evaluate transfer of the pretrained encoder to ligand-binding site and RNA–protein interface prediction using task-specific readouts. We further test whether the ligand-binding models recover holo-defined sites from apo conformations, as binding-site prediction must do when the bound state is unknown. We also test whether the same representation supports RNA structure ranking, a graph-level task that depends on the quality of a whole structural model rather than on individual binding residues (Figure 1B).

**Figure 1.**
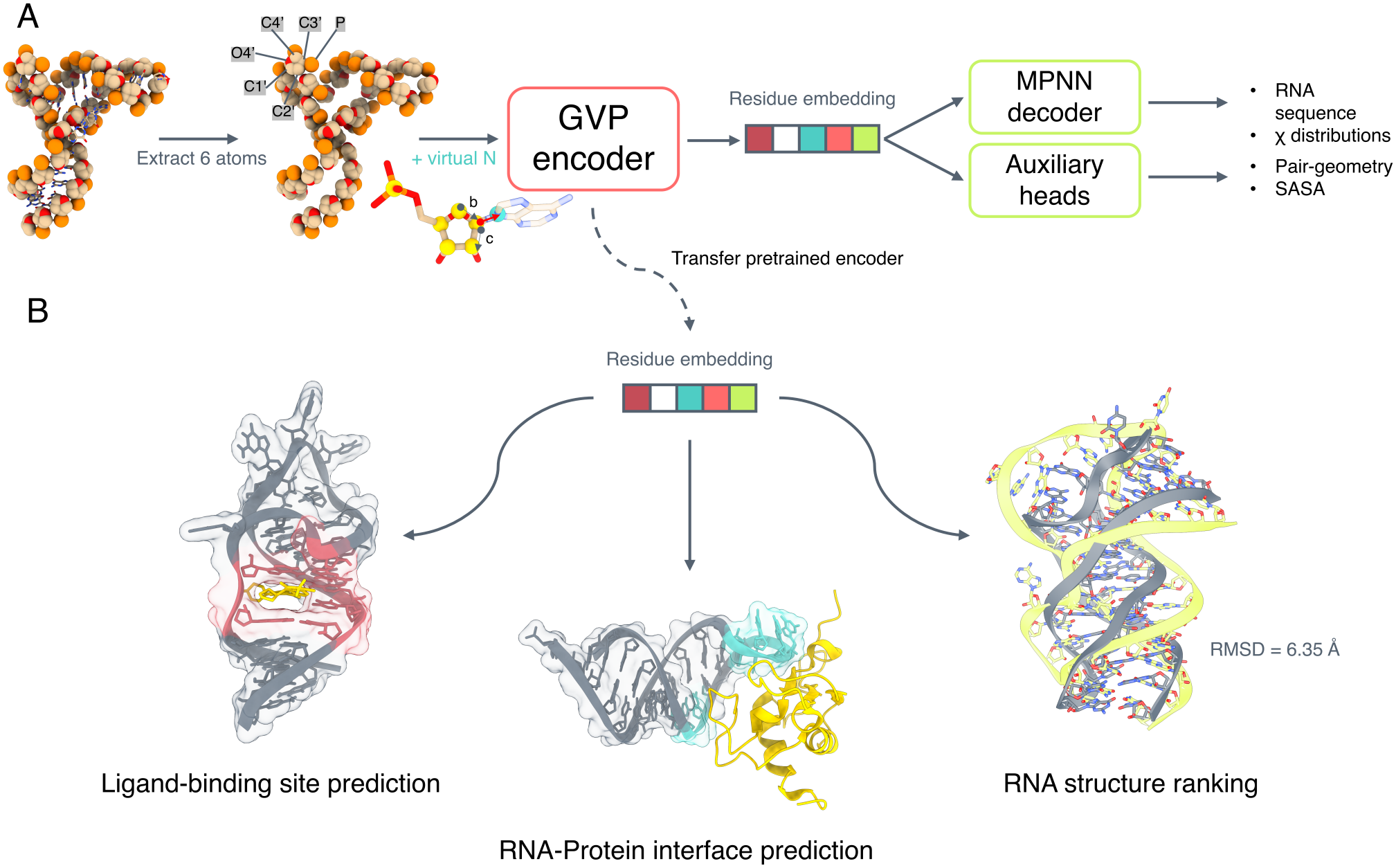
BRIDGE pretraining and downstream representation transfer. **(A)** The phosphorus atom and five ribose atoms of each nucleotide (P, C1′, C2′, C3′, C4′ and O4′), supplemented by a deterministic sugar-derived virtual nitrogen, are encoded by a four-layer geometric vector perceptron (GVP) network. In the inset, vectors **b** (O4′ →C1′) and **c** (C1′ →C2′), together with their plane normal, determine the virtual-N position. A two-layer causal message-passing decoder predicts nucleotide identities and glycosidic-angle (*X*) distributions. Separate linear auxiliary heads predict solvent-accessible surface area (SASA) and pairwise base geometry directly from the encoder representations. Observed base coordinates provide pretraining supervision but are not encoder inputs. **(B)** The pretrained encoder is transferred to ligand-binding site prediction, RNA–protein interface prediction and RNA structure ranking through task-specific readouts. The pretraining decoder and auxiliary heads are not transferred. The ranking example superposes a native structure and a decoy from RNA-Puzzle 21, with a heavy-atom RMSD of 6.35 Å. The native structure is used only for evaluation.

To separate what pretraining contributes from what a downstream model can learn on its own, we use three matched comparisons. Comparing frozen pretrained and random-frozen encoders with identical readouts asks whether pretraining makes useful information available. Pretrained and scratch encoders trained with a shared full-adaptation recipe test the value of initialization. Objective-removal variants, with architecture and inputs held fixed, measure the contribution of local and pairwise base-geometry supervision. We apply these comparisons to ligand-binding sites and RNA–protein interfaces. Released gRNAde and NA-MPNN encoders are included for context.

## Methods

### Backbone inputs and pretraining objectives

BRIDGE observes six atoms per nucleotide: P, C1′, C2′, C3′, C4′ and O4′. A deterministic virtual nitrogen is constructed from the sugar frame using fixed NA-MPNN coefficients [18, 20]. Observed N1/N9 coordinates and presence masks are erased before feature construction; invalid frames are masked. Native base geometry is available only as supervision, not as an encoder input. The virtual-point formula, atom slots and geometric conventions are specified in Supplementary Section 1.4.

A four-layer geometric vector perceptron (GVP) encoder [21], with scalar/vector widths 128/16 and dropout 0.1, produces a 128-dimensional representation per nucleotide. Pretraining uses C4′-kNN32 edges and sequence adjacency. A two-layer causal decoder, motivated by LASErMPNN [22, 23], predicts nucleotide identity and four nucleotide-specific, 24-bin glycosidic-angle (*X*) distributions. During training, only neighbours preceding the current nucleotide in a random decoding order contribute native sequence context [9]. Source diagnostics instead use one soft predicted-prefix argmax rollout, marginalizing angle predictions over predicted nucleotide probabilities. Linear heads predict solvent-accessible surface area (SASA) and symmetric pairwise base geometry directly from encoder states.

Sequence cross-entropy uses label smoothing 0.05. Absolute SASA targets are standardized using training statistics and fitted by masked mean squared error. PairGeom predicts minimum base-heavy-atom distance in seven bins and C1′–glyN–glyN–C1′ torsion in 24 circular bins. Eligible pairs are existing undirected graph pairs separated by at least four sequence positions with valid target atoms. Local-angle and pair cross-entropies are normalized by training-set prior entropies. The complete objective is

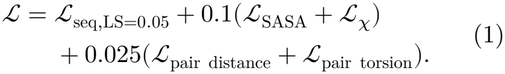

S0 uses sequence and SASA supervision; S0+PairGeom and S0+*X* add the indicated geometry objective. G0 combines both and is the complete-objective reference. G1 retains these losses and adds preceding-angle context to the decoder. Removing an objective preserves architecture, initialization, data and duration; remaining losses are not renormalized. The sequence-only control also removes SASA, retaining inactive auxiliary modules to preserve paired initialization. SASA carries base-dependent information despite restricted encoder inputs, so this additional control distinguishes sequence reconstruction from all auxiliary supervision.

Each variant uses three independent pretraining seeds, 100 epochs and the final checkpoint, with FP32 AdamW, learning rate 2 × 10^−4^, weight decay 10^−4^, gradient clipping at 1 and a 1,500-node batch cap. Records longer than 500 nucleotides are excluded, not cropped. Only encoder weights transfer; the decoder and auxiliary heads are discarded. Random-frozen trains a readout over a fixed random encoder, whereas Scratch-N trains both from random initialization; N denotes the virtual-N input. Loss weights were not systematically tuned. Model roles and the emphasis on epoch 100 were chosen retrospectively after ear-lier checkpoint evaluations. Restricted-input transfer does not establish superiority over other atom sets or robustness to missing coordinates. Full implementation, capacity and timing details are in Supplementary Sections 1 and 16.

### Datasets and exposure control

Pretraining uses the RNASolo-derived corpus processed for gRNAde [17, 24], with 2,501/82/68 training/validation/test records after length and expo-sure filtering. Registered downstream test and external identities were screened by provenance PDB and normalized exact sequence. Exclusions propagated through shared source identities and provenance PDBs; identified RNA-Puzzle 21 relatives were also excluded. This identity-based screen does not establish comprehensive homology or family independence. Downstream training and validation structures remain without functional labels, so validation-based model selection is not strictly inductive. Later RNA3D-Decoy evaluation used a retrospective exposure audit.

Supplementary Table S1 distinguishes pretraining exposure from downstream fitting. Source-test record 3GS5_1_C shares training provenance; its exclusion leaves 67 records/5,532 nucleotides and is reported as a saved-score sensitivity in Supplementary Table S9. The separate common-26 subset excludes PDB matches to both released inverse-folding inventories; it is not comprehensive sequence/family clearance (Supplementary Table S48). Complete memberships and exclusion audits are in Supplementary Sections 1.7, 1.9, 7 and 8.1.

### Downstream adaptation and model selection

#### Ligand-binding sites

The supplied combined HARIBOSS–TR60 benchmark contains 331/41/43 training/validation/test chains [7]; it is not the original 60-RNA TR60 collection alone [25]. BRIDGE uses a two-layer width 128 C4′-kNN16 GNN read-out and weighted binary cross-entropy for 120 epochs. Full adaptation and Scratch-N share inputs, architecture and selection, with encoder/head learning rates 10^−4^/10^−3^. LoRA uses rank 4, scale 2 and learning rate 10^−3^, so this comparison also changes encoder learning rate. Validation AP is evaluated every five epochs, retaining the earliest best checkpoint. Frozen conditions fit only the readout.

The RBind-derived RB9 subset [26], JL10 and TL12 [27], and TE18 [25] contain 9/10/12/17 records after filtering. TE18 excludes 4F8U; TL12 retains training-overlapping 8F0N, with an exclusion sensitivity reported separately. The external pool contains 48 records/44 PDBs and 2,369 residues. Shared RB9/TE18 chains retain their supplied labels and duplicated contribution. Labels are the supplied 4 Å contact arrays; all included positions are retained and missing geometry is masked. Upstream chemical annotation policies were not independently reconstructed.

#### RNA–protein interfaces

RNAProt CLEAN filters the existing RNAglib partitions [28] by PDB identity: retain all test graphs, remove validation PDB overlaps with test, then remove training overlaps with either retained partition. Removing 20 validation and 54 training graphs yields 835/171/191 graphs, with 10,780 test residues from 177 PDB units. Filtering does not consult test labels. GNN readouts are evaluated for 40 epochs using pretraining seed 0 and three downstream fits; GAT results are supplementary (Table S31). A separate three-pretraining-seed objective-removal experiment is kept distinct from this readout comparison.

#### Structure ranking

ATOM3D RSR [29] contains 12,479/4,000/4,000 training/validation/test decoys; test Puzzles 18–21 contribute 1,000 each. Mean-pooling MLP and residual GNN readouts regress RMSD with MSE for 20 epochs and validation selection, fixing pretraining epoch 100/seed 0. Metrics are computed separately for three downstream seeds and then averaged. Figure 2E/F shows target-level performance; complete numeric results are in Supplementary Tables S32 and S33.

**Figure 2.**
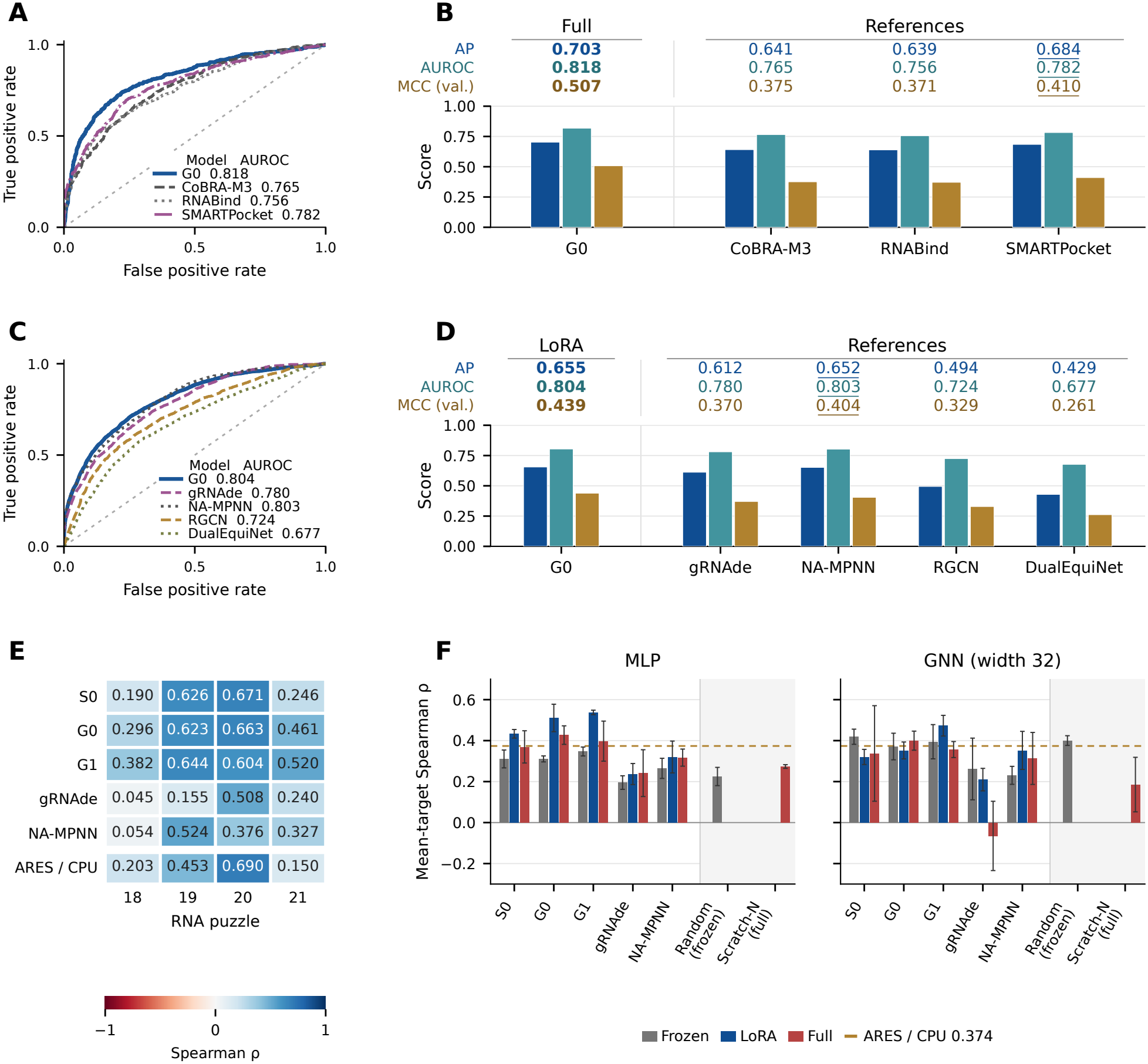
Transfer across RNA recognition and structure-ranking tasks. **(A, C)** ROC curves for the ligand external pool (48 records, 44 PDBs; 2,369 residues, 773 positives) and RNAProt CLEAN test set (191 graphs, 177 PDB units; 10,780 residues, 2,980 positives). Curves use three-fit mean probabilities; SMARTPocket averages five stochastic passes of one released model. A and C use the same models and adaptation settings as B and D, respectively: G0 uses full adaptation in A/B and LoRA in C/D. **(B, D)** Pooled average precision (AP), AUROC and Matthews correlation coefficient at validation-selected thresholds (MCC val). B compares full-adaptation G0 with CoBRA-M3, RNABind and SMARTPocket; D compares G0 with LoRA against full-adapted gRNAde and NA-MPNN and the local RGCN and DualEquiNet baselines. Both G0 configurations use the GNN readout. These panels show selected comparisons; the full adaptation and control panels remain in Tables 2 and 4. Rows above the bars give point estimates to three decimals, without confidence intervals. Bold and underline mark the best and second-best distinct unrounded values among the displayed conditions within each panel and metric; exact ties share emphasis. Rankings are descriptive. References share test support but differ in inputs, training data and budgets. **(E)** Heatmap of per-target Spearman *ρ* between predicted and reference RMSD for MLP+LoRA on RNA puzzles 18–21 (1,000 correlated decoys each), averaged across three fits. This configuration was emphasized post hoc, with retrospective validation support for S0/G0/G1, not as a prespecified primary analysis. **(F)** Equally weighted mean-target *ρ* for MLP and GNN readouts across frozen, LoRA and full adaptation. Error bars show SD across three fits on the same four targets, not target-sampling confidence intervals. MLP mean-target correlations and seed SD are reported in Table 5; pooled correlations and GNN results are in Supplementary Tables S32 and S33. Random-frozen and Scratch-N/full are shown only where evaluated; absent bars were not run. ARES is a single released atomic predictor without MLP/LoRA or fit-seed SD; its dashed reference is *ρ* = 0.374. G0 is the complete-objective reference, S0 an ablation and G1 a variant.

RNA3D-Decoy and CASP16 provide additional ranking evaluations [30]. RNA-only fitting excludes CASP-overlapping targets and AF3-generated decoys: 329 targets/40,138 decoys train the models, while validation retains 40 targets/6,120 decoys including AF3. Twenty-epoch fits select the earliest maximum validation target-macro Spearman; CASP16 is not used for selection. Labels are sequence-registered, common-heavy-atom RMSDs after rigid alignment, minimized over included native conformers. Native coordinates define labels only. RNA3D38 (38 targets/5,814 decoys) is a post-hoc exposure-filtered subset, not an un-touched or comprehensively family-independent test; CASP16 retains 20 targets/3,789 submissions. Detailed exclusions and additional readouts are in Supplementary Sections 7 and 10–12.

**Table 1.** Pretraining diagnostics on 68 source-test records at epoch 100, averaging three pretraining seeds.

| Model | Recovery | *A*<sub>30</sub> | MAE (°) |
|---|---|---|---|
| Sequence-only | 0.6088 | – | – |
| S0 | 0.6219 | – | – |
| S0+PairGeom | 0.6245 | – | – |
| S0+χ | **0.6276** | **0.9527** | 10.6661 |
| G0 | 0.6238 | 0.9489 | **10.5779** |
| G1 | 0.6181 | 0.9486 | 10.6773 |
| Training prior | – | 0.8909 | 15.0389 |

Recovery and *A*<sub>30</sub> are fractions; MAE is angular error (lower is better). Training prior predicts the modal centre (−157.5°) of the training-only global 24-bin soft-count distribution for every residue. It has no test-fitted parameters and is excluded from rank emphasis. Dashes denote untrained/inapplicable outputs. The 67-record exclusion of training-linked 3GS5 and complete angular metrics are in Supplementary Table S9.

**Table 2.**
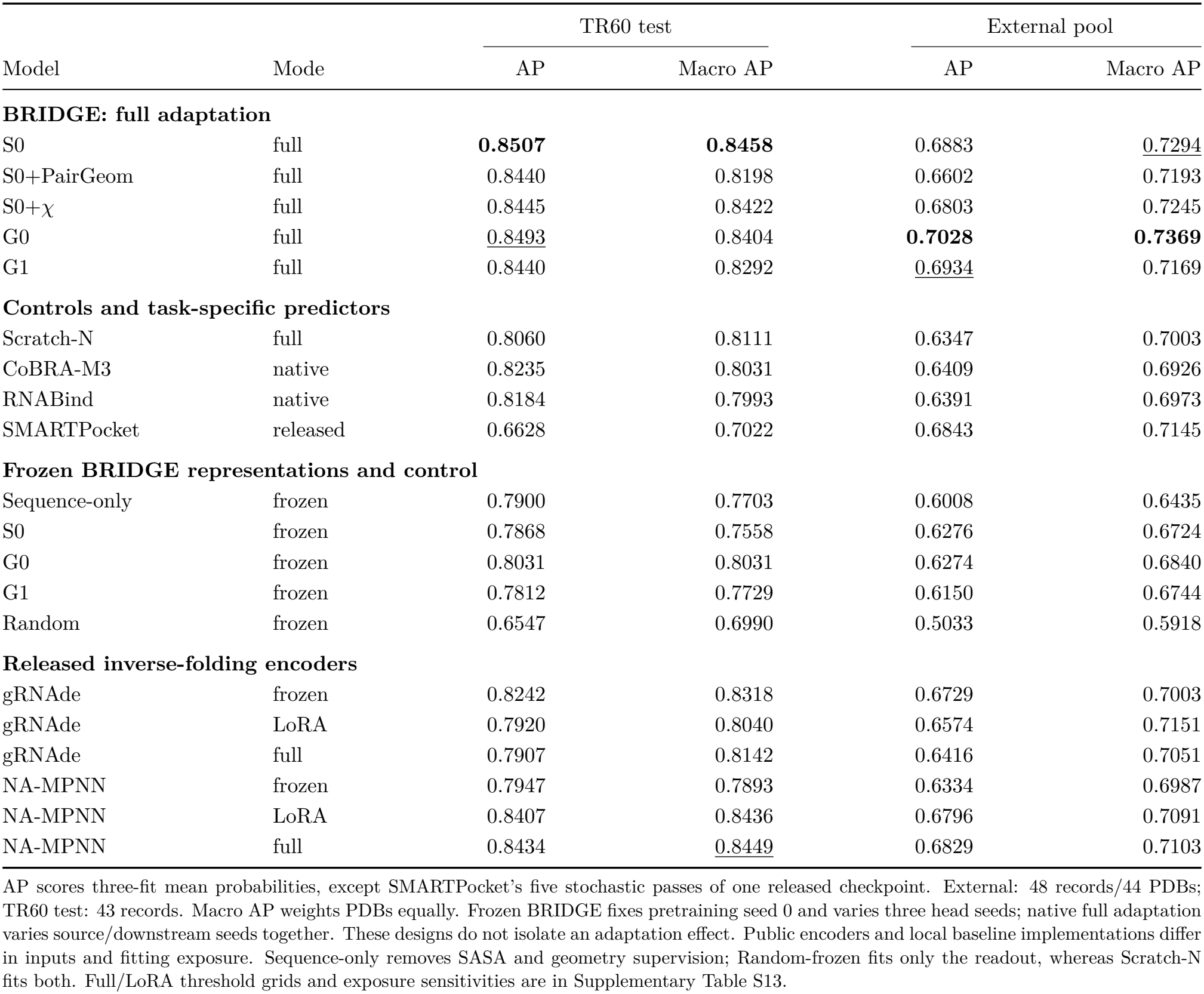
BRIDGE, locally evaluated binding-site methods and released inverse-folding encoders adapted on TR60.

**Table 3.**
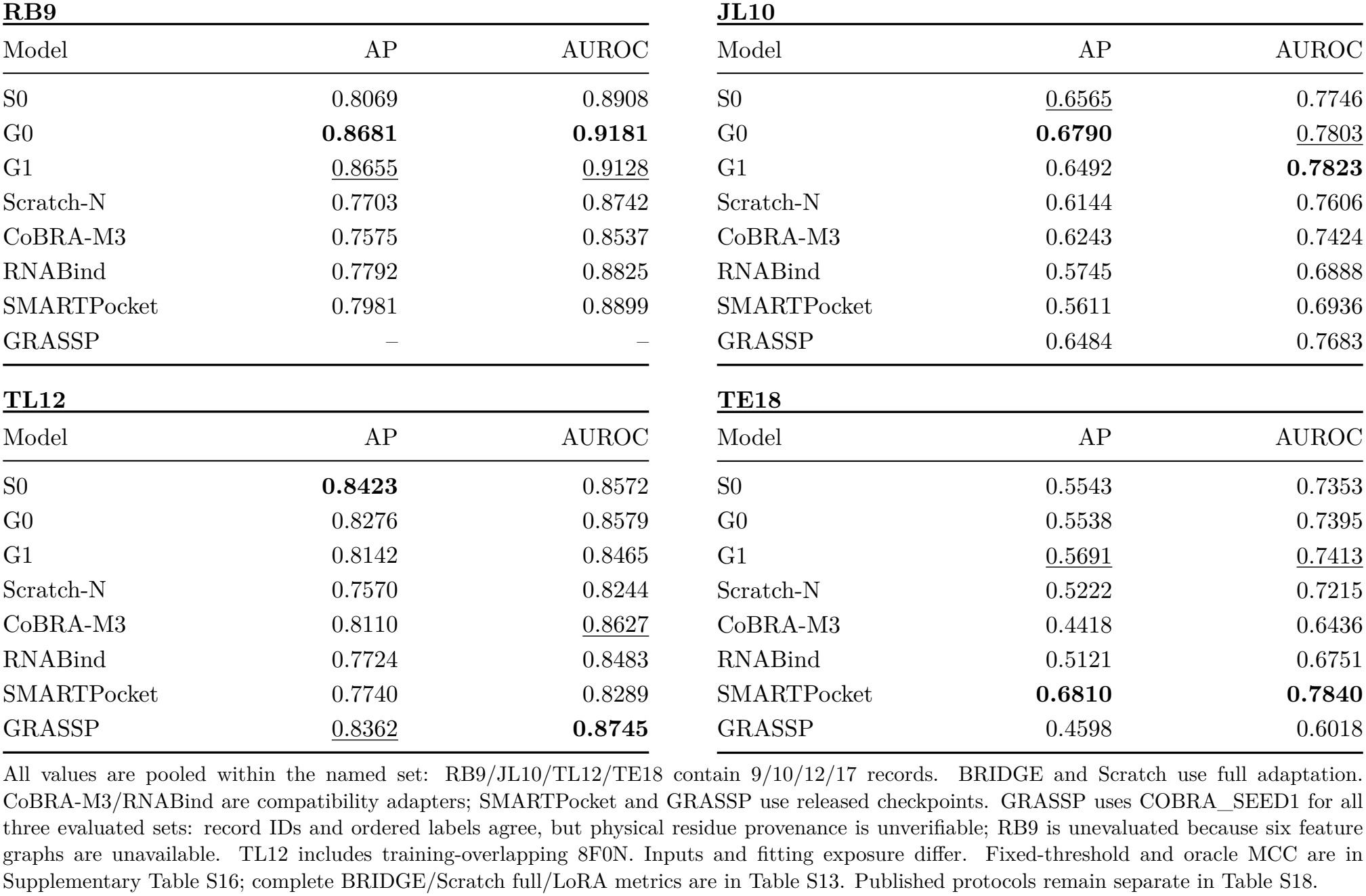
Local evaluation on each external ligand-binding benchmark.

| **RB9** |  |  | **JL10** |  |  |
| Model | AP | AUROC | Model | AP | AUROC |
|---|---|---|---|---|---|
| S0 | 0.8069 | 0.8908 | S0 | <u>0.6565</u> | 0.7746 |
| G0 | **0.8681** | **0.9181** | G0 | **0.6790** | <u>0.7803</u> |
| G1 | <u>0.8655</u> | <u>0.9128</u> | G1 | 0.6492 | **0.7823** |
| Scratch-N | 0.7703 | 0.8742 | Scratch-N | 0.6144 | 0.7606 |
| CoBRA-M3 | 0.7575 | 0.8537 | CoBRA-M3 | 0.6243 | 0.7424 |
| RNABind | 0.7792 | 0.8825 | RNABind | 0.5745 | 0.6888 |
| SMARTPocket | 0.7981 | 0.8899 | SMARTPocket | 0.5611 | 0.6936 |
| GRASSP | – | – | GRASSP | 0.6484 | 0.7683 |
| **TL12** |  |  | **TE18** |  |  |
| Model | AP | AUROC | Model | AP | AUROC |
| S0 | **0.8423** | 0.8572 | S0 | 0.5543 | 0.7353 |
| G0 | 0.8276 | 0.8579 | G0 | 0.5538 | 0.7395 |
| G1 | 0.8142 | 0.8465 | G1 | <u>0.5691</u> | <u>0.7413</u> |
| Scratch-N | 0.7570 | 0.8244 | Scratch-N | 0.5222 | 0.7215 |
| CoBRA-M3 | 0.8110 | <u>0.8627</u> | CoBRA-M3 | 0.4418 | 0.6436 |
| RNABind | 0.7724 | 0.8483 | RNABind | 0.5121 | 0.6751 |
| SMARTPocket | 0.7740 | 0.8289 | SMARTPocket | **0.6810** | **0.7840** |
| GRASSP | <u>0.8362</u> | **0.8745** | GRASSP | 0.4598 | 0.6018 |

All values are pooled within the named set: RB9/JL10/TL12/TE18 contain 9/10/12/17 records. BRIDGE and Scratch use full adaptation. CoBRA-M3/RNABind are compatibility adapters; SMARTPocket and GRASSP use released checkpoints. GRASSP uses COBRA\_SEED1 for all three evaluated sets: record IDs and ordered labels agree, but physical residue provenance is unverifiable; RB9 is unevaluated because six feature graphs are unavailable. TL12 includes training-overlapping 8F0N. Inputs and fitting exposure differ. Fixed-threshold and oracle MCC are in Supplementary Table S16; complete BRIDGE/Scratch full/LoRA metrics are in Table S13. Published protocols remain separate in Table S18.

**Table 4.**
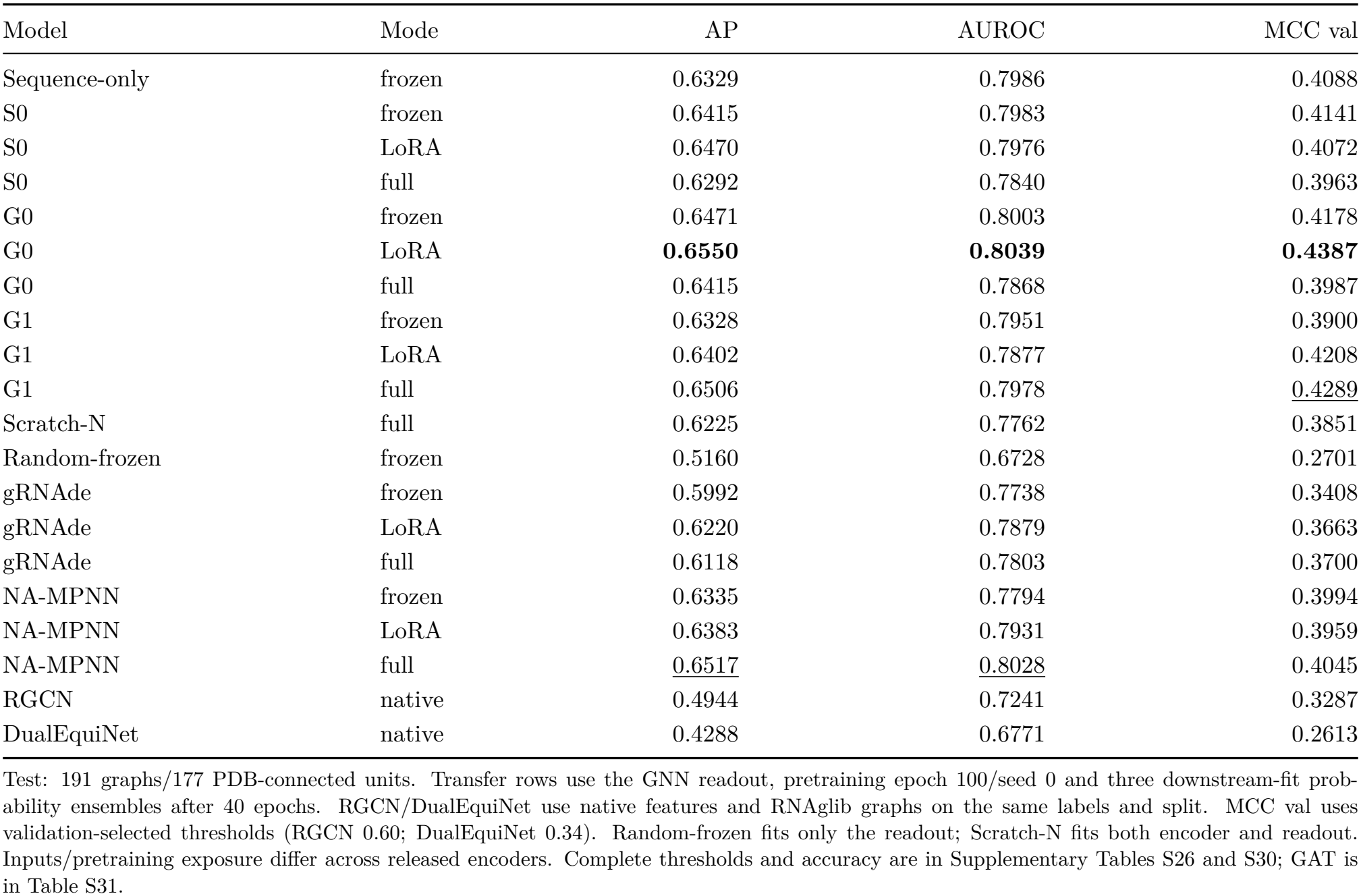
RNAProt CLEAN: GNN transfer readouts and native baselines.

| Model | Mode | AP | AUROC | MCC val |
|---|---|---|---|---|
| Sequence-only | frozen | 0.6329 | 0.7986 | 0.4088 |
| S0 | frozen | 0.6415 | 0.7983 | 0.4141 |
| S0 | LoRA | 0.6470 | 0.7976 | 0.4072 |
| S0 | full | 0.6292 | 0.7840 | 0.3963 |
| G0 | frozen | 0.6471 | 0.8003 | 0.4178 |
| G0 | LoRA | **0.6550** | **0.8039** | **0.4387** |
| G0 | full | 0.6415 | 0.7868 | 0.3987 |
| G1 | frozen | 0.6328 | 0.7951 | 0.3900 |
| G1 | LoRA | 0.6402 | 0.7877 | 0.4208 |
| G1 | full | 0.6506 | 0.7978 | 0.4289 |
| Scratch-N | full | 0.6225 | 0.7762 | 0.3851 |
| Random-frozen | frozen | 0.5160 | 0.6728 | 0.2701 |
| gRNAde | frozen | 0.5992 | 0.7738 | 0.3408 |
| gRNAde | LoRA | 0.6220 | 0.7879 | 0.3663 |
| gRNAde | full | 0.6118 | 0.7803 | 0.3700 |
| NA-MPNN | frozen | 0.6335 | 0.7794 | 0.3994 |
| NA-MPNN | LoRA | 0.6383 | 0.7931 | 0.3959 |
| NA-MPNN | full | 0.6517 | 0.8028 | 0.4045 |
| RGCN | native | 0.4944 | 0.7241 | 0.3287 |
| DualEquiNet | native | 0.4288 | 0.6771 | 0.2613 |

Test: 191 graphs/177 PDB-connected units. Transfer rows use the GNN readout, pretraining epoch 100/seed 0 and three downstream-fit prob-ability ensembles after 40 epochs. RGCN/DualEquiNet use native features and RNAglib graphs on the same labels and split. MCC val uses validation-selected thresholds (RGCN 0.60; DualEquiNet 0.34). Random-frozen fits only the readout; Scratch-N fits both encoder and readout. Inputs/pretraining exposure differ across released encoders. Complete thresholds and accuracy are in Supplementary Tables S26 and S30; GAT is in Table S31.

**Table 5.** ATOM3D RNA structure ranking with the MLP readout: mean-target Spearman *ρ*.

| Encoder | Frozen | LoRA | Full |
|---|---|---|---|
| S0 | 0.3106 ± 0.0436 | 0.4333 ± 0.0209 | 0.3694 ± 0.0787 |
| G0 | 0.3107 ± 0.0146 | 0.5107 ± 0.0670 | **0.4270** ± 0.0452 |
| G1 | **0.3470** ± 0.0218 | **0.5377** ± 0.0109 | 0.3975 ± 0.0977 |
| gRNAde | 0.1953 ± 0.0332 | 0.2369 ± 0.0514 | 0.2413 ± 0.1142 |
| NA-MPNN | 0.2646 ± 0.0492 | 0.3201 ± 0.0778 | 0.3170 ± 0.0418 |
| Random-frozen | 0.2249 ± 0.0445 | – | – |
| Scratch-N | – | – | 0.2737 ± 0.0090 |
| **Released atomic reference** |  |  |  |
| ARES [36] | 0.3739 (released scores) |  |  |

Values are mean±sample SD across three downstream fits, each scored by equally weighted Spearman correlation across four RNA puzzles (1,000 decoys each). The MLP has 16,641 parameters and is fitted for 20 epochs, with checkpoint selection by validation mean-target correlation. Native pretraining seed 0 and released public checkpoints are fixed. Seed SD measures fitting variability, not target-sampling uncertainty; four targets limit population-level inference. Bold and underline mark the best and second-best distinct means within each adaptation column; SD is not ranked. Rankings are descriptive. Random-frozen fits only the readout; Scratch-N/full trains encoder and readout. Dashes indicate unevaluated conditions. ARES uses one released all-atom model and archived scores, without fit-seed SD or adaptation modes; it is separate from the fresh CPU replay in Figure 2. Inputs and training exposure differ across released models. Pooled correlations and the GNN readout are in Supplementary Tables S32–S33; the atom-input refits use the distinct initialization described in Supplementary Section 12.

#### Apo/holo transfer

Six SMARTPocket-derived pairs comprise seven graphs per state, with 605/607 labelled residues and 64 positives per state. Fixed residue mappings project holo labels onto apo structures. This measures recovery of holo-defined sites from apo coordinates, not experimental apo-state binding. Existing TR60 full-adaptation checkpoints and validation thresholds are applied without refitting. Four holo targets occur in downstream training; structural identity does not establish coordinate/label identity. No direct PDB or exact-sequence pretraining-training match was found for the six target identities. Pair mappings and exposure details are in Supplementary Sections 1.9.2, 1.11 and 4.

### Comparators and comparison scope

CoBRA-M3 [7] and RNABind [31] are common-split compatibility adapters, not exact native reproductions. SMARTPocket [32] uses one released check-point averaged over five stochastic passes. GRASSP [33] uses COBRA_SEED1 native feature graphs for JL10/TL12/filtered TE18. Normalized PDB–chain IDs and ordered labels match all 39 common records, but missing residue identifiers/sequences prevent independent physical-residue reconstruction. Training graphs include 8F0N. Six RB9 graphs and a complete raw feature pipeline are unavailable, precluding a common 48-record pooled score. This differs from the published TE18 protocol using a separately trained TR60 model. Published benchmarks are separate contextual comparisons [7, 34, 32, 33].

RGCN [28] and DualEquiNet [35] are trained on CLEAN with native nucleotide features and RNAglib graphs, identical labels/masks, 40 epochs and three initialization seeds. These are local fixed-budget reruns, not reproductions of optimally tuned published settings. Released gRNAde and NA-MPNN encoders [17, 20] are transferred without new pretraining, using the same task readouts, budgets and validation selection. Their inputs differ: gRNAde uses observed glycosidic N and coordinate-derived secondary structure; NA-MPNN uses twelve backbone atoms and virtual N. Public-model training exposure also differs, so these are contextual, not input- and exposure-matched ablations.

Figure 2 reuses saved predictions or replays selected checkpoints without refitting; ARES [36] is replayed on CPU with its released atomic transform. Additional small/large readouts and matched controls were introduced after earlier benchmark inspection and remain exploratory. Full baseline recipes, check-point provenance and all conditions are in Supplementary Sections 3, 6, 8 and 10–16 and the indexed Supplementary Data. Equal-budget results estimate initialization benefit under a shared recipe. RNAProt learning-rate sensitivity evaluates validation-selected lower-rate checkpoints on the existing test set without changing the primary rate; other tasks lack independently tuned comparisons.

### Metrics, seed design and uncertainty

AP is stepwise average precision, not trapezoidal PR area. Pooled AP ranks all admitted residues together and therefore depends on score comparability across units; it is not a weighted average of within-unit AP. Macro AP weights units equally: PDBs for lig- and binding and paired targets for apo/holo, pooling chains within each state. The ligand pool retains set-qualified duplicate records. RNAProt’s supplementary positive-support macro AP includes 122 units; it excludes 55 all-negative units and is not a full-177-unit macro estimate. Definitions and equations are in Supplementary Sections 1.12–1.13.

MCC uses a fixed 0.5 threshold, a validation-selected threshold, or a test-label oracle. The oracle is a nondeployable diagnostic; target-macro oracles optimize independently within each target. Main classification tables emphasize AP, AUROC and, where applicable, validation-MCC; complete threshold and accuracy results are in the supplementary tables and indexed Supplementary Data. Bold and underline identify the best and second-best distinct displayed estimates within comparable panels and metric columns; ties share emphasis. These descriptive ranks do not establish statistical superiority.

Binary-task ensembles average three fitted probability vectors before scoring; individual-run metric means and sample SDs are separate summaries. CoBRA full/LoRA and apo/holo vary pretraining and downstream seeds together. Frozen Co-BRA fixes encoder seed 0 and varies three head seeds; S0/G0/G1/random-frozen share 120 epochs and paired head initialization. RNAProt fixes pre-training seed 0 and combines three downstream fits. RSR likewise fixes pretraining seed 0 but averages three seed-specific metrics, not predictions. These seed axes differ, so frozen-versus-adapted contrasts do not isolate an adaptation effect.

Mean-target Spearman weights ranking targets equally. Three seeds characterize fitting variability without increasing independent target counts: ATOM3D has four puzzles and apo/holo six targets. Reported paired percentile-bootstrap intervals use 10,000 draws of PDBs, RNAProt PDB-connected units, puzzles, RNA3D targets or paired apo/holo targets; CASP16 resamples sequence–structure clusters. Source diagnostics use 20,000 draws. Intervals condition on saved fitted models, omit retraining variance and are unadjusted for multiple comparisons. A resolved contrast excludes zero under its stated metric and aggregation; an unresolved contrast is not evidence of equivalence. Exact sign-flip sensitivities for selected small-target panels are in Supplementary Section 1.14; numerical intervals are in the task-specific supplementary tables and Supplementary Data. Point estimates without intervals carry no significance claim. Saved-score subsets and post-hoc extensions do not create independent test sets.

## Results

### Pretraining captures sequence preferences and local base geometry

We first examined what the pretrained encoders learn on the pretraining test set, using sequence recovery and glycosidic-angle accuracy. G0/G1 sequence recovery is 0.6238/0.6181 (Table 1). The fraction of predicted glycosidic angles within 30^°^ of the native value is 0.9489/0.9486, versus 0.8909 for a training-derived prior. Variants trained without the *X* loss have untrained angle heads; their outputs are reported only as diagnostics in Supplementary Table S10, not as learned angular performance. The pretraining test set contains one 3GS5-derived record that shares provenance with the pretraining training set. Excluding its 36 nucleotides leaves G0/G1 recovery at 0.6238/0.6179 and 30^°^ accuracy at 0.9488/0.9485 (Supplementary Table S9), so this record does not affect the model ranking.

Sequence-only pretraining reaches recovery 0.6088, compared with 0.6219 for S0, or 0.6092 versus 0.6222 without 3GS5. On the full test set, its paired differences from S0 and G0 remain unresolved. Individual seeds and uncertainty are supplementary.

Released inverse-folding models provide a complementary reconstruction diagnostic (Supplementary Table S48). NA-MPNN reaches a pooled recovery of 70.62% from sampled sequences, versus 62.38% for G0, 61.81% for G1 and 56.75% for gRNAde. On the 26 records whose PDB entries are absent from the pretraining data of both released encoders, the corresponding values are 64.71%, 61.30%, 59.71% and 52.76%, and sequence-only recovery is 58.30%. BRIDGE decodes by soft predicted-prefix argmax rollout, whereas the released models sample categorical sequences at temperature 0.1, and inputs and exposure also differ, so this ordering is contextual. The controlled objective-removal panel below is the stronger within-protocol test of recovery versus transfer.

### Pretrained features improve ligand-binding site prediction

We next tested whether BRIDGE representations improve ligand-binding site prediction. Readouts were trained on the combined HARIBOSS–TR60 split with the encoder frozen, adapted with LoRA or fully adapted. Models were evaluated on the TR60 test set and on a 48-record external pool formed by RB9, JL10, TL12 and TE18. Figure 2A,B compares full-adaptation G0 with reference methods using external-pool ROC curves and pooled metrics, respectively. With the encoder frozen and the same GNN readout, G0 and G1 reach pooled AP 0.6274 and 0.6150 on the external pool, versus 0.5033 for a random-frozen encoder. For G0, the pooled and macro gains are 0.1241 and 0.0921, with paired analyses in the supplement. Because only the head is fitted, this comparison shows that the features transfer without encoder adaptation. At the same 120-epoch budget, S0 reaches pooled/macro AP 0.6276/0.6724, compared with 0.6274/0.6840 for G0. Its gains over random-frozen are resolved under both estimands, while its external differences from G0 remain unresolved, and its TR60 test macro AP is lower (0.7558 versus 0.8031). Seed results and paired contrasts for S0 are reported in Supplementary section *S0 frozen readout at a matched 120-epoch budget*. The complete frozen panel is in Table 2.

Removing SASA supervision does not remove the frozen-transfer signal. Sequence-only pretraining reaches external AP 0.6008 and TR60-test AP 0.7900, compared with 0.5033 and 0.6547 for random-frozen, and the external pooled gain of 0.0975 is resolved by paired PDB analysis. Adding SASA gives S0 an unresolved pooled-AP increment of 0.0268 but a resolved macro increment of 0.0288, so the added value of SASA targets depends on the estimand. All pretraining-seed recovery results and downstream head-seed results are supplementary.

Under the matched full-adaptation protocol, pretraining also improves ligand-binding site prediction. On the external pool, G0 reaches pooled AP 0.7028 versus 0.6347 for scratch, a gain of 0.0681, and macro AP improves by 0.0366. Excluding the known training-overlap PDB 8F0N gives G0 AP 0.6919 and a pooled gain of 0.0700, and removing the three duplicated external chains together with 8F0N leaves a pooled gain of 0.0699. On the TR60 test set, the pooled gain is 0.0433 and the macro gain of 0.0293 is unresolved (Supplementary Table S12). We also varied the readout head (Methods). With width32/64 GNN heads, full adaptation gives external AP 0.6633/0.6542 for G0, 0.6729/0.6759 for G1 and 0.6436/0.6675 for S0, so smaller heads do not uniformly improve AP. Because the main panel also varies pretraining seeds, the difference between the two panels does not isolate the effect of width. Individual-run and probability-ensemble metrics are reported separately in Supplementary Table S14; their model ordering can differ. Full membership, uncertainty and seed-level analyses, and the controls, per-set results and macro AP for these heads, are supplementary.

S0, which omits the explicit *X* and PairGeom losses but retains base-dependent SASA supervision, also improves external pooled AP over scratch, by 0.0535. Under full adaptation, its external difference from G0 (−0.0146) is unresolved, as is its pooled-AP difference on the TR60 test set (+0.0015). With LoRA, S0 reaches external pooled AP 0.6938 versus 0.6678 for G0 (Supplementary Table S13). Removing both geometric losses thus changes full-adaptation AP little, whereas removing *X* or PairGeom alone lowers external AP by 0.0426 or 0.0225. These effects are non-additive, so they do not show that either auxiliary loss is generally necessary. G1–G0 is unresolved for pooled AP (−0.0094) but negative for macro AP (−0.0200).

Transfer varies across the external sets (Table 3). G1 has a higher TE18 point estimate than G0, whereas G0 has the higher pooled external value. We use G0 as the reference model across all external sets. Shared RB9/TE18 chains are grouped for uncertainty estimation.

### BRIDGE is competitive with task-specific predictors but not uniformly best

We then compared BRIDGE with task-specific binding-site predictors and with released inverse-folding encoders on the same splits (Table 2). On the external pool, CoBRA-M3, RNABind and SMART-Pocket reach AP 0.6409, 0.6391 and 0.6843, compared with 0.7028 for G0. On TE18, however, SMART-Pocket reaches AP 0.6810 versus 0.5538 for G0. GRASSP achieves AP 0.8158 on its 44 released test graphs, which we report separately from the shared 43-record TR60 test comparison because the released test set differs and residue-level correspondence has not been verified.

The released-encoder comparison is adaptation-dependent. On the same 48-record external pool, full-adapted gRNAde and NA-MPNN reach AP 0.6416 and 0.6829, compared with G0/G1 0.7028/0.6934. LoRA yields 0.6574/0.6796 for the released models versus G0/G1 0.6678/0.6644. Frozen GNN read-outs yield 0.6729/0.6334 versus 0.6274/0.6150. The BRIDGE advantage under full adaptation therefore does not carry over to the frozen setting, where gR-NAde is higher, or to LoRA, where NA-MPNN is higher. After excluding 8F0N, full-adaptation AP is 0.6919/0.6822 for G0/G1 and 0.6310/0.6701 for gRNAde/NA-MPNN. In a paired analysis, the full-adaptation difference between G0 and NA-MPNN remains unresolved (supplement), and these comparisons do not equalize inputs or pretraining histories. Supplementary Table S18 places nine additional baselines summarized in SMARTPocket Table 1, together with MVRBind, CoBRA, SMARTPocket and GRASSP, under their published protocols [34, 7, 32, 33]. Published TE18 AUROC is 0.745 for MVRBind, 0.746 for GRASSP and 0.781 for SMARTPocket, compared with 0.7395 for G0/full on the 17-record set. GRASSP reports TL12 AUROC 0.874, above G0/full 0.8579. BRIDGE is therefore not the strongest method on every external set. The local comparison in Table 3 keeps the common filtered test sets and makes the remaining input and fitting differences explicit. Local GRASSP AUROC is 0.7683, 0.8745 and 0.6018 on JL10, TL12 and TE18, respectively; these values use the same released CoBRA checkpoint for all three sets and must not be equated with its published TE18 result. Published summaries cannot provide a paired comparison or a pooled external AP without the underlying predictions.

### Frozen pretrained features improve RNA–protein interface prediction

We also evaluated RNA–protein interface prediction on CLEAN, a PDB-disjoint RNAProt split that we constructed (Methods). Figure 2C,D compares G0 with LoRA against reference methods using ROC curves and pooled metrics, respectively. With a GNN head, frozen G0 features reach AP 0.6471 versus 0.5160 for random-frozen features, a gain of 0.1312 computed from unrounded scores (Table 4). Frozen sequence-only features reach AP 0.6329, a resolved gain of 0.1169 over random-frozen, and their difference from frozen S0 (AP 0.6415) remains unresolved. Under full adaptation with the same GNN head, G0 reaches AP 0.6415 versus 0.6225 for scratch (ΔAP 0.0190). With the GAT head, G0 exceeds scratch by 0.0244, a positive but unresolved difference. G0– LoRA–GNN has the highest AP in Table 4 (0.6550). The corresponding GAT result is 0.6574 (Supplementary Table S31), but the difference of 0.0024 is unre-solved. Macro AP over the 122 units with positives is 0.6214 for G0/full and 0.5896 for scratch, and the 55 all-negative units contribute only false-positive-rate diagnostics.

For full GNN, single-run AP mean±SD is 0.6260±0.0083 for G0 versus 0.5294±0.1182 for scratch, whose mean is pulled down by one low-scoring fit (seed APs 0.6093/0.5852/0.3937). Omitting each seed in turn (supplement) shows that dropping this run leaves the scratch ensemble AP at 0.6225 after rounding, so we keep it in the primary result. These runs repeat downstream fitting with pretraining seed 0 fixed, so they do not capture pretraining variability. In a learning-rate sensitivity analysis, full-adapted G0 and scratch reach AP 0.6478/0.6120 at encoder rate 10^−5^. The pretrained advantage remains resolved under this alternative recipe, while the within-model changes from 10^−4^ are unresolved. The main comparison keeps the original learning rate, and conditional intervals are supplementary.

With the same GNN readout, released gRNAde frozen/LoRA/full AP is 0.5992/0.6220/0.6118, and NA-MPNN reaches 0.6335/0.6383/0.6517. G0’s frozen/LoRA values (0.6471/0.6550) are numerically higher, whereas full NA-MPNN is close to G1 (0.6506) and above G0 (0.6415). These comparisons use the GNN head throughout, and the ranking of pretrained encoders changes with the adaptation mode. With GAT heads (Supplementary Table S31), gRNAde frozen/LoRA/full AP is 0.5812/0.6292/0.6268, NA-MPNN reaches 0.6432/0.6532/0.6436, and S0 reaches frozen/full AP 0.6500/0.6525, compared with 0.6292 for its full GNN, so changing the head reorders the models instead of improving all of them. With small GNN heads, G0 LoRA reaches AP 0.6333/0.6570 at width32/64, while random-frozen gives 0.4900/0.4820 and scratch-full 0.5849/0.6120. The supplement reports all modes and both widths, and no test maximum is promoted to the primary configuration.

Native baselines on the same test set extend the protein-interface comparison (Table 4). On the CLEAN test set, RGCN reaches AP/AUROC 0.4944/0.7241 and DualEquiNet reaches 0.4288/0.6771, compared with 0.6471/0.8003 for frozen G0 with the GNN readout. The local results favor G0 numerically under these recipes, while native inputs, architectures and optimization differ. DualEquiNet’s MCC is only 0.0239 at thresh-old 0.5, but increases to 0.2613 at its validation-fixed threshold of 0.34, close to its diagnostic oracle MCC of 0.2665. Its accuracy of 0.7229 is also close to the all-negative baseline of 0.7236, so accuracy alone barely separates it from a trivial predictor. Published RGCN and DualEquiNet values, obtained under their original partitions, are shown separately in Supplementary Table S30 [28, 35]. The native reruns use a fixed 40-epoch budget, with two DualEquiNet checkpoints selected at the final epoch, so they do not establish optimally tuned performance. Seedwise estimates are in Supplementary Data D124 and conditional PDB-level intervals are in Supplementary Table S74.

### Structure ranking depends on adaptation and readout

We next asked whether the representation supports a graph-level task, regressing decoy RMSD on the ATOM3D RSR benchmark (Table 5; Figure 2E,F). With matched full adaptation and the MLP read-out, G0 mean-target Spearman is 0.4270 versus 0.2737 for scratch, a difference of 0.1533 that remains un-resolved. LoRA raises G0 from 0.3107 (frozen) to 0.5107, and G1 with LoRA gives the highest point estimate in Table 5 (0.5377), 0.1907 above its frozen value, with heterogeneous puzzle-specific effects. Because population-level resolution depends on the four test puzzles rather than the 4,000 decoys, exact sign-flip sensitivities do not resolve these differences (supplement).

Under the same MLP readout and 20-epoch budget, BRIDGE variants have higher mean-target correlation point estimates than the released encoders in each adaptation mode (Table 5). With only four test targets and different pretraining exposures, these comparisons remain descriptive.

ARES [36], an atomic equivariant RMSD scorer, provides a further comparison. We aligned its released scores to all 4,000 test decoys by puzzle and tag and verified agreement in RMSD and Rosetta energy. On these decoys, ARES reaches mean-target/pooled Spearman 0.3739/0.4145, versus 0.4270/0.5726 for G0/full and 0.5107/0.5898 for G0/LoRA with the MLP readout. ARES uses one published model and all-atom inputs, whereas BRIDGE values average three downstream-seed metrics, so test identities and labels are aligned but training histories and input representations differ. Effects vary across the four puzzles, and per-target results and uncertainty comparisons are supplementary. Figure 2E,F uses a fresh CPU replay of the released ARES model, yielding mean-target Spearman 0.373846; Supplementary Table S32 retains the archived released-score reference. With the width32 residual GNN readout (Supplementary Table S33), the random-frozen control is strong: its mean-target correlation is 0.4003 versus 0.3712 for frozen G0, while frozen S0 and G1 reach 0.4189 and 0.3947 and G0 full adaptation reaches 0.3998 versus 0.1855 for scratch. A fitted graph read-out can therefore extract ranking information without a pretrained encoder, and these point estimates alone do not establish a pretraining advantage on this four-target panel. The graph readout also does not uniformly improve ranking, since G1 with LoRA reaches 0.4729, below its MLP value of 0.5377. Mean-target and pooled correlations describe different endpoints and can reorder models, and equal-budget comparisons do not establish optimally tuned scratch performance. The MLP83k, residual-GNN83k and width64 comparisons are in the supplementary tables; complete ranking metrics and optimization diagnostics are in Supplementary Data D043–D045 and D117–D119. With the same GNN readout, gRNAde frozen/LoRA/full mean-target Spearman is 0.2617/0.2097/−0.0651, and NA-MPNN reaches 0.2305/0.3523/0.3130. G1/LoRA reaches 0.4729, the highest point estimate within this GNN block, whereas G0/LoRA and NA-MPNN/LoRA are nearly identical at 0.3520 and 0.3523. G0/full is numerically above NA-MPNN/full (0.3998 versus 0.3130), but their paired target-level difference remains unresolved. The negative gRNAde/full result comes from a run that completed training and replayed its validation scores exactly, so it is not an execution failure, although its cause is unknown. Adaptation effects therefore depend on the encoder and readout. The full seed-level, puzzle-level and paired uncertainty results are supplementary.

A separate frozen-readout ablation uses a 100-epoch MLP and three pretraining seeds, with read-out seed 0 fixed (Supplementary subsection *Separate frozen-readout loss ablation*). Mean-target Spearman is 0.2648 for G0, 0.2977 for G1, 0.2691 with S0+PairGeom, 0.2426 with S0+*X* and 0.1979 with S0, so the complete objective (G0) is below both G1 and S0+PairGeom. This panel is not directly comparable to the 20-epoch adaptation panel because the training budget, batch size and seed axis differ. Within the 20-epoch panel, only the joint removal (S0) is tested, and matched individual-loss removals under adaptation remain untested.

### Ranking transfer to RNA3D38 and CASP16 depends on readout and test set

To test ranking beyond the four ATOM3D puzzles, we fitted quality predictors on RNA3D-Decoy with the MLP readout and evaluated them on RNA3D38, a retrospective exposure-screened subset rather than an untouched benchmark, and on CASP16 (Supplementary Table S40). Under full adaptation, G0 reaches mean-target Spearman 0.3120 on RNA3D38 and 0.2857 on CASP16, versus 0.2103 and 0.0310 for scratch. The RNA3D38 difference is unresolved, whereas the CASP16 difference is resolved under cluster resampling. Each fitted configuration is evaluated unchanged on both test sets, without test-set-specific model selection, and all frozen, LoRA and full results are in the supplement.

With full adaptation, released gRNAde and NA-MPNN reach 0.3109 and 0.3771 on RNA3D38 and 0.1604 and 0.2699 on CASP16, so NA-MPNN is higher than G0 on RNA3D38 while G0 is higher on CASP16, and full adaptation is not uniformly the best mode (supplement). Atom inputs, pretraining exposure and native regularization differ between the models, and paired contrasts, results on targets absent from the released encoders’ pretraining data, seed-level results and ARES inference are in the supplement.

With the MLP readout, a random-frozen control (Methods) limits pretraining attribution: RNA3D38 mean-target correlation is 0.2902/0.2646 for G0/G1 versus 0.3097 for random features, and CASP16 is 0.1185/0.1207 versus 0.1087. These matched frozen contrasts remain unresolved.

We also applied the larger RSR readouts MLP83k and ResidualGNN83k, keeping the MLP results as the main analysis. With ResidualGNN83k, frozen G0 exceeds random on RNA3D38 (0.3444 versus 0.2483) and CASP16 (0.1851 versus 0.0654) in the conditional paired analysis. Full G0 reaches CASP16 0.2901 versus scratch 0.1330, but RNA3D38 0.2776 versus scratch 0.2723 remains unresolved. The matched CASP16 G0/full head contrast favors Resid-ualGNN83k over MLP83k (0.2901 versus 0.1928), without a corresponding resolved RNA3D38 advantage. Ranking transfer therefore depends on the read-out and the test set. All conditions, validation results and uncertainty analyses are supplementary.

S0, run under the same protocol, reaches meantarget Spearman 0.3639 on RNA3D38 and 0.2025 on CASP16 with the MLP readout and LoRA. With MLP83k and LoRA it reaches 0.3903 and 0.0772, so a strong within-RNA3D point estimate need not carry over to CASP16. All nine S0 head and mode conditions are reported with the other conditions in the supplement.

### Pretrained models recover holo-defined sites from apo structures

Finally, we applied the ligand-binding models with-out refitting to six apo/holo pairs from the SMART-Pocket evaluation [32], asking whether sites defined on the holo structures are recovered from the apo structures. These models recover holo-defined sites from apo structures better when initialized from BRIDGE: G0 reaches apo pooled AP 0.3993 versus 0.2533 for scratch (Supplementary Table S19), compared with 0.6924 versus 0.4657 on the holo structures themselves. Four of the six holo structures (3SUX, 2YIE, 4LVX, 2L8H) were in the ligand-binding training set, so for these targets the apo test asks whether a site seen in one conformation is recognized in another. The gain appears in both exposure tiers: target-averaged apo AP is 0.6781 versus 0.4136 for the four training-seen targets and 0.3201 versus 0.2339 for the two without a direct training match (3D0X, 6XJZ). With only two targets, the second tier is descriptive. Both states for S0, G0, G1 and Scratch-N are reported in Supplementary Table S64.

Per-target effects are heterogeneous: G0 is lower than scratch for both 3D0X/3D0U states, for apo 6WJR and for holo 2L8H, while 3SUY/3SUX has large gains. Leave-one-target-out pooled effects range from 0.1237 to 0.1849 for apo and from 0.1692 to 0.3699 for holo. These ranges show how much single targets influence the pooled effect, but they are not independent replications. Macro-AP sign-flip sensitivities remain unresolved. Per-target scores, exposure tiers and uncertainty conventions are given in the supplement.

Among the eight three-fit ensembles in Supplementary Table S19, S0+*X* has the highest pooled AP in both states (apo 0.4337, holo 0.7422), consistent with task-dependent auxiliary benefits. Among the released encoders (Supplementary Table S19), NA-MPNN exceeds G0 on apo pooled AP (0.4120 versus 0.3993), whereas G0 is higher on apo target-macro AP and both holo endpoints. These descriptive ensemble results do not show that G0 loses less from holo to apo. Five of the six targets in each state have PDB matches in the gRNAde pretraining data, and three in the NA-MPNN pretraining data, and shared TR60 exposure further precludes unseen-target interpretation. Small GNN heads give the same apo-focused picture: G1 full width32/64 gives apo AP 0.4130/0.4181 and holo reference AP 0.7147/0.7610, versus scratch-full apo AP 0.2239/0.2026. All small-head modes are retained in Supplementary Table S69, with complete ensemble metrics in Supplementary Data D100–D104. Released-encoder seed outcomes and per-target AP for the original readout are in D079–D081.

Table 6 compares all nine models using equal-target macro balanced accuracy (BA), validation-threshold MCC and AUROC on the same six target identities, residue mappings and 4 Å labels. The released SMARTPocket checkpoint, averaged over five seeded stochastic passes, reaches apo/holo BA of 0.8500/0.9452, MCC of 0.5580/0.7686 and AUROC of 0.8790/0.9732, the highest point estimate in each column. Among the three-fit ensembles, S0+PairGeom has the highest BA in both states (0.7718/0.8901), whereas the leading MCC and AUROC configurations depend on the state. BA and MCC use each model’s unchanged TR60 validation-MCC-selected threshold; these cutoffs were not optimized for BA. Fixed-threshold and per-target oracle results remain in Supplementary Table S24 and illustrate operating-point sensitivity without establishing probability calibration. SMARTPocket uses full RNA heavy atoms and one released checkpoint, whereas BRIDGE uses its backbone representation and three fitted models. Its configured validation split also contains 2YIE, and released-checkpoint exposure is not proven absent; this is a descriptive local comparison rather than an exposure-clean ranking. Additional pooled results on the retrospectively screened SMARTFlex49 cohort, with label and exposure details, are reported in Supplementary Section 4.4 and Table S25.

**Table 6.** Apo/holo inference without refitting: balanced accuracy, MCC and AUROC across six paired targets.

|  | Apo |  |  | Holo |  |  |
|---|---|---|---|---|---|---|
| Model | BA val | MCC val | AUROC | BA val | MCC val | AUROC |
| S0 | 0.7108 | 0.3683 | 0.8369 | 0.8840 | <u>0.6627</u> | 0.9398 |
| S0+PairGeom | <u>0.7718</u> | 0.4162 | 0.8292 | <u>0.8901</u> | 0.6227 | 0.9327 |
| S0+χ | 0.7142 | 0.3889 | 0.8430 | 0.8543 | 0.6423 | <u>0.9480</u> |
| G0 | 0.7299 | 0.3735 | <u>0.8559</u> | 0.8749 | 0.6254 | 0.9429 |
| G1 | 0.7546 | <u>0.4408</u> | 0.8310 | 0.8570 | 0.6059 | 0.9379 |
| Scratch-N | 0.6569 | 0.2513 | 0.7645 | 0.7543 | 0.4530 | 0.8858 |
| gRNAde | 0.6674 | 0.2425 | 0.8053 | 0.8046 | 0.5064 | 0.9049 |
| NA-MPNN | 0.7396 | 0.4010 | 0.7952 | 0.8419 | 0.5780 | 0.9081 |
| SMARTPocket local | **0.8500** | **0.5580** | **0.8790** | **0.9452** | **0.7686** | **0.9732** |

Every metric is an equal-weight mean over six targets per state. BA, balanced accuracy, is the mean of sensitivity and specificity within each target; MCC, Matthews correlation coefficient; AUROC, area under the receiver operating characteristic curve. BA val and MCC val use the same frozen TR60 validation-MCC-selected cutoff for each model; AUROC is threshold-independent. Models reuse full-adaptation TR60 checkpoints and three-fit probability means; SMARTPocket local uses one released checkpoint and five stochastic passes (cutoff 0.28). Residue mappings, 4 Å labels and masks are shared (605 apo/607 holo labelled residues; 64 positives each). Four holo targets occur in BRIDGE downstream training, and SMARTPocket's configured validation set contains 2YIE. Model inputs and checkpoint exposure differ. Bold/underline mark the best/second-best distinct unrounded value per column, descriptively. AP and additional threshold endpoints remain in Supplementary Tables S19 and S24.

## Discussion

Representations learned from phosphorus and ribose coordinates supported prediction beyond sequence reconstruction even though observed base atoms were withheld. A plausible explanation is that the back-bone retains geometric correlates of molecular recognition. Inter-strand spacing, backbone curvature and sugar orientations describe aspects of groove shape and the organization of loops, bulges and junctions. These features can distinguish exposed surfaces from compact environments and indicate where a binding pocket may form. Structural studies of TAR recognition and experimentally validated geometry-based RNA pocket detection support a connection between RNA conformation and ligand recognition [5, 6]. We therefore hypothesize that inverse-folding-based pre-training makes such recognition-relevant geometry accessible to downstream predictors. This interpretation does not require the backbone to specify every base-mediated contact: it can identify a favorable structural environment without fully resolving the chemical interactions that determine binding specificity.

Excluding observed base atoms is important because it separates the structural detail used for supervision from that required at application time. BRIDGE can learn from sequence, accessibility and base-geometry targets derived from detailed pretraining structures, while its transferred encoder requires only the selected phosphorus and ribose coordinates. This offers a potential route to analyzing backbone-only design intermediates and structural models in which base placement is uncertain. Such uncertainty is relevant to cryo-EM RNA modeling, where flexible regions can be misrepresented by a single atomic model [37]. These applications require sufficiently reliable coordinates for the six input atoms. BRIDGE does not operate directly on density maps or arbitrary coarse-grained traces. Our decoy-ranking experiments assess structure quality and do not test binding-site prediction on predicted coordinates, so compatibility with predicted structures does not yet demonstrate robustness.

The frozen comparisons are consistent with the hypothesis that pretraining makes recognition-relevant geometry accessible. With frozen encoders and matched readouts, G0 improves external ligand-binding AP from 0.5033 for the random-frozen control to 0.6274, an absolute gain of 0.1241. Protein-interface AP similarly increases from 0.5160 to 0.6471, a gain of 0.1312. These improvements indicate that the pretrained representation contributes information that the readout cannot extract as effectively from random features. During full adaptation, both the encoder and the head can learn task-relevant geometry, potentially narrowing the advantage of pretrained initialization. Within RNAProt, where both panels fix pretraining seed 0 and vary the same three down-stream seeds, the pretrained–control gap is smaller under full adaptation: G0 achieves AP 0.6415 versus 0.6225 for scratch, a difference of 0.0190. Pretraining may therefore provide a useful starting representation whose advantage becomes less pronounced as supervised learning reshapes the encoder. Optimization sensitivity offers an alternative explanation to test through controlled optimization and representation-change analyses. For ligand-binding prediction, the frozen and full panels use different pretraining-seed designs, so they do not isolate this narrowing effect.

The mismatch between sequence recovery and downstream performance suggests that reconstruction and functional recognition draw on overlapping but nonidentical structural information. Sequence reconstruction rewards compatibility between nucleotide identity and its structural environment, whereas recognition tasks may weight accessibility and the spatial arrangement of residues more heavily, so improving one objective need not improve the other. Recovery and angular accuracy are therefore useful pretraining diagnostics but not sufficient criteria for selecting a transferable encoder.

The auxiliary-objective results fit this interpretation. The matched sequence-only control transfers without SASA targets, reaching external and protein-interface AP 0.6008 and 0.6329 against random-frozen 0.5033 and 0.5160, so the transfer signal does not depend on base-dependent accessibility supervision. Adding SASA (S0) improves external macro AP, whereas the pooled difference remains unresolved. S0 has no explicit *X* or PairGeom loss and still transfers effectively, while adding or removing the geometric objectives produces non-additive patterns across tasks and adaptation modes at the tested loss weights, and on the apo/holo panel S0+*X* has the highest pooled AP among the eight three-fit ensembles. In the matched-duration frozen comparison, S0 and G0 have similar external pooled AP, while TR60 test macro AP favors G0. The value of an auxiliary objective therefore cannot be summarized by one pooled score, and a loss-weight-scale sweep is needed to test how broadly these patterns hold. Auxiliary supervision appears to change which information the representation emphasizes rather than add a uniform increment in downstream performance.

These results extend protein precedents for transferring inverse-folding representations to molecular recognition [10, 11]. Compared with large-scale protein pretraining using predicted structures, the RNA setting examined here combines a much smaller experimental structural corpus with conformational heterogeneity that motivates evaluation across alternative structures [8, 17]. Within this setting, under full adaptation, G0 achieves a protein-interface AP only 0.0102 below NA-MPNN while requiring coordinates for six observed backbone atoms per nucleotide in-stead of twelve. However, gRNAde’s stronger frozen ligand-binding result shows that relative performance depends on the task and adaptation regime. For ATOM3D structure ranking, G0 with LoRA and the MLP readout reaches mean-target Spearman correlation 0.5107, compared with 0.3739 for the released ARES scores [36]. The absolute difference is 0.1368 across the same four test puzzles. LoRA also raises G0 from its frozen value of 0.3107 by 0.2000, although this four-target difference is not resolved by exact sign-flip tests. The width32 GNN comparison further illustrates this dependence: G1/LoRA reaches 0.4729 versus NA-MPNN/LoRA 0.3523, while gR-NAde/full gives a negative correlation. The strong random-frozen GNN control also prevents attributing every ranking result to pretraining. Differences in inputs and training histories make the cross-model comparisons contextual rather than controlled tests of encoder architecture.

The SMARTPocket comparison shows that useful binding-site information can transfer to un-bound conformations, while threshold selection remains consequential. On the local six-target evaluation, SMARTPocket exceeds BRIDGE in target-macro BA, validation-threshold MCC and AUROC for apo and holo structures. Its full-atom inputs, checkpoint history and validation exposure differ from BRIDGE, limiting attribution of this advantage to the model architecture alone. Known apo/holo PDB and exact-sequence matches were excluded from BRIDGE encoder pretraining. However, four holo targets were present in downstream TR60 training, so their apo counterparts assess cross-conformation transfer rather than generalization to entirely unseen RNA targets.

Future work should test these explanations directly. Probing groove dimensions, backbone curvature and loop/bulge organization, followed by controlled per-turbations of local and nonlocal geometry, would help identify which structural properties support transfer. Matched label-budget experiments, controlled optimization comparisons and measurements of representation change during adaptation could test whether the narrowing pretraining advantage reflects supervised acquisition of useful geometry or changes in the preservation of pretrained features. A six-, eight- and twelve-atom comparison is reported in Supplementary section *Exploratory observed-backbone atom ablation*. Building on that comparison, controlled perturbations and evaluations stratified by coordinate uncertainty should test whether the restricted input provides a practical advantage when base placement is incomplete or unreliable.

The main limitations are the small independent-target panels for ATOM3D and apo/holo, the retro-spective construction of RNA3D38, targeted rather than comprehensive homology screening, and down-stream exposure of four holo targets. Model reporting roles and several readout extensions were retro-spective. Full-adaptation gains are conditional on the shared optimization recipe. The validation-selected lower-rate RNAProt sensitivity retains a pretrained advantage, although it does not establish optimally tuned performance across tasks. Sequence-only down-stream comparisons fix one pretraining seed and do not measure pretraining-seed variability in transfer. Robustness to missing or uncertain coordinates remains untested, and released-encoder comparisons differ in inputs and pretraining histories.

## Conclusion

BRIDGE shows that representations learned from phosphorus and ribose atoms through inverse folding can transfer beyond sequence reconstruction. In matched frozen-readout and full-adaptation comparisons, pretraining improves ligand-binding and protein-interface prediction (for protein interfaces, with the GNN readout). Sequence-only pretraining also transfers to both residue-level tasks without SASA or explicit base-geometry supervision, and the additional value of auxiliary targets depends on the task, aggregation and adaptation. Sequence recovery and angular accuracy are useful pretraining diagnostics but not sufficient criteria for selecting a transferable encoder. Structure-ranking and apo/holo bench-marks extend the assessment, but their small target panels and training exposure limit what they can show. Larger, independently curated apo/holo and decoy panels would allow a more precise assessment of transfer to new RNA targets. Because the transferred encoder requires only phosphorus and ribose coordinates, a next step is to test whether this restricted input gives a practical advantage when base placement is incomplete or unreliable. Within this scope, our results support inverse-folding pretraining as a practical route to transferable RNA structural representations. Their downstream utility, like BRIDGE’s ranking against other methods, depends on the task, readout and adaptation strategy.

## Data availability

Detailed numerical tables are supplied as Supplementary Data in CSV and XLSX formats. Data mappings, implementation details, checkpoint identities and score-replay specifications are documented in the supplement and accompanying evidence files. Public code and checkpoint deposition is pending. The current record supports local verification, not independent end-to-end reproduction.

## AI assistance

AI tools assisted code development, evidence organization and manuscript editing. Their outputs are not independent experimental evidence. The authors retain responsibility for validating the analyses, citations and final text. The supplementary disclosure describes these uses.

## Key Points

- BRIDGE is an RNA encoder pretrained by inverse folding on 2,501 RNA-chain records, using the phosphorus atom and five ribose atoms per nucleotide without observed base coordinates.
- BRIDGE features improve ligand-binding and RNA–protein interface prediction over matched unpretrained controls, both when frozen and as an initialization for full adaptation (with the GNN readout for protein interfaces).
- Sequence-only pretraining also transfers without accessibility or base-geometry targets. The added value of the auxiliary targets depends on the task, the score aggregation and the adaptation mode.
- Structure-ranking transfer depends on the read-out and adaptation mode. On the six-pair apo/holo panel, BRIDGE-initialized models improve pooled AP for recovering holo-defined sites from apo structures, although four holo targets were seen during downstream training.
- Higher sequence recovery during pretraining does not always identify the encoder that transfers best.

## Notes

### Competing Interest Statement

The authors have declared no competing interest.

