## Supplementary data for "BRIDGE: Backbone Representation for Inverse-folding-Driven Generalizable RNA Encoding"

**Supplementary Data.** Detailed seed, target and expanded comparison tables accompany this document as CSV files and an indexed XLSX workbook. Stable identifiers D011, D015, etc. refer to data tables; continued parts share the same identifier. The workbook index gives titles, source-table identifiers and current PDF cross-references. Values retain their reported precision, cohort definitions and seed/ensemble distinctions. Unavailable metrics remain unavailable. The detailed data and concise PDF together constitute the supplementary results.

### 1 Experimental design and reproducibility

**Table S1. Data partitions, exposure controls, and task-specific graph/readout settings.**

| Task | Partition / size | Exposure control and limitations | Graph and readout settings |
|---|---|---|---|
| Source pretraining | Train/validation/test: 2501/82/68 records | Known downstream test matches removed from fitting. Downstream train/validation structures retained. One 3GS5-derived test record shares source-training provenance. | *Encoder:* Directed $C4'$ kNN32 plus both sequence-adjacent directions, with original retained positions. *Readout:* Source causal decoder uses supplied graph, and is discarded at transfer. |
| Ligand binding | Combined HARIBOSS–TR60 train/validation/test: 331/41/43 chains. External: 48 records/44 PDBs | External TL12 retains training-overlap 8F0N. Shared RB9/TE18 chains remain in pooled scores. Exclusion and deduplication sensitivities are reported. | *Encoder:* kNN32 plus index adjacency, with benchmark positions retained, including masked-geometry rows. *Readout:* $C4'$ kNN16 GNN on final scalar states. |
| Protein interface | RNAProt CLEAN train/validation/test: 835/171/191 graphs | Downstream PDB-disjoint split. Test contains 177 PDB-connected units. | *Encoder:* kNN32 plus same-chain residue-number adjacency, with chain offsets 10,000. Inherited admitted nodes are retained. *Readout:* Separate $C4'$ kNN16 GNN/GAT, without sequence or contact labels as features. |
| ATOM3D ranking | Train/validation/test: 12,479/4,000/4,000 decoys | Four test puzzles. Known target aliases and identified Puzzle-21 family relatives excluded from source fitting. | *Encoder:* $C4'$ kNN30, stable distance ordering, no additional adjacency edges, with chain offsets 1000. *Readout:* Mean-pooling MLP/MLP83k, or a separate kNN16 residual GNN. |
| RNA3D / CASP16 | RNA3D train/validation: 329/40 targets. RNA3D38/CASP16 test: 38/20 targets | CASP-overlapping training targets removed. RNA3D38 is a post-hoc sensitivity, not an untouched test. | *Encoder:* $C4'$ kNN30, stable distance ordering, no additional adjacency edges, with sequential admitted positions. *Readout:* Original MLP, exploratory MLP83k and kNN16 ResidualGNN83k extension. |
| Apo/holo | Six paired targets, seven graphs per state. No additional fitting | No known direct retained-source PDB/exact-sequence matches. Four holo targets occur in downstream ligand-binding training. | *Encoder:* kNN32 plus index adjacency, multi-strand 2YIE uses offset 1000. *Readout:* Same TR60-fitted kNN16 GNN, without refitting. |

Counts refer to the datasets as used. PDB and exact-sequence screening is targeted and does not amount to comprehensive homology clearance. Graph settings describe BRIDGE, not the native internals of the released public encoders. Encoder and readout graphs are distinct; see Section 1.5 for edge-direction and node-admission conventions.

Pretraining uses 2,501/82/68 training/validation/test records and fixed epoch-100 checkpoints. CoBRA full/LoRA and the corresponding conformation evaluations pair three pretraining and downstream seeds. Frozen binding-site readouts, RNAProt and RSR fix encoder seed 0 and vary three downstream seeds. Classification metrics score the mean probabilities; ranking metrics average seed-specific correlations. Comparisons across these panels therefore differ in both adaptation and seed design.

S0 retains sequence and SASA supervision, G0 uses the complete objective, and G1 changes decoder context. Model reporting roles were assigned retrospectively. Test-set maxima, including RNAProt G0–LoRA–GAT AP 0.6574 and RSR G1–LoRA–MLP mean-target Spearman 0.5377, are descriptive and do not represent prospective model selection.

#### 1.1 Input, objectives and masks

**Table notation.** Bold and underlined values indicate the best and second-best distinct estimates within the comparison blocks defined by each caption. Higher values are preferred for recovery, angular accuracy, AP, AUROC, MCC and correlation; lower values are preferred for MAE, RMSE and top-1 regret. Individual seeds, SDs, contrasts and confidence intervals are not ranked. Ranking and tie conventions are table-specific and descriptive; they do not imply matched inputs, training exposure or statistical significance.

The feature interface comprises 284 scalar/16 vector node channels and 108 scalar/one vector edge channels. A 13-slot tensor does not mean 13 observed atoms: P,C1′,C2′,C3′,C4′,O4′ and deterministic virtual N are active. Observed N1/N9 and their masks are erased before encoding. Invalid coordinates and degenerate frames are masked rather than imputed. NA-MPNN construction follows commit 9fabc2482092b725e067969fba21297a806b6fda. It uses unnormalized vectors in Å.

The four-layer width 128 GVP encoder has dropout 0.1. The two-layer source decoder uses mean messages, SiLU, residual connections and LayerNorm. Its 29 context channels encode four nucleotide probabilities, 24 circular-bin weights and one availability indicator. G0 zeros preceding-angle channels. G1 uses them causally. Shapes and initialization are unchanged across loss-removal cells. Native identity chooses the source training branch only. Test diagnostics marginalize branches using predicted sequence probabilities.

PairGeom predicts minimum base-heavy-atom distance at boundaries 4, 4.5, 6, 8, 12, 20Å (seven bins) and C1′–glyN–glyN–C1′ torsion in 24 bins. It uses existing undirected graph pairs separated by at least four sequence positions. Torsion targets require distance below 8Å and valid defining atoms. Targets do not add graph edges or encoder inputs. Local $\chi$ targets interpolate neighboring bin centers. Pair-torsion targets use hard bins. Absolute SASA is source-cache supervision with training-only normalization. Upstream ligand occlusion is not independently verified. Relative SASA is not used.

Sequence CE uses label smoothing 0.05 during training. SASA and $\chi$ weights are 0.1 each. Pair-distance/pair-torsion weights are 0.025 each. S0 keeps sequence CE and base-dependent SASA, not sequence CE alone. The original five-variant panel does not separate sequence reconstruction from SASA; the additional matched sequence-only (SASA-off) control in Section 16.1 addresses this distinction. Removing losses does not preserve total gradient magnitude. The four-cell objective design allows an outcome-scale interaction, but no joint interaction interval or biological synergy claim is made.

#### 1.2 Model capacity and computational cost

Parameter counts separate transferred encoders from source-only modules. Counts were obtained from the instantiated BRIDGE and NA-MPNN modules and the released gRNAde parameter tensors.

**Table S2. Parameter counts of transferred encoders and source-only modules.**

| Model | Encoder | Source-only modules | Total source model |
|---|---|---|---|
| BRIDGE G0/G1 | 1,226,692 | 306,820 | 1,533,512 |
| gRNAde | 1,080,470 | 1,067,544 | 2,148,014 |
| NA-MPNN | 1,592,240 | 701,217 | 2,293,457 |

The comparisons therefore differ in capacity as well as observed atom inputs, pretraining data and training history. The counts refer to the original four-layer BRIDGE model reported here.

Six G0/G1 pretraining fits required 640.914–655.655 s each (3,891.473 s in total), using FP32 on one RTX 5070 Ti GPU per fit. These timings exclude preprocessing, evaluation and downstream fitting. Exact historical PyTorch/CUDA versions were not recorded, and no harmonized cross-model runtime or memory benchmark was performed.

#### 1.3 Encoder and training implementation

**Source admission and topology.** Sequences are uppercased with T mapped to U. A source conformer is selected lexicographically by the number of canonical residues with all five sugar-ring atoms, then the number with C4′, then the earliest conformer index. Canonical residues with observed C4′ are retained with their original position indices. Native glycosidic N is not a row-admission requirement. In the archived source-coordinate convention, presence requires finite coordinates with norm greater than $10^{-3}$Å to reject rotated missing-atom sentinels. This origin-dependent sentinel rule is not a generic missing-atom detector for arbitrary coordinates. Records exceeding 500 admitted residues are excluded before graph construction, not randomly cropped. The subsequent retained-source membership fixes the 2501/82/68 records used here.

Each source node receives directed edges from its nearest min(32, $n-1$) C4′ neighbors. Both directions of sequence-position adjacency ($|i-j| = 1$) are added and duplicate edges removed. A singleton receives a self edge. There is no additional radius cutoff. Sequence adjacency is an index-based flag, not a fresh chemical-bond validation. This encoder graph is distinct from the downstream C4′-kNN16 readout graph. Source batches pack whole records up to 1500 nodes, without inter-record edges. Every retained training record is visited once per epoch.

**Sugar frame and feature channels.** The local frame uses the least-variance singular vector of the centered five-atom ribose ring as its normal $z$, with sign chosen by $(C2' - C1') \times (C3' - C2')$. Axis $x$ is the normalized in-plane projection of $C4' - C1'$ and $y = z \times x$. Feature translations use C4′. Validity requires all five ring atoms, projection conditioning greater than $10^{-3}$ and smallest/largest singular-value ratio below 0.5. Invalid frames are zeroed without removing the residue. Virtual N also requires a valid frame and its defining sugar atoms. All observed native-N channels and masks are erased before feature construction.

For $K$ equally spaced centers $c_k$ from 0 to $D$, the radial feature is $\exp[-((d - c_k)/(D/(K - 1)))^2]$. Each of 13 fixed atom slots contributes presence, 16 radial features ($D = 12$Å) and three local unit-direction components relative to C4′. Inactive or missing slots contribute zero. Only P, C1′, C2′, C3′, C4′, O4′ and virtual N are active. Two trace-neighbor blocks each contribute presence, eight radial features ($D = 12$Å) and three local direction components, giving $13(1 + 16 + 3) + 2(1 + 8 + 3) = 284$ node scalars. The 16 node vectors are 13 masked unit offsets and three frame axes.

The 108 edge scalars comprise 16 C4′-distance radial features ($D = 20$Å), $3 \times 3 \times 8 = 72$ anchor-pair radial features for P/C4′/virtual N ($D = 20$Å), six endpoint-local displacement components, nine relative-rotation entries, three sequence-separation features and two adjacency flags. The separation features are $\text{clip}(ps - pd, -32, 32)/32$, $\sin((ps - pd)\pi/16)$ and $\cos((ps - pd)\pi/16)$ for stored source/destination positions $ps, pd$. The edge vector is the unit C4′ displacement. Unit normalization clamps norms at $10^{-8}$. Invalid frames zero local-direction/rotation channels. Valid radial and presence channels remain available.

**Encoder, source decoder and transfer.** Node scalar/vector widths project to 128/16 and edge scalar/vector widths to 128/4. Four GVP layers use dropout 0.1. The transferred embedding is the final LayerNorm-normalized 128-dimensional scalar state. The two-layer causal decoder projects the 108 edge scalars to width 128, uses mean message aggregation, residual/dropout/LayerNorm updates and a 128–256–128 SiLU feedforward block. Each record receives a random decoding permutation. Its 29 context channels comprise four sequence channels, 24 circular-angle channels and one angle-validity flag. G0 zeros the latter 25. At every layer, self/future messages use the original encoder state and zero target context, preventing multi-layer target leakage. Training is teacher-forced. Diagnostic rollout uses predicted context. Sequence and local-$\chi$ outputs read decoder states. SASA and pair heads read encoder states. Pair heads are linear maps of $[hi + hj, |hi - hj|]$. Only encoder features/states, not decoder states or auxiliary heads, transfer.

**Masks, reductions and training-only normalization.** Sequence CE averages over all admitted residues in a batch with label smoothing 0.05. For the local targets, bond ok requires all checkable intra-residue bonds to have lengths in $[0.9, 2.0]$Å. Missing endpoints are skipped. This target mask does not delete encoder nodes or sequence labels. SASA loss is masked mean squared error against the training-standardized absolute SASA target, using mean 185.36969 and sample SD 57.69235 Å$^2$. Local $\chi$ additionally requires its defining atoms. Its target interpolates two adjacent circular bin centers. The native-nucleotide branch's soft-target CE is averaged over valid residues and divided by the retained-training support-weighted conditional prior entropy, 1.76840536 nats.

PairGeom uses canonical undirected pairs already in the source graph, with sequence separation at least four, bond ok and at least three observed base-heavy atoms at each endpoint. Minimum base distance supplies a hard seven-class target at boundaries 4, 4.5, 6, 8, 12 and 20Å (a boundary belongs to the lower bin). Torsion additionally requires distance below 8Å and observed C1′/native glycosidic N at both endpoints. Unlike local $\chi$, pair torsion uses hard 24-bin targets, not adjacent-bin interpolation. Distance and valid-torsion CE are separately averaged across eligible batch pairs and divided by their retained-training marginal entropies, 1.43441993 and 2.76132569 nats. Empty masked supports contribute zero. These normalized losses are the terms weighted by 0.1 and 0.025 in the main objective. Loss removals do not renormalize the remaining terms. Training logs aggregate batch summaries with residue weights and are not substitutes for exact whole-dataset valid-target averages.

**LoRA placement.** LoRA wraps every linear module inside the encoder, including scalar and vector projections, not just an attention block. For rank 4 and alpha 8, $W\text{eff} = W + 2BA$. $A$ is initialized from a normal distribution with SD 0.01 and $B$ to zero. Original encoder parameters (including biases and LayerNorm parameters) are frozen, while adapters and the separately initialized task head are trained. The effective-weight interface also covers vector projections that access a linear module's weight directly. Feature construction is unchanged and source-only decoder/auxiliary heads are not adapted.

**Controls.** S0 uses the preceding-sequence decoder context of G0 and retains the same modules while disabling the angle and pair-geometry losses. Random-frozen uses an unpretrained encoder with a trained readout and no encoder updates; Scratch-N/full trains both encoder and readout.

#### 1.4 Atom slots, geometric conventions and head dimensions

The fixed slot order is P, C4′, glyN, O5′, C5′, C3′, O3′, O4′, C1′, C2′, O2′, OP1, OP2. Here glyN denotes the glycosidic attachment nitrogen (N9 for A/G, N1 for C/U) in the raw storage schema. Its observed coordinates and presence mask are erased. The active glyN slot is filled only by a deterministic virtual point. With coordinates in Å,

$$b = C1' - O4', \quad c = C2' - C1', \quad a = b \times c,$$

$$\widetilde{N} = C1' - 0.56967352a + 0.51055973b - 0.53122153c.$$

The implementation follows NA-MPNN [6, 1]. The coefficients apply to this fixed coordinate-unit convention, not arbitrary rescaling. There is no native-base-dependent coefficient choice. Virtual-point validity requires the three defining atoms and the valid five-atom sugar frame specified above. Rigid transformations preserve this construction. The use of an unnormalized cross product does not imply invariance to bond-length scaling.

For an edge with stored source $s$ and destination $d$, the displacement is $u = (C4'_d - C4'_s)/\max(\|C4'_d - C4'_s\|, 10^{-8})$. Endpoint-local directions are $R_s^T u$ and $-R_d^T u$, relative rotation is $R_d^T R_s$, and separation is $p_s - p_d$, where $p$ is the stored position index. Previous/following trace blocks use position differences $-1/+1$ among existing incoming edges. They are not additional chemical-bond tests. Their displacement is neighbor C4′ minus the current C4′, with local unit directions. Missing trace neighbors contribute zero. Chain offsets prevent false sequence-adjacency flags across represented chains. Spatial edges can still connect chains within the same record.

The local $\chi$ quartets are O4′–C1′–N9–C4 for A/G and O4′–C1′–N1–C2 for C/U. Let $K = 24$, $w = 2\pi/K$, and $c_k = -\pi + (k + 1/2)w$. Bin intervals are $[-\pi + kw, -\pi + (k+1)w)$ with circular wrapping. Training interpolates centers, not hard intervals: for $q = ((\chi + \pi) \bmod 2\pi)/w - 1/2$, place weights $1 - (q - \lfloor q \rfloor)$ and $q - \lfloor q \rfloor$ on $\lfloor q \rfloor \bmod K$ and its successor. Numerically near-integer coordinates are snapped within $16\epsilon K$. The four identity-conditional branches emit $4 \times 24$ logits. Identity selects the branch only during supervision. Diagnostic rollout marginalizes using predicted sequence probabilities. This source-only sequence/conformation decoder is inspired by LASErMPNN, but does not import its protein weights [3, 8]. Below, $h_i$ denotes the final normalized width-128 encoder scalar state. Sequence label smoothing uses target mass $(1 - 0.05)\mathbf{1}_{k=y_i} + 0.05/4$, distinct from circular interpolation of $\chi$ targets.

**Exact target-bond mask.** All checkable backbone bonds are P–OP1, P–OP2, P–O5′, O5′–C5′, C5′–C4′, C4′–O4′, C4′–C3′, O4′–C1′, C1′–C2′, C2′–O2′, C2′–C3′ and C3′–O3′. Purine bonds add C1′–N9, N9–C8, C8–N7, N7–C5, C5–C6, C6–N1, N1–C2, C2–N3, N3–C4, C4–C5 and C4–N9. Pyrimidines instead add C1′–N1, N1–C2, C2–O2, C2–N3, N3–C4, C4–C5, C5–C6 and C6–N1. Base-specific additions are C6–N6 (A), C6–O6 and C2–N2 (G), C4–N4 (C), and C4–O4 (U). Present endpoints must lie within [0.9, 2.0]Å. Missing endpoints are skipped. This is a supervision mask, not a requirement that every listed atom be present and not an encoder-node deletion rule.

**Table S3. Source prediction heads and downstream readout architectures.**

| Module | Executed dimensions and operation |
|---|---|
| Source heads | Sequence 128 → 4, absolute SASA 128→ 1, local $\chi$ 128 → 96, pair-distance 256 → 7, pair-torsion 256 → 24. All are linear maps, including biases. Pair input is $[h_i + h_j, \lvert h_i - h_j \rvert]$. |
| GNN (82,689) | 128 → 128 projection, two updates each summing separate 128 → 128 self and mean-neighbor maps, ReLU/dropout 0.3, 128 → 1 output. |
| GAT (82,561) | 128 → 172 projection, two additive-attention layers with four 43-wide heads and residual self path, ELU/dropout 0.3, 172 → 1 output. This is a local GAT-style head, not a named published model. |
| MLP (16,641) | Mean embedding 128 → 128 → 1, ReLU and dropout 0.1. Used for the principal RSR/RNA3D panels. |
| MLP83k (83,031) | 128 → 230 → 230 → 1, ReLU/dropout after hidden layers. Dropout is 0.3 in the earlier frozen-head panel and 0.1 in the RSR adaptation20 panel. |
| Residual GNN (83,201) | GNN dimensions above with two width-128 LayerNorms and residual additions, RSR adaptation20 sets dropout to 0.1Node outputs are mean-pooled to one ranking score. |

#### 1.5 Encoder topology is not readout topology

The feature builder consumes supplied edges. Transferring encoder weights does not force a universal graph constructor. The graph settings in Table S1 describe BRIDGE, not the native internals of the released public encoders. $k$ is capped by $n - 1$, with no inter-record edges.

Graph and readout settings are summarized alongside dataset splits and exposure controls in Table S1.

For source/CoBRA/RNAProt edges, stored source indices are neighbors of the destination. The ranking constructors store each query as source and its neighbors as destinations. Direction-sensitive channels follow the stored source/destination convention. The ranking adapter is therefore not claimed to reproduce the source topology exactly. RSR requires the five sugar atoms, allows terminal missing P and rejects duplicate active atoms. RNAProt inherits its cache admission. Merely masking native N channels does not undo that admission. Cross-task score differences do not isolate pretraining from adapter/topology differences.

#### 1.6 Optimization and seed design

Primary BRIDGE and released-encoder downstream fits use FP32 Adam with weight decay $10^{-4}$. Full adaptation uses encoder/head rates $10^{-4}/10^{-3}$; LoRA uses rank 4, alpha 8 and adapter/head rates $10^{-3}/10^{-3}$. Frozen fits update only the readout. Cross-mode contrasts therefore combine update parameterization and encoder learning rate; same-mode objective-removal contrasts keep these rates fixed. Task-specific budgets, seed axes and selection are given in Table S4 and Section 1.10. Native RNAProt baselines and learning-rate sensitivities retain their separately stated recipes.

**Table S4. Training budgets, seed axes and checkpoint selection.**

| Panel | Source / downstream seeds | Epochs | Selection and reported aggregation |
|---|---|---|---|
| Pretraining | Three independent seeds 0,1,2 | 100 | Fixed epoch 100, mean of three diagnostic metrics. |
| CoBRA frozen S0/G0/G1/random | Source 0 fixed, three head seeds | 120 | Validation-selected, three-probability ensemble. |
| CoBRA adapted / conformation full | Paired source/downstream $s = 0, 1, 2$, fresh head seed $s$ | 120 | Earliest maximum validation pooled AP every five epochs, three-probability ensemble. Conformation applies the same CoBRA weights and cuts. |
| RNAProt main head panel | Source 0 fixed, head 91000+$s$, training RNG 92000 $+s$ | 40 | Earliest maximum validation pooled AP every five epochs, probability ensemble. Scratch initialization also fixed at 0. |
| RNAProt older loss panel | Paired source/downstream $s$, fresh head reseeds global RNG to $s$ | 40 | Same AP cadence/earliest rule, but distinct RNG/source-seed contract, probability ensemble. Scratch initializes with $s$. |
| RSR adaptation | Source 0 fixed, head 91000+$s$, training RNG 92000 $+s$ | 20 | Earliest finite validation release-target macro Spearman, mean of three seed-specific correlations. |
| RSR loss-removal probe | Three source seeds, readout seed 0 | 100 | Frozen embedding regression, batch 256, distinct from adaptation20. |
| RNA3D / CASP | Source 0 fixed, three downstream seeds | 20 | Earliest finite validation target-macro Spearman, same selected model on both cohorts, seed-metric mean. |
| Released IF encoders | One fixed released checkpoint, three downstream seeds | 120/40/20 | CoBRA/RNAProt/ranking budgets respectively, same endpoint and aggregation as each matched task panel, but different inputs and pretraining inventories. |

The main RNAProt head/adaptation panel fixes pretraining seed 0, whereas the objective-removal panel pairs three pretraining and downstream seeds. Their G0/G1/Scratch full-GNN ensemble AP values are 0.6415/0.6506/0.6225 and 0.6272/0.6305/0.5970, respectively. These panels cannot isolate a head effect by direct subtraction.

#### 1.7 Pretraining exposure controls

The source screen retains 2,501/82/68 records from an initial 2,534/84/79. It removes known downstream test/external PDB and exact-sequence matches, propagates exclusions through shared source identities and provenance PDBs, and excludes identified Puzzle-21 relatives. Downstream training/validation structures were retained; their classification and ranking labels were not used in pretraining. Validation-structure exposure nevertheless limits a strictly inductive interpretation.

This screen uses exact identity and provenance closure, rather than a new alignment-based homology search. Of 2,501 retained training records, 1,547 lack significant chain-level Rfam assignments; these unknown assignments do not establish family independence.

#### 1.8 SASA targets

BRIDGE imports absolute per-residue SASA from sasa list in the processed RNASolo/gRNAde source. The same conformer index and retained residue positions select coordinates and SASA. These targets are not recomputed from the six observed encoder atoms and are not divided by nucleotide-specific maximum-accessibility constants. Thus they are absolute SASA, not RSA. Base atoms can contribute to the target surface and to occlusion. Excluding their coordinates from encoder inputs does not make the target base-independent.

The upstream preprocessing calls `biotite.structure.sasa` and sums atom areas by residue. Inspection of Biotite 1.6.0 gives Shrake–Rupley defaults of a 1.4 Å probe, 1,000 Fibonacci sphere points and ProtOr radii, excluding solvent, monoatomic ions and hydrogens. Other loaded atoms can occlude RNA atoms. The serialized targets do not record the original Biotite version or atom/assembly context, so these inspected defaults cannot be assumed to describe every historical target.

Only retained-training residues passing bond ok determine the normalization: mean 185.36969 Å$^2$ and sample SD 57.69235 Å$^2$. The linear SASA head reads encoder states. Its standardized target is $z_i = (s_i - \mu_{\text{train}})/\sigma_{\text{train}}$. With prediction $\hat{z}_i$, the term is the mean of $(\hat{z}_i - z_i)^2$ over valid residues, weighted by 0.1 in the total objective. Validation/test statistics do not set this normalization. This source target is distinct from any separately computed downstream hand-crafted SASA feature.

#### 1.9 Datasets and residue labels

The retained source removes known downstream TEST/external provenance-PDB and normalized exact-sequence matches with source-cluster closure, while retaining downstream training/validation. Identified Puzzle 21 Rfam relatives 5NZ3, 5O62, 5O69 were also excluded. This is a targeted retrospective screen, not a complete alignment-based homology audit. Source-test still shares 3GS5 provenance with training and is diagnostic, not an independent generalization endpoint.

Ligand-binding labels and splits are inherited from the HARIBOSS–TR60 sequence/contact arrays [4], with 331/41/43 admitted training/validation/test chains. Sequences are uppercased with T mapped to U; native indices are A=0, G=1, C=2 and U=3. Records are keyed by dataset and chain. Missing geometry is masked while admitted label positions are retained. Upstream ligand chemistry and modified-residue annotation were not independently validated for every training record.

RB9/TE18 share 1NEM A, 1Q8N A and 2TOB A (23, 38 and 20 residues). TL12 has two chains from 8HB1. An exact comparison of the admitted record bank confirms that each shared RB9/TE18 chain has identical labels, nucleotide identities, positions, coordinates, masks and edges. Both set-qualified copies remain in the external 48 pool (2369 residues/44 PDBs). PDB resampling groups them but does not remove their duplicated contribution. Excluding 8F0N yields 47 records/43 PDBs. This removes a known downstream-training overlap, not the duplicates or all sequence/family exposure. The original result is retained. An explicit deduplicated sensitivity is now reported in Section 9.

RNAProt uses CLEAN=1, with 835/171/191 graphs and 177 PDB-connected test units. Inherited residue admission can depend on atom availability, even though base geometry is not an encoder channel. No comprehensive family-independent or language-model-exposure clearance is asserted. RSR native targets 18–21 and identified aliases/family relatives were screened from source fitting. Four test puzzles still provide only four independent units.

##### 1.9.1 Direct identity overlap

Each registered downstream identity was compared with all retained source-training provenance PDBs and normalized cached or pre-crop-coordinate sequences. No direct PDB or exact-sequence match remained. The table counts registry entries, including multiple views of the same conformation targets, rather than independent evaluation samples. Coordinate-fragment sequences are not necessarily complete biological sequences.

**Table S5. Retained-pretraining matches to downstream identity registries.**

| Registered identity view | Records | Source-train PDB hits | Exact seq. hits |
|---|---|---|---|
| CoBRA-current | 91 | 0 | 0 |
| CoBRA-raw | 49 | 0 | 0 |
| HARI | 66 | 0 | 0 |
| RNAProt | 191 | 0 | 0 |
| RSR | 4 | 0 | 0 |
| RSR-known-relatives | 1 | 0 | 0 |
| SHAMAN-derived | 81 | 0 | 0 |
| SMARTPocket | 15 | 0 | 0 |
| SMARTPocket-combined | 14 | 0 | 0 |
| SMARTPocket-historical | 12 | 0 | 0 |

All 14 conformation graph sequences also lacked direct retained-pretraining matches. These checks do not establish alignment-based homology or family independence.

##### 1.9.2 Pretraining and downstream conformation exposure

**Table S6. Pretraining and downstream-training exposure of conformation pairs.**

| Apo/holo pair | Source PDB/seq. hit | TR60 train PDB hit | TR60 train seq. hit |
|---|---|---|---|
| 1ANR/2L8H | No | Yes | Yes |
| 3D0X/3D0U | No | No | No |
| 3SUY/3SUX | No | Yes | No |
| 6WJR/2YIE | No | Yes | Yes |
| 6XJZ/6XJQ | No | No | No |
| 7KD1/4LVX | No | Yes | Yes |

Source columns describe pretraining exposure; TR60 columns describe downstream-training exposure of either member of each pair. Four holo identities occur in TR60 training, so their apo partners test recognition across conformations of an RNA seen during downstream fitting. Identity matches do not establish identical coordinates or labels. The combined 2YIE record contains chains X/Z. The remaining two pairs have no direct audited match but have not undergone comprehensive homology screening. Scratch and pretrained models share downstream exposure, whose effect may differ between initializations.

**Table S7. Training-chain identities in the conformation exposure tiers.**

| Apo/holo pair | Downstream exposure tier | TR60 training PDB/chain |
|---|---|---|
| 1ANR/2L8H | Holo-seen | 2L8H_A |
| 3D0X/3D0U | No direct match | – |
| 3SUY/3SUX | Holo-seen | 3SUX_X |
| 6WJR/2YIE | Holo-seen | 2YIE_X, 2YIE_Z |
| 6XJZ/6XJQ | No direct match | – |
| 7KD1/4LVX | Holo-seen | 4LVX_A |

#### 1.10 Optimization, seeds and selection

Source pretraining uses FP32 AdamW for 100 epochs, learning rate 0.0002, weight decay 0.0001, gradient clipping at 1, a 500-node sample cap and a 1,500-node batch cap. CoBRA uses the downstream rates in Section 1.6, 120 epochs, one graph per update and positive-class weight 2.239531. Validation AP every five epochs selects the earliest maximum; scratch uses the same budget and selection. Validation preserves training RNG state and modes.

RNAProt follows Section 1.6 for 40 epochs, with one graph per update and training-derived negative/positive BCE weighting, without gradient clipping or embedding normalization. Pretrained and random-frozen encoder seed 0 is fixed. Head, training-RNG and shuffle seeds are $91000 + s$, $92000 + s$ and $100003s + \text{epoch}$. Validation AP every five epochs selects the earliest maximum.

Equal-budget controls measure initialization effects under a shared optimization recipe, rather than performance against independently optimized scratch models. RNAProt Scratch-GNN seed 2 selected epoch 5 and showed little training-loss reduction (1.0378 at epoch 1; 1.0405 at epoch 40). By contrast, the RSR residual-GNN and RNA3D MLP scratch runs reduced training loss despite selecting epoch 1. A validation-selected learning-rate sensitivity is reported in Sections 9.1 and 16.3.

**Seed reporting.** Binary tables distinguish each seed's metric, its mean and sample SD, and the metric of the probability ensemble. Sample SD uses denominator $3-1$; the ensemble is neither a fourth seed nor the mean seed metric. Source diagnostics average three independently scored pretraining runs. The seed axes specified above determine which source of fitting variability each panel measures.

RSR 20 uses Adam, full encoder/head rates 0.0001/0.001, LoRA rate 0.001, weight decay 0.0001, dropout 0.1, batch 16 and microbatch cap 1500 nodes. RMSD and embeddings use initial training-only normalization. The loss is decoy-weighted standardized-RMSD MSE. MLP heads operate on normalized mean embeddings. The residual graph head uses normalized nodes, fixed C4′-kNN16 topology and scalar mean readout. Validation release-target macro Spearman selects the earliest maximum. Bound/free Puzzle 14 validation releases are separate in the selection contract. A merged-puzzle diagnostic is not substituted. All 24 conditions/72 evaluations are retained.

#### 1.11 Conformation mapping and inference

The six apo/holo pairs are 3SUY/3SUX, 3D0X/3D0U, 6WJR/2YIE, 7KD1/4LVX, 6XJZ/6XJQ and 1ANR/2L8H. The latter includes NMR entries. The first coordinate model and last-listed alternate atom coordinates follow the archived parser. Modified terminal 3SUY/3SUX C101 (CCC) and 2YIE GTP retain canonical base labels. Holo 2YIE is represented as a two-strand record with chain offset 1000. Missing-label observed residues remain in the graph and are excluded only from the metric denominator. There are 607 observed residues per state, 605 labelled apo/607 labelled holo and 64 positives in each.

The same three TR60-fitted Scratch-N checkpoints are applied without refitting. Conformation results do not select checkpoints, thresholds or cohort membership.

#### 1.12 Metric definitions and uncertainty

For a labelled collection $D$ with binary contact/interface labels $y_i \in \{0,1\}$, average the three fitted models' probabilities, $\bar{p}_i = \frac{1}{3}\sum_{s=1}^{3} p_{is}$. Sort distinct scores into decreasing thresholds $t_1 > \cdots > t_L$, including all ties together. With predictions $\bar{p}_i \geq t_\ell$,

$$P_\ell = \frac{\mathrm{TP}_\ell}{\mathrm{TP}_\ell + \mathrm{FP}_\ell}, \qquad R_\ell = \frac{\mathrm{TP}_\ell}{\sum_{i \in D} y_i}. \tag{1}$$

Stepwise average precision, not trapezoidal PR area, is

$$\mathrm{AP}(D) = \sum_{\ell=1}^{L} (R_\ell - R_{\ell-1}) P_\ell, \qquad R_0 = 0. \tag{2}$$

For units $g = 1, \ldots, G$, let $D_g$ contain all their labelled records. We report

$$\mathrm{AP}_{\mathrm{pool}} = \mathrm{AP}\left(\biguplus_g D_g\right), \qquad \mathrm{AP}_{\mathrm{macro}} = \frac{1}{G}\sum_g \mathrm{AP}(D_g). \tag{3}$$

Pooled AP compares residues across units. It is not the residue-weighted mean of within-unit AP. Macro AP ranks within each unit and then weights units equally. CoBRA uses PDBs. SMARTPocket uses paired targets with chains pooled within each state. The multiset union retains set-qualified duplicates. In RNAProt, 55 of 177 PDB-connected units have no positives: full-unit macro AP is undefined, not replaced by a positive-only subset.

#### 1.13 Sequence and angular metric definitions

Source metrics average three independently scored decoding runs, not their predicted probabilities. Let $q_{si}(a)$ be sequence probability for nucleotide $a$ at residue $i$ under source seed $s$, and let $a_i$ be native identity. For $N = 5,568$ residues,

$$\mathrm{Recovery} = \frac{1}{3N}\sum_{s=1}^{3}\sum_{i=1}^{N} \mathbf{1}\left[\arg\max_a q_{si}(a) = a_i\right]. \tag{4}$$

These are encoder-plus-source-decoder diagnostics, not scores of a transferred encoder alone.

For the $N_\chi$ valid angle targets in mask $V$, predictions marginalize nucleotide-specific angle branches using predicted sequence probabilities, $r_{si}(b) = \sum_a q_{si}(a) r_{si}(b \mid a)$, rather than selecting with native identity. Let $\hat{\chi}_{si}$ be the centre of the highest-probability bin and

$$\delta_{si} = \left|\mathrm{wrap}_{[-180^\circ,180^\circ)}(\hat{\chi}_{si} - \chi_i)\right|. \tag{5}$$

For $\tau \in \{15^\circ, 30^\circ\}$, the tolerance accuracy and circular mean absolute error are

$$A_\tau = \frac{1}{3N_\chi}\sum_{s=1}^{3}\sum_{i \in V} \mathbf{1}[\delta_{si} \leq \tau], \qquad \mathrm{MAE}_\chi = \frac{1}{3N_\chi}\sum_{s=1}^{3}\sum_{i \in V} \delta_{si}. \tag{6}$$

The implementation includes a $10^{-9}$ degree tolerance at the comparison boundary. These are angular accuracies, not classification precision $\mathrm{TP}/(\mathrm{TP}+\mathrm{FP})$. Bin accuracy compares the predicted bin with the target's half-open bin among 24 circular bins. Percentages multiply accuracies by 100. MAE is in degrees. The training-only global prior has no fitted test parameters.

The tables below report four angular metrics with conditional component-bootstrap intervals: bin and tolerance accuracies are fractions, and angular MAE is in degrees. Removing $\chi$ or both objectives removes direct $\chi$ supervision. Their untrained-head diagnostics are labelled accordingly. S0+$\chi$ retains trained $\chi$ supervision. No source metric is used to choose a downstream model. Source diagnostics cover 68 records/5568 residues, including one 3GS5-derived record of 36 residues sharing source-training provenance. Removing it leaves 67 records/5532 residues and 5518 valid angles. Main Table 2 reports the 68-record summary; Supplementary Table S9 retains both cohorts and all angular metrics. Angular masks are inherited. Untrained angular outputs in S0+PairGeom cells appear only as explicit diagnostics in the source evidence, not as claimed conformation performance.

For ranking, $\rho_{\text{target}} = \frac{1}{4}\sum_{g=18}^{21} \text{Spearman}(\hat{r}_g, r_g)$ is computed separately for each seed, then averaged. Pooled correlation instead ranks all 4000 decoys together. Neither is a probability ensemble. Seed SD describes variation across three fits. Target-bootstrap CI resamples four puzzles while averaging the three seed metrics within each draw. These are different uncertainty summaries.

Binary metrics include AP, AUROC, Brier score and MCC at fixed 0.5, validation-selected and test-label oracle thresholds. The oracle is nondeployable. Unless stated otherwise, downstream intervals use 10,000 paired draws and source intervals use 20,000 draws. Intervals condition on fitted models, are unadjusted for multiplicity and do not include prior model/cohort selection uncertainty. Undefined macro units are retained as undefined; sample SD is not a confidence interval.

Pooled AP depends on within- and between-structure score ordering, whereas macro AP weights within-PDB AP values equally. For example, the TR60-test G0–scratch differences are 0.0433 [0.0156, 0.0723] for pooled AP and 0.0293 [-0.0027, 0.0609] for macro AP. An interval spanning zero is an unresolved contrast, not an equivalence test.

Top-1 regret for RSR target $g$ and seed $s$ is $r_{g,\arg\min_i \hat{r}_{gsi}} - \min_i r_{gi}$ in Å, where $r$ is reference RMSD and lower is better. Per-target regret rows average this metric across three seeds. They do not select a decoy from averaged predictions. Predicted-score ties follow the first minimum in the archived row order.

Conformation metrics concatenate graphs within each state/target after averaging seed probabilities; target-macro metrics weight targets equally. Leave-one-target-out analyses remove one complete target and recompute pooled AP. They measure influence, not confidence intervals. RSR instead averages seed-specific correlations. Four-puzzle resampling has only 35 unordered count patterns, with unequal multinomial probabilities, limiting population inference.

#### 1.14 Small-panel inferential sensitivity

Exact sign-flip diagnostics enumerate all $2^n$ sign assignments to the full-precision target effects and compare absolute mean differences with tolerance $10^{-12}$. SMARTPocket uses target-macro AP; RSR uses mean-target correlation after averaging seed-specific correlations.

**Table S8. Small-panel bootstrap intervals and exact sign-flip sensitivities.**

| Panel | Contrast | Bootstrap difference [95% CI] | Sign-flip $p$ |
|---|---|---|---|
| SMART apo | S0 / full – Scratch-N | 0.1713 [0.0199, 0.3719] | 0.1250 |
| SMART apo | G0 / full – Scratch-N | 0.2051 [0.0361, 0.3766] | 0.1250 |
| SMART holo | S0 / full – Scratch-N | 0.2126 [0.0139, 0.4229] | 0.1562 |
| SMART holo | G0 / full – Scratch-N | 0.2074 [0.0053, 0.4251] | 0.1875 |
| RSR | G0 LoRA – frozen | 0.2000 [0.0941, 0.3528] | 0.1250 |
| RSR | G1 LoRA – frozen | 0.1907 [0.0444, 0.3715] | 0.2500 |

These tests require sign symmetry or within-target model-label exchangeability under the null and need not be dual to percentile-bootstrap intervals. With four nonzero effects the minimum two-sided $p$ is 0.125; with six it is 0.03125. Unresolved results can therefore reflect limited panel size. These tests address target-macro metrics, not pooled AP.

### 2 Pretraining diagnostics

**Table S9. Pretraining diagnostics on the pretraining test set at epoch 100, averaging metrics over three pretraining seeds.**

| Model | Recovery | $A_{15}$ | $A_{30}$ | Bin acc. | MAE (°) |
|---|---|---|---|---|---|
| **All 68 source-test records** | | | | | |
| Sequence-only | 0.6088 | – | – | – | – |
| S0 | 0.6219 | – | – | – | – |
| S0+PairGeom | <u>0.6245</u> | – | – | – | – |
| S0+$\chi$ | **0.6276** | 0.8718 | **0.9527** | 0.6174 | <u>10.6661</u> |
| G0 | 0.6238 | <u>0.8730</u> | <u>0.9489</u> | <u>0.6260</u> | **10.5779** |
| G1 | 0.6181 | **0.8735** | 0.9486 | **0.6289** | 10.6773 |
| Training prior | – | 0.7285 | 0.8909 | 0.4418 | 15.0389 |
| **Excluding 3GS5: 67 records** | | | | | |
| Sequence-only | 0.6092 | – | – | – | – |
| S0 | 0.6222 | – | – | – | – |
| S0+PairGeom | <u>0.6241</u> | – | – | – | – |
| S0+$\chi$ | **0.6276** | 0.8713 | **0.9526** | 0.6171 | <u>10.6691</u> |
| G0 | 0.6238 | <u>0.8725</u> | <u>0.9488</u> | <u>0.6263</u> | **10.5713** |
| G1 | 0.6179 | **0.8731** | 0.9485 | **0.6288** | 10.6772 |
| Training prior | – | 0.7282 | 0.8913 | 0.4416 | 15.0410 |

All 68 records contain 5,568 nucleotides and 5,554 valid angles. Excluding 3GS5 leaves 67 records, 5,532 nucleotides and 5,518 angles. Training prior is a structure-independent baseline obtained by normalizing global soft counts over 24 circular $\chi$ bins from the retained training set, pooled across nucleotide identities. It predicts the centre of the most probable bin (−157.5°) for every residue, without test-set fitting. Values are fractions except angular MAE (degrees, lower is better). Bold and underline mark the best and second-best distinct values within each record set (all 68 records or the 67 without 3GS5), excluding the prior. Emphasis is descriptive. Dashes indicate untrained or inapplicable outputs. Uncertainty for all 68 records, individual seeds and the separate sensitivity analysis on the common 26-record subset are supplementary. Ties at the displayed precision share emphasis. $A_{15}$ and $A_{30}$ are angular accuracies within 15 and 30 degrees; bin acc. is the 24-bin classification accuracy; MAE is mean absolute circular error.

Angular outputs from models without angle supervision are included only as untrained-head diagnostics.

**Table S10. Confidence intervals for pretraining diagnostics.**

| **Model** | **Metric** | **Estimate [95% CI]** |
|---|---|---|
| S0 | bin_accuracy | 0.0137 [0.0119, 0.0155] |
| S0 | circular_mae_deg | 97.4455 [95.7236, 99.3131] |
| S0 | pooled_recovery | 0.6219 [0.5054, 0.7825] |
| S0 | within15 | 0.0328 [0.0265, 0.0374] |
| S0 | within30 | 0.0865 [0.0738, 0.1027] |
| S0+PairGeom | bin_accuracy | 0.0160 [0.0122, 0.0204] |
| S0+PairGeom | circular_mae_deg | 96.7135 [94.7742, 97.6803] |
| S0+PairGeom | pooled_recovery | 0.6245 [0.5151, 0.7754] |
| S0+PairGeom | within15 | 0.0375 [0.0262, 0.0487] |
| S0+PairGeom | within30 | 0.0947 [0.0647, 0.1246] |
| S0+$\chi$ | bin_accuracy | 0.6174 [0.6041, 0.6369] |
| S0+$\chi$ | circular_mae_deg | 10.6661 [9.9092, 11.3458] |
| S0+$\chi$ | pooled_recovery | 0.6276 [0.5229, 0.7759] |
| S0+$\chi$ | within15 | 0.8718 [0.8602, 0.8967] |
| S0+$\chi$ | within30 | 0.9527 [0.9465, 0.9609] |
| G0 | bin_accuracy | 0.6260 [0.6110, 0.6468] |
| G0 | circular_mae_deg | 10.5779 [9.7183, 11.2685] |
| G0 | pooled_recovery | 0.6238 [0.5146, 0.7760] |
| G0 | within15 | 0.8730 [0.8596, 0.9042] |
| G0 | within30 | 0.9489 [0.9418, 0.9587] |
| G1 | bin_accuracy | 0.6289 [0.6143, 0.6477] |
| G1 | circular_mae_deg | 10.6773 [9.9308, 11.2807] |
| G1 | pooled_recovery | 0.6181 [0.5072, 0.7717] |
| G1 | within15 | 0.8735 [0.8614, 0.9016] |
| G1 | within30 | 0.9486 [0.9412, 0.9584] |
| Training prior | bin_accuracy | 0.4418 [0.4077, 0.5039] |
| Training prior | circular_mae_deg | 15.0389 [14.3897, 15.3832] |
| Training prior | within15 | 0.7285 [0.6981, 0.7767] |
| Training prior | within30 | 0.8909 [0.8836, 0.8993] |

**Supplementary Data D011.** Recovery and angular accuracy by pretraining seed.

#### 2.1 Common public-inventory-unexposed source subset

The common public-inventory-unmatched subset contains 26 records, 2,148 nucleotides and 2,142 valid angles. It excludes provenance matches to either public encoder inventory and differs from the 3GS5-only exclusion sensitivity. Values average the same three pretraining seeds; inventory absence does not establish family independence.

**Table S11. Pretraining diagnostics on the common inventory-unmatched 26-record subset.**

| Model | Recovery | $A_{15}$ | $A_{30}$ | Bin acc. | MAE (°) |
|---|---|---|---|---|---|
| S0 | 0.5975 | – | – | – | – |
| S0+PairGeom | 0.6040 | – | – | – | – |
| S0+$\chi$ | 0.6027 | 0.8719 | **0.9535** | 0.6173 | 11.0920 |
| G0 | **0.6130** | **0.8772** | 0.9504 | 0.6310 | **10.8729** |
| G1 | 0.5971 | 0.8771 | 0.9500 | **0.6335** | 10.9962 |
| Training prior | – | 0.7521 | 0.8912 | 0.4762 | 14.8900 |

### 3 Ligand-binding results and comparator protocols

All current structural cells use source 100 and validation selection through downstream 120.

**Table S12.** Pretraining and simpler-objective contrasts, with conditional unadjusted 95% intervals. S0 retains sequence CE and SASA. It is not scratch. Full and LoRA remain separate protocols. Complete objective contrasts are provided in Supplementary Data D015.

| Cohort | Contrast | Δpooled | AP [95% CI] | Δmacro | AP [95% CI] |
|---|---|---|---|---|---|
| External 48 | S0 / full – Scratch-N | 0.0535 | [0.0209, 0.0827] | 0.0291 | [0.0057, 0.0530] |
| External 48 | S0 / full – G0 / full | -0.0146 | [-0.0370, 0.0077] | -0.0075 | [-0.0280, 0.0125] |
| External 48 | S0 / LoRA – G0 / LoRA | 0.0260 | [0.0045, 0.0522] | 0.0379 | [0.0164, 0.0642] |
| External 48 | G0 / full – Scratch-N | 0.0681 | [0.0322, 0.0991] | 0.0366 | [0.0138, 0.0615] |
| TR60 test | S0 / full – Scratch-N | 0.0447 | [0.0145, 0.0761] | 0.0347 | [-0.0024, 0.0718] |
| TR60 test | S0 / full – G0 / full | 0.0015 | [-0.0129, 0.0144] | 0.0054 | [-0.0209, 0.0362] |
| TR60 test | S0 / LoRA – G0 / LoRA | 0.0327 | [0.0139, 0.0576] | 0.0462 | [0.0193, 0.0776] |
| TR60 test | G0 / full – Scratch-N | 0.0433 | [0.0156, 0.0723] | 0.0293 | [-0.0027, 0.0609] |

S0 improves external pooled AP over scratch by 0.0535 [0.0209, 0.0827], while its difference from G0/full is unresolved. S0 LoRA exceeds G0 LoRA under the same LoRA protocol. Individual loss-removal effects and joint removal are therefore not interchangeable. No universal auxiliary necessity, equivalence, or biological synergy is asserted. For the directional comparisons described in the main text, G0 minus S0+PairGeom full adaptation is 0.0426 [0.0173, 0.0613], and G0 minus S0+$\chi$ is 0.0225 [0.0012, 0.0405] on external pooled AP. Supplementary Data D015 reports the reverse orientations with reversed signs.

The complete cohort table below includes validation as a fitting diagnostic, not unbiased evaluation.

**Table S13. Complete ligand objective/adaptation results by cohort.** Full and LoRA remain separate protocols. The first part covers the four external sets and TR60 test; the second retains external pooling, 8F0N exclusion and validation diagnostics.

| Cohort | Model | AP | Macro AP | AUROC | MCC .5 / val / oracle |
|---|---|---|---|---|---|
| JL10 | S0 / full | 0.6565 | 0.6691 | 0.7746 | 0.3579 / 0.3777 / 0.3957 |
| JL10 | S0 / LoRA | **0.6776** | **0.7022** | **0.7833** | 0.3524 / 0.3820 / **0.4302** |
| JL10 | S0+PairGeom / full | 0.6502 | 0.6613 | 0.7586 | 0.3481 / 0.3481 / 0.3995 |
| JL10 | S0+PairGeom / LoRA | 0.6535 | 0.6871 | 0.7668 | 0.3794 / 0.3819 / 0.4001 |
| JL10 | S0+$\chi$ / full | 0.6508 | 0.6631 | **0.7826** | 0.3844 / **0.3859** / 0.4024 |
| JL10 | S0+$\chi$ / LoRA | 0.6612 | 0.6704 | 0.7740 | 0.3499 / 0.3521 / 0.3882 |
| JL10 | G0 / full | **0.6790** | **0.6959** | 0.7803 | **0.4009** / 0.3847 / **0.4226** |
| JL10 | G0 / LoRA | 0.6644 | 0.6694 | 0.7796 | **0.4030** / **0.3842** / 0.4150 |
| JL10 | G1 / full | 0.6492 | 0.6665 | 0.7823 | 0.3556 / 0.3679 / 0.4029 |
| JL10 | G1 / LoRA | 0.6610 | 0.6776 | 0.7819 | 0.3820 / 0.3730 / 0.4174 |
| JL10 | Scratch-N | 0.6144 | 0.6483 | 0.7606 | 0.3309 / 0.3493 / 0.3908 |
| RB9 | S0 / full | 0.8069 | 0.8637 | 0.8908 | 0.6254 / 0.5904 / 0.6483 |
| RB9 | S0 / LoRA | **0.8582** | **0.9105** | **0.9112** | **0.6519** / **0.6614** / **0.6865** |
| RB9 | S0+PairGeom / full | 0.8077 | 0.8546 | 0.9020 | 0.6588 / 0.6588 / 0.7287 |
| RB9 | S0+PairGeom / LoRA | 0.8225 | 0.8435 | 0.8924 | 0.5917 / 0.6276 / 0.6607 |
| RB9 | S0+$\chi$ / full | 0.8385 | 0.8791 | 0.9132 | 0.6383 / 0.6631 / 0.7161 |
| RB9 | S0+$\chi$ / LoRA | 0.8302 | 0.8828 | **0.9112** | 0.6393 / 0.6489 / 0.6630 |
| RB9 | G0 / full | **0.8681** | **0.9018** | **0.9181** | 0.6672 / 0.6932 / **0.7393** |
| RB9 | G0 / LoRA | 0.7900 | 0.8363 | 0.8816 | 0.5943 / 0.6153 / 0.6311 |
| RB9 | G1 / full | 0.8655 | 0.8758 | 0.9128 | **0.7083** / **0.7098** / 0.7357 |
| RB9 | G1 / LoRA | 0.8138 | 0.8736 | 0.8932 | 0.5792 / 0.6525 / 0.6577 |
| RB9 | Scratch-N | 0.7703 | 0.8413 | 0.8742 | 0.5857 / 0.6248 / 0.6582 |
| TE18 | S0 / full | 0.5543 | **0.6577** | 0.7353 | 0.4066 / 0.3595 / 0.4365 |
| TE18 | S0 / LoRA | **0.5663** | **0.6811** | **0.7597** | **0.4165** / **0.4063** / **0.4322** |
| TE18 | S0+PairGeom / full | 0.5413 | 0.6556 | 0.7388 | 0.3966 / 0.3966 / 0.4074 |
| TE18 | S0+PairGeom / LoRA | 0.5218 | 0.6066 | 0.7230 | 0.3731 / 0.3660 / 0.3813 |
| TE18 | S0+$\chi$ / full | 0.5463 | 0.6431 | **0.7426** | 0.3937 / 0.3811 / **0.4389** |
| TE18 | S0+$\chi$ / LoRA | 0.5284 | 0.6350 | 0.7224 | 0.3604 / 0.3513 / 0.3753 |
| TE18 | G0 / full | 0.5538 | 0.6369 | 0.7395 | 0.3954 / 0.3920 / 0.4000 |
| TE18 | G0 / LoRA | 0.5443 | 0.6467 | 0.7441 | 0.3680 / 0.3660 / 0.4171 |
| TE18 | G1 / full | **0.5691** | 0.6242 | 0.7413 | **0.4105** / **0.4003** / 0.4226 |
| TE18 | G1 / LoRA | 0.5268 | 0.6200 | 0.7134 | 0.3117 / 0.3509 / 0.3675 |
| TE18 | Scratch-N | 0.5222 | 0.6246 | 0.7215 | 0.3813 / 0.3501 / 0.3813 |
| TL12 | S0 / full | **0.8423** | **0.8453** | 0.8572 | 0.6364 / **0.6804** / 0.6804 |
| TL12 | S0 / LoRA | 0.8145 | **0.8340** | 0.8621 | **0.6694** / 0.6585 / **0.6791** |
| TL12 | S0+PairGeom / full | 0.7887 | 0.8142 | 0.8496 | 0.6316 / 0.6316 / 0.6571 |
| TL12 | S0+PairGeom / LoRA | **0.8149** | 0.8320 | 0.8567 | 0.6594 / 0.6319 / 0.6594 |
| TL12 | S0+$\chi$ / full | 0.8205 | 0.8294 | **0.8581** | 0.6448 / 0.6146 / 0.6568 |
| TL12 | S0+$\chi$ / LoRA | 0.7999 | 0.8178 | 0.8412 | 0.6527 / 0.6358 / 0.6672 |
| TL12 | G0 / full | 0.8276 | 0.8447 | 0.8579 | **0.6665** / 0.6654 / **0.6895** |
| TL12 | G0 / LoRA | 0.8059 | 0.8128 | **0.8630** | 0.6601 / **0.6619** / 0.6711 |
| TL12 | G1 / full | 0.8142 | 0.8317 | 0.8465 | 0.6598 / 0.6588 / 0.6771 |
| TL12 | G1 / LoRA | 0.7920 | 0.8195 | 0.8469 | 0.6470 / 0.6507 / 0.6591 |
| TL12 | Scratch-N | 0.7570 | 0.8018 | 0.8244 | 0.5738 / 0.5768 / 0.5859 |
| TR60_test | S0 / full | **0.8507** | **0.8458** | 0.8908 | 0.6504 / **0.6730** / **0.6779** |
| TR60_test | S0 / LoRA | **0.8586** | **0.8509** | 0.8994 | **0.6676** / **0.6749** / **0.6822** |
| TR60_test | S0+PairGeom / full | 0.8440 | 0.8198 | 0.8931 | 0.6445 / 0.6445 / 0.6681 |
| TR60_test | S0+PairGeom / LoRA | 0.8392 | 0.8379 | **0.9022** | 0.6460 / 0.6463 / 0.6518 |
| TR60_test | S0+$\chi$ / full | 0.8445 | 0.8422 | 0.8915 | 0.6495 / 0.6539 / 0.6619 |
| TR60_test | S0+$\chi$ / LoRA | 0.8421 | 0.8307 | 0.8910 | 0.6299 / 0.6583 / 0.6612 |
| TR60_test | G0 / full | 0.8493 | 0.8404 | **0.8975** | 0.6435 / 0.6523 / 0.6599 |
| TR60_test | G0 / LoRA | 0.8258 | 0.8047 | 0.8777 | 0.6124 / 0.6205 / 0.6512 |
| TR60_test | G1 / full | 0.8440 | 0.8292 | 0.8883 | **0.6507** / 0.6470 / 0.6576 |
| TR60_test | G1 / LoRA | 0.8347 | 0.8292 | 0.8910 | 0.6300 / 0.6369 / 0.6493 |
| TR60_test | Scratch-N | 0.8060 | 0.8111 | 0.8817 | 0.6165 / 0.6161 / 0.6352 |

**Table S13. (continued)**

| Cohort | Model | AP | Macro AP | AUROC | MCC .5 / val / oracle |
|---|---|---|---|---|---|
| external_pooled | S0 / full | 0.6883 | 0.7294 | 0.8054 | 0.4819 / 0.4820 / 0.5032 |
| external_pooled | S0 / LoRA | **0.6938** | **0.7532** | **0.8195** | **0.4946** / **0.5021** / **0.5125** |
| external_pooled | S0+PairGeom / full | 0.6602 | 0.7193 | 0.8006 | 0.4822 / 0.4822 / 0.4982 |
| external_pooled | S0+PairGeom / LoRA | 0.6702 | 0.7081 | 0.8002 | 0.4795 / 0.4791 / 0.4894 |
| external_pooled | S0+$\chi$ / full | 0.6803 | 0.7245 | 0.8126 | 0.4909 / 0.4881 / **0.5177** |
| external_pooled | S0+$\chi$ / LoRA | 0.6742 | 0.7201 | 0.8044 | 0.4729 / 0.4710 / 0.4819 |
| external_pooled | G0 / full | **0.7028** | **0.7369** | **0.8178** | **0.5082** / **0.5074** / 0.5132 |
| external_pooled | G0 / LoRA | 0.6678 | 0.7153 | 0.8080 | 0.4871 / 0.4849 / 0.4910 |
| external_pooled | G1 / full | 0.6934 | 0.7169 | 0.8115 | 0.5041 / 0.5058 / 0.5164 |
| external_pooled | G1 / LoRA | 0.6644 | 0.7156 | 0.8003 | 0.4579 / 0.4824 / 0.4838 |
| external_pooled | Scratch-N | 0.6347 | 0.7003 | 0.7868 | 0.4439 / 0.4523 / 0.4621 |
| external_without_8F0N | S0 / full | 0.6762 | 0.7238 | 0.7989 | 0.4712 / 0.4705 / 0.4924 |
| external_without_8F0N | S0 / LoRA | **0.6841** | **0.7494** | **0.8135** | **0.4839** / **0.4913** / **0.5018** |
| external_without_8F0N | S0+PairGeom / full | 0.6505 | 0.7147 | 0.7943 | 0.4712 / 0.4712 / 0.4884 |
| external_without_8F0N | S0+PairGeom / LoRA | 0.6588 | 0.7016 | 0.7941 | 0.4705 / 0.4709 / 0.4803 |
| external_without_8F0N | S0+$\chi$ / full | 0.6674 | 0.7183 | 0.8069 | 0.4810 / 0.4775 / **0.5083** |
| external_without_8F0N | S0+$\chi$ / LoRA | 0.6643 | 0.7162 | 0.7982 | 0.4619 / 0.4593 / 0.4713 |
| external_without_8F0N | G0 / full | **0.6919** | **0.7313** | **0.8118** | **0.4980** / **0.4969** / 0.5026 |
| external_without_8F0N | G0 / LoRA | 0.6553 | 0.7089 | 0.8015 | 0.4772 / 0.4748 / 0.4812 |
| external_without_8F0N | G1 / full | 0.6822 | 0.7116 | 0.8055 | 0.4934 / 0.4962 / 0.5069 |
| external_without_8F0N | G1 / LoRA | 0.6526 | 0.7090 | 0.7932 | 0.4468 / 0.4712 / 0.4726 |
| external_without_8F0N | Scratch-N | 0.6219 | 0.6937 | 0.7795 | 0.4329 / 0.4407 / 0.4509 |
| validation | S0 / full | 0.8815 | 0.9015 | 0.9034 | 0.6882 / **0.7075** / **0.7086** |
| validation | S0 / LoRA | 0.8709 | 0.8953 | 0.8982 | 0.6750 / 0.6881 / **0.6912** |
| validation | S0+PairGeom / full | 0.8743 | 0.8885 | 0.8963 | 0.6933 / 0.6933 / 0.6965 |
| validation | S0+PairGeom / LoRA | **0.8813** | **0.9072** | **0.9078** | **0.6821** / 0.6901 / **0.6912** |
| validation | S0+$\chi$ / full | 0.8732 | 0.8875 | 0.8930 | 0.6680 / 0.6782 / 0.6805 |
| validation | S0+$\chi$ / LoRA | 0.8652 | 0.8829 | 0.8909 | 0.6664 / 0.6809 / 0.6841 |
| validation | G0 / full | **0.8854** | **0.9065** | **0.9057** | **0.6969** / 0.7028 / 0.7049 |
| validation | G0 / LoRA | 0.8716 | 0.8846 | 0.8985 | 0.6740 / 0.6773 / 0.6773 |
| validation | G1 / full | 0.8732 | 0.8845 | 0.8969 | 0.6838 / 0.6955 / 0.6963 |
| validation | G1 / LoRA | 0.8751 | 0.8897 | 0.8980 | 0.6742 / **0.6912** / **0.6912** |
| validation | Scratch-N | 0.8432 | 0.8604 | 0.8797 | 0.6151 / 0.6283 / 0.6336 |

**Supplementary Data D015.** Complete CoBRA objective and adaptation contrasts.

#### 3.1 Individual runs and probability ensembles

Individual-seed AP, mean and sample SD, and probability-ensemble AP are shown for TR60 test and the external 48-record set.

**Average precision (AP).**

**Table S14. CoBRA individual-seed and probability-ensemble AP.**

| Cohort | Model | Seed 0 | Seed 1 | Seed 2 | Mean $\pm$ SD | Ensemble |
|---|---|---|---|---|---|---|
| TR60_test | S0 / full | 0.8232 | 0.8252 | 0.8116 | 0.8200 $\pm$ 0.0073 | 0.8507 |
| TR60_test | S0 / LoRA | 0.8206 | 0.8348 | 0.8217 | 0.8257 $\pm$ 0.0079 | 0.8586 |
| TR60_test | S0+PairGeom / full | 0.8169 | 0.8091 | 0.8161 | 0.8141 $\pm$ 0.0043 | 0.8440 |
| TR60_test | S0+PairGeom / LoRA | 0.8046 | 0.8070 | 0.8136 | 0.8084 $\pm$ 0.0047 | 0.8392 |
| TR60_test | S0+$\chi$ / full | 0.8284 | 0.8226 | 0.8180 | 0.8230 $\pm$ 0.0052 | 0.8445 |
| TR60_test | S0+$\chi$ / LoRA | 0.8152 | 0.8133 | 0.8088 | 0.8124 $\pm$ 0.0033 | 0.8421 |
| TR60_test | G0 / full | 0.8317 | 0.8054 | 0.8234 | 0.8202 $\pm$ 0.0134 | 0.8493 |
| TR60_test | G0 / LoRA | 0.7826 | 0.7929 | 0.8038 | 0.7931 $\pm$ 0.0106 | 0.8258 |
| TR60_test | G1 / full | 0.8139 | 0.7911 | 0.8098 | 0.8049 $\pm$ 0.0121 | 0.8440 |
| TR60_test | G1 / LoRA | 0.8087 | 0.7952 | 0.8119 | 0.8053 $\pm$ 0.0089 | 0.8347 |
| TR60_test | Scratch-N | 0.7976 | 0.7618 | 0.7331 | 0.7642 $\pm$ 0.0323 | 0.8060 |
| external_pooled | S0 / full | 0.6473 | 0.6551 | 0.6831 | 0.6618 $\pm$ 0.0188 | 0.6883 |
| external_pooled | S0 / LoRA | 0.6485 | 0.6961 | 0.6641 | 0.6696 $\pm$ 0.0243 | 0.6938 |
| external_pooled | S0+PairGeom / full | 0.6553 | 0.6453 | 0.6360 | 0.6456 $\pm$ 0.0096 | 0.6602 |
| external_pooled | S0+PairGeom / LoRA | 0.6250 | 0.6405 | 0.6729 | 0.6461 $\pm$ 0.0244 | 0.6702 |
| external_pooled | S0+$\chi$ / full | 0.6539 | 0.6721 | 0.6663 | 0.6641 $\pm$ 0.0093 | 0.6803 |
| external_pooled | S0+$\chi$ / LoRA | 0.6613 | 0.6307 | 0.6732 | 0.6551 $\pm$ 0.0220 | 0.6742 |
| external_pooled | G0 / full | 0.6759 | 0.6487 | 0.6836 | 0.6694 $\pm$ 0.0183 | 0.7028 |
| external_pooled | G0 / LoRA | 0.6506 | 0.6555 | 0.6431 | 0.6497 $\pm$ 0.0063 | 0.6678 |
| external_pooled | G1 / full | 0.6636 | 0.6692 | 0.6346 | 0.6558 $\pm$ 0.0186 | 0.6934 |
| external_pooled | G1 / LoRA | 0.6381 | 0.6394 | 0.6487 | 0.6421 $\pm$ 0.0058 | 0.6644 |
| external_pooled | Scratch-N | 0.6194 | 0.6112 | 0.5703 | 0.6003 $\pm$ 0.0263 | 0.6347 |

#### 3.2 Same-cohort named implementations

Aggregate results are in main Table 2; a machine-readable copy is provided in Supplementary Data D017.

CoBRA-M3 uses a frozen RiNALMo compatibility path (SDPA rather than validated native attention) with 100 head epochs. RNABind uses the same TR60 splits and 120 epochs with a numerically stable logits loss, not the complete native 300-epoch recipe. Their three-model ensembles are local measurements. SMARTPocket is one released checkpoint averaged across five noisy passes. Its native training exposure is not verified. These rows are not exact native reproductions or a comprehensive modern-method leaderboard.

**Table S15. Comparison protocols for locally evaluated ligand-binding methods.**

| Comparison | What is matched | What is not established |
|---|---|---|
| G0/S0 vs Scratch | Input, architecture, data, full-120 budget, validation rule | Source exposure is asymmetric. Separate scratch-optimal hyperparameters were not searched. |
| G0 vs CoBRA-M3 | Admitted TR60 splits and scored labels | LM/structural inputs, 100 vs 120 epochs, attention runtime and source exposure differ. No current-G0 paired interval. |
| G0 vs RNABind | Admitted TR60 splits, labels and local 120 epochs | Inputs/architecture differ. Native 300-epoch recipe is not reproduced. No current-G0 paired interval. |
| G0 vs released SMARTPocket | Shared CoBRA scored cohort in the local adapter table | Released fitting history, representation and stochastic-pass averaging differ from three independently fitted models. |
| G0 vs published SMARTPocket apo/holo | Twelve PDB identities, per-PDB metric names and equal-PDB summary | Published rounded values only. Residue labels/admission, source exposure and ensemble recipe are not fully matched. No paired significance test. |
| RSR vs literature | Task name and reported endpoint definitions | Native input, split handling, training/head/selection and replication contracts are not reproduced. |

On the matched TE18 cohort, local SMARTPocket AP is 0.6810 versus G0 0.5538. Inputs and training exposure differ between these methods.

**Supplementary Data D019.** Set-specific metrics of local ligand-binding implementations.

#### 3.3 Released GRASSP and local external evaluation

GRASSP is locally evaluated with COBRA_SEED1 native feature graphs on JL10 (10 records), TL12 (12) and filtered TE18 (17). Normalized PDB–chain identities and ordered labels match all 39 common records. The release lacks residue identifiers and sequences, so physical residue provenance cannot be independently reconstructed. Its training graph collection includes TL12 PDB 8F0N. Six RB9 feature graphs and a complete raw feature-generation pipeline are unavailable; the three available RB9 graphs are not reported as if they represented RB9. No 48-record pooled GRASSP result is reported. The fixed CoBRA checkpoint differs from the published TE18 protocol using the separately trained TR60 model.

Local GRASSP AP/AUROC is 0.6484/0.7683 on JL10, 0.8362/0.8745 on TL12 and 0.4598/0.6018 on TE18. Main Table 3 and Supplementary Table S16 report AUROC and threshold-dependent MCC.

**Table S16. Local ligand-binding benchmarks: AUROC and threshold sensitivity.**

**Local re-evaluation**

**RB9 (9 records)**

| Method | AUROC | MCC .5 | MCC oracle |
|---|---|---|---|
| CoBRA-M3 | 0.8537 | 0.6390 | 0.6631 |
| RNABind | 0.8825 | 0.6441 | 0.6527 |
| SMARTPocket | 0.8899 | 0.5002 | 0.6211 |
| GRASSP (released) | – | – | – |
| S0 / full | 0.8908 | 0.6254 | 0.6483 |
| G0 / full | **0.9181** | <u>0.6672</u> | **0.7393** |
| G1 / full | <u>0.9128</u> | **0.7083** | <u>0.7357</u> |

**JL10 (10 records)**

| Method | AUROC | MCC .5 | MCC oracle |
|---|---|---|---|
| CoBRA-M3 | 0.7424 | 0.2963 | 0.3785 |
| RNABind | 0.6888 | 0.2945 | 0.3344 |
| SMARTPocket | 0.6936 | 0.2038 | 0.3111 |
| GRASSP (released) | 0.7683 | 0.3566 | 0.3972 |
| S0 / full | 0.7746 | <u>0.3579</u> | 0.3957 |
| G0 / full | <u>0.7803</u> | **0.4009** | **0.4226** |
| G1 / full | **0.7823** | 0.3556 | <u>0.4029</u> |

**TL12 (12 records)**

| Method | AUROC | MCC .5 | MCC oracle |
|---|---|---|---|
| CoBRA-M3 | <u>0.8627</u> | 0.5552 | 0.5921 |
| RNABind | 0.8483 | 0.5585 | 0.5676 |
| SMARTPocket | 0.8289 | 0.5065 | 0.5412 |
| GRASSP (released) | **0.8745** | 0.5959 | 0.6449 |
| S0 / full | 0.8572 | 0.6364 | <u>0.6804</u> |
| G0 / full | 0.8579 | **0.6665** | **0.6895** |
| G1 / full | 0.8465 | <u>0.6598</u> | 0.6771 |

**TE18 (17 records)**

| Method | AUROC | MCC .5 | MCC oracle |
|---|---|---|---|
| CoBRA-M3 | 0.6436 | 0.1726 | 0.2422 |
| RNABind | 0.6751 | 0.2461 | 0.2956 |
| SMARTPocket | **0.7840** | 0.3405 | **0.4547** |
| GRASSP (released) | 0.6018 | 0.1252 | 0.2031 |
| S0 / full | 0.7353 | <u>0.4066</u> | <u>0.4365</u> |
| G0 / full | 0.7395 | 0.3954 | 0.4000 |
| G1 / full | <u>0.7413</u> | **0.4105** | 0.4226 |

Each dataset is shown in a separate subtable with its own AUROC and MCC columns. Published results are reported separately in Table S18. Local MCC .5 and oracle use a fixed 0.5 threshold and a test-label oracle threshold, respectively, the latter a nondeployable diagnostic ceiling. Set-level validation-selected MCC and AP are reported in Supplementary Table S13 for the BRIDGE variants and Scratch-N, and in Supplementary Data D019 for CoBRA-M3, RNABind and SMARTPocket. Local test sets contain 9/10/12/17 records, with 4F8U excluded from TE18, compared with 18 in the published TE18 set. CoBRA-M3/RNABind are compatibility adapters. Local SMARTPocket averages five passes of one released checkpoint. BRIDGE uses the same three-model full-adaptation ensemble (120 epochs) for every set. Published GRASSP uses original TR60 for TE18 and combined HARIBOSS–TR60 training for JL10/TL12. Local GRASSP instead uses one released CoBRA checkpoint (COBRA_SEED1) and its native feature graphs for all three supported sets. Normalized PDB–chain identifiers and ordered label arrays match the common test records; released graphs lack residue identifiers and sequences, so physical residue provenance cannot be independently reconstructed. This is a released-model re-evaluation, not a reproduction of the published TE18 protocol. Its released training graph collection contains PDB 8F0N from TL12. Full RB9 evaluation is unavailable because six feature graphs and the raw feature-generation pipeline are not released. Dashes indicate unavailable results. Bold and underline mark the best and second-best distinct displayed estimates within each protocol block and metric, respectively; ties share emphasis. Columns with fewer than two estimates are not ranked. No ranking across protocol blocks or statistical superiority is implied. AUROC, area under the receiver operating characteristic curve; MCC, Matthews correlation coefficient.

The separate native release evaluation below retains its original validation43/test44/TE18=18 supports. Its test AP 0.8158 is not a shared-43-record TR60 test estimate. Those original-cohort values must not be substituted into the local common-set comparison.

**Table S17. GRASSP evaluation on native release cohorts.**

| Native cohort | Graphs | AP | Macro AP | AUROC | MCC .5 / val / oracle |
|---|---|---|---|---|---|
| JL10 | 10 | 0.6484 | 0.6863 | 0.7683 | 0.3566 / 0.3478 / 0.3972 |
| TE18 | 18 | 0.4524 | 0.5391 | 0.5982 | 0.1071 / 0.1088 / 0.1919 |
| TL12 | 12 | 0.8362 | 0.8251 | 0.8745 | 0.5959 / 0.6026 / 0.6449 |
| TR60_test | 44 | 0.8158 | 0.8164 | 0.8835 | 0.5928 / 0.5854 / 0.6371 |
| validation | 43 | 0.8233 | 0.8034 | 0.8705 | 0.6129 / 0.6212 / 0.6229 |

#### 3.4 Published ligand-binding benchmark context

Table S18 retains every published-method row, including missing values, under each paper's reported protocol. Published TE18 contains 18 records, whereas the local filtered set contains 17. Reported MCC thresholds and averaging differ. Published and local estimates are not ranked together.

#### Table S18. Task-specific ligand-binding site benchmarks.

##### Published protocols

**RB9**

| Method | AUROC | MCC reported |
|---|---|---|
| Rsite[a] | 0.528 | 0.159 |
| Rsite2[a] | 0.575 | 0.072 |
| RNAsite[a] | 0.790 | 0.426 |
| RBind[a] | 0.592 | 0.287 |
| RLBind[a] | 0.851 | 0.546 |
| ZHmolReSTasite[a] | 0.786 | 0.398 |
| RNET[a] | – | 0.465 |
| RNABind[a] | <u>0.865</u> | – |
| GATRSite[a] | 0.850 | <u>0.570</u> |
| MVRBind [13] | – | – |
| CoBRA-M3 [4] | 0.844 | 0.557 |
| SMARTPocket [14] | **0.880** ± 0.016 | **0.574** ± 0.032 |
| GRASSP [15] | – | – |

**JL10**

| Method | AUROC | MCC reported |
|---|---|---|
| Rsite[a] | 0.477 | -0.046 |
| Rsite2[a] | 0.504 | 0.007 |
| RNAsite[a] | – | – |
| RBind[a] | 0.532 | 0.083 |
| RLBind[a] | – | – |
| ZHmolReSTasite[a] | 0.592 | 0.211 |
| RNET[a] | – | – |
| RNABind[a] | 0.667 | – |
| GATRSite[a] | – | – |
| MVRBind [13] | – | – |
| CoBRA-M3 [4] | <u>0.739</u> | 0.293 |
| SMARTPocket [14] | 0.731 ± 0.014 | <u>0.330</u> ± 0.040 |
| GRASSP [15] | **0.760** | **0.343** |

**TL12**

| Method | AUROC | MCC reported |
|---|---|---|
| Rsite[a] | – | – |
| Rsite2[a] | – | – |
| RNAsite[a] | – | – |
| RBind[a] | – | – |
| RLBind[a] | – | – |
| ZHmolReSTasite[a] | 0.704 | 0.440 |
| RNET[a] | – | – |
| RNABind[a] | 0.754 | – |
| GATRSite[a] | – | – |
| MVRBind [13] | – | – |
| CoBRA-M3 [4] | 0.816 | <u>0.546</u> |
| SMARTPocket [14] | <u>0.832</u> ± 0.020 | 0.545 ± 0.020 |
| GRASSP [15] | **0.874** | **0.611** |

**TE18**

| Method | AUROC | MCC reported |
|---|---|---|
| Rsite[a] | 0.474 | 0.071 |
| Rsite2[a] | 0.509 | 0.010 |
| RNAsite[a] | 0.776 | 0.253 |
| RBind[a] | 0.559 | 0.187 |
| RLBind[a] | 0.720 | 0.324 |
| ZHmolReSTasite[a] | 0.709 | 0.327 |
| RNET[a] | – | 0.307 |
| RNABind[a] | 0.737 | – |
| GATRSite[a] | <u>0.780</u> | 0.320 |
| MVRBind [13] | 0.745 | 0.351 |
| CoBRA-M3 [4] | 0.647 | 0.190 |
| SMARTPocket [14] | **0.781** ± 0.007 | **0.410** ± 0.026 |
| GRASSP [15] | 0.746 | <u>0.359</u> |

[a] Values for the nine marked baselines were transcribed from Table 1 of SMARTPocket [14] (preprint version posted 19 June 2026). That table attributes RLBind, RNABind, RNET and GATRSite to their original publications, and the other five added methods to the ZHmolReSTasite publication. The MVRBind, CoBRA-M3, SMARTPocket and GRASSP rows follow their respective publications [13, 4, 14, 15]. These are published summaries, not local reruns. The AUC heading in SMARTPocket Table 1 denotes AUROC. Missing rows and unreported metrics are shown as dashes; no values are inferred. The spelling RNAsite follows the table. SMARTPocket values are mean±SD across five random seeds. Published protocols and MCC thresholds are retained. GRASSP uses original TR60 for TE18 and combined HARIBOSS–TR60 training for JL10/TL12. Published TE18 has 18 records; the local evaluation in Table S16 contains 17 after excluding 4F8U. Bold and underline mark the highest and second-highest distinct displayed estimates per dataset and metric among all 13 published methods; ties share emphasis, and SD is not ranked. This is descriptive, not a matched-protocol superiority test. AUROC, area under the receiver operating characteristic curve; MCC, Matthews correlation coefficient.

### 4 Paired conformation results

**Table S19. Apo-focused conformational inference with holo reference values: full-adaptation (120-epoch) three-seed probability ensembles and TR60-fitted GNN heads.**

| Model | AP | Macro AP | AUROC | MCC .5 | MCC val | MCC oracle |
|---|---|---|---|---|---|---|
| **Apo** | | | | | | |
| S0 | 0.3478 | 0.5250 | 0.8053 | 0.3378 | 0.3389 | 0.3594 |
| S0+PairGeom | 0.3484 | 0.5344 | 0.8219 | <u>0.3754</u> | <u>0.3754</u> | 0.3999 |
| S0+$\chi$ | **0.4337** | <u>0.5597</u> | <u>0.8320</u> | 0.3565 | 0.3698 | **0.4433** |
| G0 | 0.3993 | 0.5588 | **0.8435** | 0.3337 | 0.3411 | 0.4013 |
| G1 | <u>0.4189</u> | **0.5683** | 0.8239 | **0.3852** | **0.3977** | <u>0.4221</u> |
| Scratch-N | 0.2533 | 0.3537 | 0.7580 | 0.2418 | 0.2640 | 0.2854 |
| gRNAde | 0.2428 | 0.3835 | 0.7338 | 0.2332 | 0.2327 | 0.2690 |
| NA-MPNN | 0.4120 | 0.5312 | 0.7967 | 0.3253 | 0.3590 | 0.3843 |
| **Holo** | | | | | | |
| S0 | 0.7144 | 0.7406 | 0.9506 | 0.4886 | **0.5972** | 0.6618 |
| S0+PairGeom | 0.6654 | 0.7476 | 0.9443 | <u>0.5447</u> | 0.5447 | 0.6494 |
| S0+$\chi$ | **0.7422** | **0.7745** | <u>0.9546</u> | 0.5200 | <u>0.5944</u> | <u>0.6990</u> |
| G0 | 0.6924 | 0.7354 | **0.9555** | 0.5322 | 0.5360 | 0.6882 |
| G1 | <u>0.7378</u> | <u>0.7603</u> | 0.9533 | **0.5548** | 0.5498 | **0.7345** |
| Scratch-N | 0.4657 | 0.5279 | 0.8857 | 0.4386 | 0.4228 | 0.4698 |
| gRNAde | 0.5275 | 0.5585 | 0.9021 | 0.4523 | 0.4364 | 0.5028 |
| NA-MPNN | 0.6635 | 0.6679 | 0.9324 | 0.4849 | 0.5105 | 0.6548 |

Bold and underline mark the best and second-best distinct values among all displayed models, including released encoders, within each state and metric. Emphasis is descriptive. Released-encoder inputs and pretraining exposure differ from BRIDGE. Public frozen/LoRA results are supplementary. The panel contains six targets and seven graphs per state, with 605 apo and 607 holo labelled residues. Macro AP weights targets equally. Holo identities 3SUX, 2YIE, 4LVX and 2L8H occur in TR60 training. Their apo partners are exposure-linked. The other two pairs have no direct match, which does not establish family independence. Ties at the displayed precision share emphasis. MCC .5 uses threshold 0.5; val uses the validation-selected threshold; oracle uses test labels to maximize MCC and is a nondeployable diagnostic ceiling. AP, average precision. AUROC, area under the receiver operating characteristic curve; MCC, Matthews correlation coefficient.

The pooled contrasts are included in Table S20; Supplementary Data D024 preserves the corresponding four-row extract.

**Table S20. Complete pooled and target-macro conformation AP contrasts.**

| State | Contrast | ΔAP [95% CI] | Δmacro AP [95% CI] |
|---|---|---|---|
| apo | S0 / full – G0 / full | -0.0515 [-0.1197, 0.0237] | -0.0338 [-0.0947, 0.0264] |
| apo | S0 / full – Scratch-N | 0.0945 [0.0332, 0.2135] | 0.1713 [0.0199, 0.3719] |
| apo | S0+PairGeom / full – G0 / full | -0.0509 [-0.1047, 0.0274] | -0.0244 [-0.0681, 0.0088] |
| apo | S0+PairGeom / full – Scratch-N | 0.0951 [0.0120, 0.2819] | 0.1807 [0.0163, 0.3604] |
| apo | S0+$\chi$ / full – G0 / full | 0.0345 [-0.0401, 0.1302] | 0.0009 [-0.0956, 0.1188] |
| apo | S0+$\chi$ / full – Scratch-N | 0.1804 [0.0800, 0.2819] | 0.2060 [0.0969, 0.3050] |
| apo | G0 / full – Scratch-N | 0.1460 [0.0403, 0.2851] | 0.2051 [0.0361, 0.3766] |
| apo | G1 / full – G0 / full | 0.0196 [-0.0244, 0.0735] | 0.0095 [-0.0417, 0.0610] |
| apo | G1 / full – Scratch-N | 0.1656 [0.0376, 0.3230] | 0.2146 [0.0338, 0.4126] |
| holo | S0 / full – G0 / full | 0.0220 [-0.0653, 0.0878] | 0.0052 [-0.0331, 0.0563] |
| holo | S0 / full – Scratch-N | 0.2486 [0.0998, 0.4115] | 0.2126 [0.0139, 0.4229] |
| holo | S0+PairGeom / full – G0 / full | -0.0270 [-0.0871, 0.0760] | 0.0122 [-0.0423, 0.0928] |
| holo | S0+PairGeom / full – Scratch-N | 0.1996 [0.0408, 0.4226] | 0.2197 [0.0523, 0.4167] |
| holo | S0+$\chi$ / full – G0 / full | 0.0498 [-0.0350, 0.1232] | 0.0391 [-0.0339, 0.1236] |
| holo | S0+$\chi$ / full – Scratch-N | 0.2764 [0.1461, 0.4428] | 0.2466 [0.0974, 0.4243] |
| holo | G0 / full – Scratch-N | 0.2266 [0.0509, 0.4424] | 0.2074 [0.0053, 0.4251] |
| holo | G1 / full – G0 / full | 0.0454 [-0.0687, 0.1042] | 0.0249 [-0.0691, 0.1056] |
| holo | G1 / full – Scratch-N | 0.2720 [0.1257, 0.4422] | 0.2324 [0.0744, 0.4178] |

#### 4.1 Three-seed repeatability and ensembles

The same three TR60 fits are applied to both states. Individual-seed AP, mean and sample SD, and probability-ensemble AP are reported separately.

**Table S21. Conformation individual-seed and probability-ensemble AP.**

| Cohort | Model | Seed 0 | Seed 1 | Seed 2 | Mean ± SD | Ensemble |
|---|---|---|---|---|---|---|
| apo | S0 / full | 0.3470 | 0.2996 | 0.3106 | 0.3191 ± 0.0248 | 0.3478 |
| apo | S0+PairGeom / full | 0.3988 | 0.2310 | 0.3696 | 0.3331 ± 0.0896 | 0.3484 |
| apo | S0+$\chi$ / full | 0.5062 | 0.3799 | 0.2691 | 0.3851 ± 0.1186 | 0.4337 |
| apo | G0 / full | 0.3685 | 0.2735 | 0.3494 | 0.3305 ± 0.0502 | 0.3993 |
| apo | G1 / full | 0.3587 | 0.3785 | 0.3046 | 0.3473 ± 0.0383 | 0.4189 |
| apo | Scratch-N | 0.2459 | 0.2550 | 0.2257 | 0.2422 ± 0.0150 | 0.2533 |
| holo | S0 / full | 0.6555 | 0.6231 | 0.6359 | 0.6382 ± 0.0163 | 0.7144 |
| holo | S0+PairGeom / full | 0.6479 | 0.5776 | 0.6353 | 0.6203 ± 0.0375 | 0.6654 |
| holo | S0+$\chi$ / full | 0.7540 | 0.6761 | 0.5651 | 0.6651 ± 0.0949 | 0.7422 |
| holo | G0 / full | 0.6660 | 0.6347 | 0.6571 | 0.6526 ± 0.0161 | 0.6924 |
| holo | G1 / full | 0.6634 | 0.7063 | 0.6082 | 0.6593 ± 0.0492 | 0.7378 |
| holo | Scratch-N | 0.4062 | 0.5153 | 0.3542 | 0.4252 ± 0.0822 | 0.4657 |

#### 4.2 Per-target results and influence

**Table S22. Per-target conformation AP and pretrained-minus-scratch differences.**

| State | PDB | Pos./res. | S0 AP | G0 AP | Scratch AP | ΔS0–scratch | ΔG0–scratch |
|---|---|---|---|---|---|---|---|
| apo | 1ANR | 10/29 | 0.3928 | **0.4946** | 0.3292 | 0.0637 | 0.1655 |
| apo | 3D0X | 8/161 | 0.2486 | 0.2412 | **0.2718** | -0.0232 | -0.0305 |
| apo | 3SUY | 5/101 | **1.0000** | 0.9250 | 0.3780 | 0.6220 | 0.5470 |
| apo | 6WJR | 12/109 | 0.5615 | 0.5336 | **0.5866** | -0.0251 | -0.0530 |
| apo | 6XJZ | 14/116 | 0.3317 | **0.3990** | 0.1959 | 0.1358 | 0.2031 |
| apo | 7KD1 | 15/89 | 0.6155 | **0.7594** | 0.3608 | 0.2546 | 0.3986 |
| holo | 2L8H | 10/29 | **0.8675** | 0.7400 | 0.7426 | 0.1249 | -0.0026 |
| holo | 3D0U | 8/161 | 0.1717 | 0.2150 | **0.3440** | -0.1724 | -0.1290 |
| holo | 3SUX | 5/101 | **1.0000** | **1.0000** | 0.3699 | 0.6301 | 0.6301 |
| holo | 2YIE | 12/111 | 0.8266 | **0.8784** | 0.5182 | 0.3084 | 0.3602 |
| holo | 6XJQ | 14/116 | 0.5777 | **0.5790** | 0.2256 | 0.3521 | 0.3534 |
| holo | 4LVX | 15/89 | **1.0000** | **1.0000** | 0.9674 | 0.0326 | 0.0326 |

Per-target differences use matched labels and three-seed mean probabilities; aggregate paired uncertainty is reported separately.

**Table S23. Leave-one-target-out influence on conformation AP.**

| State | Omitted pair | Pooled ΔAP | Macro ΔAP |
|---|---|---|---|
| apo | 1ANR/2L8H | 0.1310 | 0.2130 |
| apo | 3D0X/3D0U | 0.1849 | 0.2522 |
| apo | 3SUY/3SUX | 0.1237 | 0.1367 |
| apo | 6WJR/2YIE | 0.1809 | 0.2567 |
| apo | 6XJZ/6XJQ | 0.1255 | 0.2055 |
| apo | 7KD1/4LVX | 0.1391 | 0.1664 |
| holo | 1ANR/2L8H | 0.3699 | 0.2495 |
| holo | 3D0X/3D0U | 0.2308 | 0.2747 |
| holo | 3SUY/3SUX | 0.1939 | 0.1229 |
| holo | 6WJR/2YIE | 0.2128 | 0.1769 |
| holo | 6XJZ/6XJQ | 0.1692 | 0.1783 |
| holo | 7KD1/4LVX | 0.2112 | 0.2424 |

These leave-one-target-out values are G0 minus Scratch pooled/macro AP. They quantify influence without constructing extra independent samples. No target is excluded from the reported cohort.

#### 4.3 Local and published SMARTPocket comparisons

Table S24 extends main Table 6 with fixed-threshold and per-target oracle MCC and keeps the published reference separate. Each column is an equal-target mean, not pooled AUROC/MCC. Oracle MCC optimizes independently within each test target before averaging and is not a deployable estimate.

**Table S24. Target-macro SMARTPocket comparison of selected configurations (S0+PairGeom and S0+$\chi$ are not included).**

| | Apo: target macro | | | | Holo: target macro | | | |
|---|---|---|---|---|---|---|---|---|
| Method | AUROC | MCC .5 | MCC val | MCC oracle | AUROC | MCC .5 | MCC val | MCC oracle |
| **Local evaluation** | | | | | | | | |
| S0 / full | 0.8369 | 0.3989 | 0.3683 | <u>0.5913</u> | 0.9398 | 0.5937 | <u>0.6627</u> | 0.7190 |
| G0 / full | <u>0.8559</u> | 0.3526 | 0.3735 | 0.5527 | <u>0.9429</u> | <u>0.6188</u> | 0.6254 | <u>0.7626</u> |
| G1 / full | 0.8310 | <u>0.4251</u> | <u>0.4408</u> | 0.5558 | 0.9379 | 0.6171 | 0.6059 | 0.7587 |
| Scratch-N | 0.7645 | 0.2228 | 0.2513 | 0.4426 | 0.8858 | 0.4934 | 0.4530 | 0.6101 |
| SMARTPocket local | **0.8790** | **0.6275** | **0.5580** | **0.7334** | **0.9732** | **0.8099** | **0.7686** | **0.8898** |
| **Published reference** | | | | | | | | |
| SMARTPocket reported | 0.8017 | 0.5112 | – | – | 0.9854 | 0.8341 | – | – |

Each metric is an equal-weight mean over six target PDBs per state, unlike the pooled AUROC in Table S19. Bold and underline rank the best and second-best distinct estimates within the local block; the published reference is not ranked against local reruns. Emphasis is descriptive. BRIDGE uses three-model full-adaptation (120-epoch) probability ensembles. MCC .5 uses a fixed cutoff. MCC val uses the TR60 validation-selected threshold. Oracle MCC optimizes the threshold separately using each test target's labels, then averages the six maxima. It is a diagnostic ceiling, not deployable performance or one shared threshold. SMARTPocket local uses the released checkpoint with five seeded stochastic passes, full RNA heavy-atom inputs, and the same six target identities, residue mappings, 4 Å labels and evaluation masks as BRIDGE. Its MCC val threshold (0.28) was selected on TR60 validation without using the six test targets. The reported row quotes published SMARTPocket AUROC and fixed MCC. Unavailable validation/oracle values are dashes. Four holo identities (3SUX, 2YIE, 4LVX, 2L8H) are training-seen for BRIDGE and their apo partners are exposure-linked. The SMARTPocket configured validation split contains 2YIE; checkpoint exposure is not proven absent. Model inputs and training exposure differ, so no exposure-clean or paired superiority claim is made. Ties at the displayed precision share emphasis. AUROC, area under the receiver operating characteristic curve; MCC, Matthews correlation coefficient.

SMARTPocket local was replayed on the same six target identities, residue mappings, 4 Å labels and masks as BRIDGE. Five seeded stochastic passes of one released checkpoint use full RNA heavy atoms. Threshold 0.28 was selected on TR60 validation without using these six test targets. Apo/holo target-macro AUROC is 0.8790/0.9732; MCC val is 0.5580/0.7686. Four holo targets occur in BRIDGE downstream training, while the configured SMARTPocket validation split contains 2YIE; checkpoint exposure is not proven absent. Inputs and fitting histories differ, precluding exposure-clean superiority claims.

The published per-PDB SMARTPocket rows are contextual references, distinct from the local evaluation. Published AP is unavailable, and matched PDB identities do not guarantee matched residues, labels or exposure.

BRIDGE's fixed cutoff is $p \geq 0.5$; the released SMARTPocket evaluator rounds probabilities. Validation MCC retains each model's TR60-selected threshold. Per-target oracle MCC includes the all-negative endpoint and uses test labels. The fixed/validation/oracle differences diagnose operating-point sensitivity without demonstrating probability calibration.

**Supplementary Data D030.** Per-PDB SMARTPocket comparison with local and published results.

##### 4.4 SMARTFlex49: retrospective paired-conformation evaluation

SMARTFlex49 was derived from the archived SMARTFlexDB release [16]. From 105 eligible pairs after the existing BRIDGE exposure screening, we removed the 44-pair 1X9C connected component after inspecting the full-cohort results, then excluded 12 pairs in components 1PJY, 5OR0 and 5V16 that intersected SMARTPocket's configured training PDB list. The retained 49 pairs comprise 57 unique PDB structures and 11 connected components. This is a retrospective subset, not a prospectively untouched holdout. Its pair membership and scoring masks are fixed across models; repeated structures retain their pair weights and do not constitute independent observations.

Labels reconstruct the documented CoBRA policy [4]: a residue is positive if any heavy atom lies within 4 Å of any eligible nonpolymer ligand or ion in the first coordinate model. Contacts are unioned over all deposited ligand instances and alternate locations, without symmetry expansion. HOH, SO4, PO4, GOL, EDO, PEG and covalently polymer-connected components are excluded; TRS, SPM and N3D remain eligible under this explicit list. Holo labels are projected to apo through the frozen alignments, without adding a new RNA-length filter. This endpoint differs from the database's 8 Å pocket definition. It reconstructs documented rules rather than exact archived training annotations: an independent audit of 2,083 residues from 44 shared training chains retained 74 label mismatches; those audit chains were not added to this test cohort.

All three-fit ensembles reuse the existing TR60 full-adaptation checkpoints; SMARTPocket uses one released checkpoint averaged over five stochastic passes. BA and MCC retain validation-MCC-selected cutoffs: S0 0.68, S0+PairGeom 0.50, S0+$\chi$ 0.80, G0/G1 0.54, Scratch-N 0.66, gRNAde 0.48, NA-MPNN 0.64 and SMARTPocket 0.28. SMARTPocket inherits its cutoff from the earlier validation evaluation. No fitting or threshold reselection used SMARTFlex49 labels.

**Table S25. SMARTFlex49: pooled balanced accuracy, MCC and AUROC under the documented CoBRA 4 Å label policy.**

| | Apo | | | Holo | | |
|---|---|---|---|---|---|---|
| Model | BA val | MCC val | AUROC | BA val | MCC val | AUROC |
| S0 | 0.6370 | 0.2648 | <u>0.7137</u> | 0.6840 | 0.3630 | <u>0.7648</u> |
| S0+PairGeom | 0.6501 | 0.2721 | 0.6916 | 0.6718 | 0.3149 | 0.7326 |
| S0+$\chi$ | 0.6289 | 0.2822 | 0.6991 | 0.6797 | 0.3981 | 0.7371 |
| G0 | 0.6476 | 0.2734 | 0.7050 | 0.6929 | 0.3583 | 0.7533 |
| G1 | <u>0.6536</u> | 0.2974 | **0.7322** | **0.7063** | <u>0.4000</u> | **0.7786** |
| Scratch-N | 0.6056 | 0.2257 | 0.7104 | 0.6558 | 0.3267 | 0.7363 |
| gRNAde | 0.6401 | 0.2481 | 0.6953 | 0.6767 | 0.3234 | 0.7468 |
| NA-MPNN | **0.6556** | <u>0.3131</u> | 0.7109 | 0.6718 | 0.3421 | 0.7367 |
| SMARTPocket local | 0.6397 | **0.3334** | 0.7043 | <u>0.7057</u> | **0.4863** | 0.7643 |

All 49 pairs are pooled within each state: 3,591 apo residues (812 positives) and 3,612 holo residues (813 positives). BA is the mean of pooled sensitivity and specificity; MCC is Matthews correlation coefficient; AUROC is the area under the receiver operating characteristic curve. BA val and MCC val use the same frozen validation-MCC-selected thresholds, without retuning on SMARTFlex49. Unlike main Table 6, these are not equal-target macro metrics. Bold/underline mark the best/second-best distinct unrounded value per column. Model inputs and exposure differ, and repeated structures retain their pair weights; rankings are descriptive.

No PDB or exact-sequence matches were found against retained BRIDGE pretraining train/validation or CoBRA downstream train/validation. However, gRNAde's public candidate inventory overlaps 3/49 pairs and NA-MPNN's design-training PDB list overlaps 14/49. Eight pairs also match BRIDGE's pretraining test inventory. Inventory absence does not establish historical checkpoint non-exposure or family independence. G1 has the highest pooled AUROC in both states, whereas SMARTPocket has the highest validation-threshold MCC. These descriptive rankings do not isolate architecture effects or contradict the differently weighted six-target macro comparison in main Table 6.

### 5 RNAProt head and adaptation panel

CLEAN retains all 191 preprocessed test graphs, removes 20 validation graphs sharing test PDBs, and then 54 training graphs sharing a test or retained-validation PDB. This yields 835/171/191 training/validation/test graphs and 177 test PDB-connected units, using PDB identities without consulting test labels. Table S26 reports GNN metrics including sequence-only; GAT is retained once in Table S31. Native baselines and their validation-selected thresholds are specified in Section 15.

**Table S26. RNA–protein interface prediction with the GNN readout on the RNAProt CLEAN test set (191 graphs, 177 PDB-connected units).**

| Encoder | Mode | AP | AUROC | MCC .5 | MCC val | MCC oracle |
|---|---|---|---|---|---|---|
| **GNN readout** | | | | | | |
| Sequence-only | frozen | 0.6329 | 0.7986 | 0.3943 | 0.4088 | 0.4183 |
| S0 | frozen | 0.6415 | 0.7983 | 0.4101 | 0.4141 | 0.4202 |
| S0 | LoRA | 0.6470 | 0.7976 | <u>0.4194</u> | 0.4072 | 0.4226 |
| S0 | full | 0.6292 | 0.7840 | 0.3881 | 0.3963 | 0.4013 |
| G0 | frozen | 0.6471 | 0.8003 | 0.4102 | 0.4178 | 0.4216 |
| G0 | LoRA | **0.6550** | **0.8039** | **0.4239** | **0.4387** | **0.4402** |
| G0 | full | 0.6415 | 0.7868 | 0.3893 | 0.3987 | 0.4162 |
| G1 | frozen | 0.6328 | 0.7951 | 0.3900 | 0.3900 | 0.4136 |
| G1 | LoRA | 0.6402 | 0.7877 | 0.4099 | 0.4208 | 0.4252 |
| G1 | full | 0.6506 | 0.7978 | 0.4153 | <u>0.4289</u> | <u>0.4293</u> |
| Scratch-N | full | 0.6225 | 0.7762 | 0.3379 | 0.3851 | 0.4020 |
| Random-frozen | frozen | 0.5160 | 0.6728 | 0.2226 | 0.2701 | 0.2917 |
| gRNAde | frozen | 0.5992 | 0.7738 | 0.3473 | 0.3408 | 0.3567 |
| gRNAde | LoRA | 0.6220 | 0.7879 | 0.3504 | 0.3663 | 0.4052 |
| gRNAde | full | 0.6118 | 0.7803 | 0.3645 | 0.3700 | 0.3836 |
| NA-MPNN | frozen | 0.6335 | 0.7794 | 0.3753 | 0.3994 | 0.4238 |
| NA-MPNN | LoRA | 0.6383 | 0.7931 | 0.3895 | 0.3959 | 0.4090 |
| NA-MPNN | full | <u>0.6517</u> | <u>0.8028</u> | 0.3955 | 0.4045 | 0.4096 |

BRIDGE uses pretraining epoch 100/seed 0. Public checkpoints are fixed. Three downstream-seed probability ensembles use 40 epochs and validation-selected checkpoints. Bold and underline mark the best and second-best distinct values among displayed encoders and adaptation modes within the GNN panel for each metric, with each MCC component ranked separately. Emphasis is descriptive. The GAT panel is reported in Supplementary Table S31. Released-encoder inputs and pretraining exposure differ from BRIDGE. Random-frozen trains only the head. Scratch-N full trains the encoder and head from random initialization. Macro AP over all units is undefined because 55 units have no positives. Ties at the displayed precision share emphasis. MCC .5 uses threshold 0.5; val uses the validation-selected threshold; oracle uses test labels to maximize MCC and is a nondeployable diagnostic ceiling. AP, average precision. AUROC, area under the receiver operating characteristic curve; MCC, Matthews correlation coefficient.

#### 5.1 Three downstream seeds for each head/adaptation condition

Source seed 0 is fixed. Each row contains all three individual-run scores and their mean±SD alongside the separately computed probability-ensemble metric.

##### Average precision (AP).

**Table S27. RNAProt individual-seed and probability-ensemble AP.**

| Cohort | Model | Seed 0 | Seed 1 | Seed 2 | Mean ± SD | Ensemble |
|---|---|---|---|---|---|---|
| CLEAN test | G0 / frozen / GNN | 0.6366 | 0.6216 | 0.6504 | 0.6362 ± 0.0144 | 0.6471 |
| CLEAN test | G0 / lora / GNN | 0.6536 | 0.6109 | 0.6322 | 0.6323 ± 0.0213 | 0.6550 |
| CLEAN test | G0 / full / GNN | 0.6247 | 0.6350 | 0.6185 | 0.6260 ± 0.0083 | 0.6415 |
| CLEAN test | G0 / frozen / GAT | 0.6250 | 0.6222 | 0.6133 | 0.6202 ± 0.0061 | 0.6384 |
| CLEAN test | G0 / lora / GAT | 0.6277 | 0.6317 | 0.6493 | 0.6362 ± 0.0115 | 0.6574 |
| CLEAN test | G0 / full / GAT | 0.6206 | 0.6062 | 0.6304 | 0.6191 ± 0.0122 | 0.6397 |
| CLEAN test | G1 / frozen / GNN | 0.6190 | 0.6229 | 0.6258 | 0.6226 ± 0.0034 | 0.6328 |
| CLEAN test | G1 / lora / GNN | 0.6189 | 0.6297 | 0.6281 | 0.6256 ± 0.0059 | 0.6402 |
| CLEAN test | G1 / full / GNN | 0.6374 | 0.6304 | 0.6431 | 0.6370 ± 0.0063 | 0.6506 |
| CLEAN test | G1 / frozen / GAT | 0.6287 | 0.6450 | 0.6362 | 0.6366 ± 0.0081 | 0.6523 |
| CLEAN test | G1 / lora / GAT | 0.6283 | 0.6180 | 0.6294 | 0.6252 ± 0.0063 | 0.6477 |
| CLEAN test | G1 / full / GAT | 0.6284 | 0.6296 | 0.6297 | 0.6292 ± 0.0007 | 0.6412 |
| CLEAN test | Scratch-N / full / GNN | 0.6093 | 0.5852 | 0.3937 | 0.5294 ± 0.1182 | 0.6225 |
| CLEAN test | Scratch-N / full / GAT | 0.5820 | 0.5960 | 0.6105 | 0.5961 ± 0.0142 | 0.6153 |
| CLEAN test | Random-frozen / frozen / GNN | 0.5125 | 0.4991 | 0.5149 | 0.5088 ± 0.0085 | 0.5160 |
| CLEAN test | Random-frozen / frozen / GAT | 0.4825 | 0.4823 | 0.4956 | 0.4868 ± 0.0076 | 0.5036 |

**Table S28. Paired RNAProt head and adaptation AP contrasts.** The archived label Scratch-N / frozen denotes the random-frozen control.

| **Candidate − reference** | **ΔAP [95% CI]** |
|---|---|
| G0 / frozen / gnn – Scratch-N / frozen / gnn | 0.1312 [0.0733, 0.2001] |
| G0 / lora / gnn – G0 / frozen / gnn | 0.0079 [-0.0104, 0.0252] |
| G0 / lora / gnn – Scratch-N / full / gnn | 0.0325 [0.0112, 0.0557] |
| G0 / full / gnn – G0 / frozen / gnn | -0.0057 [-0.0279, 0.0145] |
| G0 / full / gnn – Scratch-N / full / gnn | 0.0190 [0.0028, 0.0363] |
| G0 / frozen / gat – G0 / frozen / gnn | -0.0087 [-0.0233, 0.0034] |
| G0 / frozen / gat – Scratch-N / frozen / gat | 0.1348 [0.0763, 0.2016] |
| G0 / lora / gat – G0 / lora / gnn | 0.0024 [-0.0125, 0.0173] |
| G0 / lora / gat – G0 / frozen / gat | 0.0190 [0.0004, 0.0388] |
| G0 / lora / gat – Scratch-N / full / gat | 0.0421 [0.0225, 0.0626] |
| G0 / full / gat – G0 / full / gnn | -0.0018 [-0.0183, 0.0152] |
| G0 / full / gat – G0 / frozen / gat | 0.0012 [-0.0306, 0.0311] |
| G0 / full / gat – Scratch-N / full / gat | 0.0244 [-0.0006, 0.0495] |
| G1 / frozen / gnn – Scratch-N / frozen / gnn | 0.1169 [0.0571, 0.1900] |
| G1 / lora / gnn – G1 / frozen / gnn | 0.0074 [-0.0123, 0.0260] |
| G1 / lora / gnn – Scratch-N / full / gnn | 0.0177 [-0.0009, 0.0387] |
| G1 / full / gnn – G1 / frozen / gnn | 0.0178 [-0.0045, 0.0371] |
| G1 / full / gnn – Scratch-N / full / gnn | 0.0281 [0.0124, 0.0455] |
| G1 / frozen / gat – G1 / frozen / gnn | 0.0195 [0.0044, 0.0324] |
| G1 / frozen / gat – Scratch-N / frozen / gat | 0.1487 [0.0915, 0.2142] |
| G1 / lora / gat – G1 / lora / gnn | 0.0075 [-0.0085, 0.0242] |
| G1 / lora / gat – G1 / frozen / gat | -0.0046 [-0.0229, 0.0153] |
| G1 / lora / gat – Scratch-N / full / gat | 0.0324 [0.0123, 0.0536] |
| G1 / full / gat – G1 / full / gnn | -0.0094 [-0.0225, 0.0050] |
| G1 / full / gat – G1 / frozen / gat | -0.0111 [-0.0343, 0.0133] |
| G1 / full / gat – Scratch-N / full / gat | 0.0259 [0.0059, 0.0479] |
| Scratch-N / frozen / gat – Scratch-N / frozen / gnn | -0.0123 [-0.0347, 0.0085] |
| Scratch-N / full / gat – Scratch-N / full / gnn | -0.0072 [-0.0274, 0.0138] |

**Table S28. (continued)**

#### 5.2 Objective ablation with paired pretraining/downstream seeds

The individual AP values and sample SDs in Supplementary Data D035 are from three paired source/downstream runs in the completed archived analysis. Ensemble AP is computed from mean probabilities, not the mean of these three APs. The random initialization also varies across seeds for Scratch-N in this panel.

**Supplementary Data D035.** RNAProt objective-ablation AP across paired pretraining and downstream seeds.

**Table S29. Probability-ensemble metrics for the paired-source-seed RNAProt objective ablation.**

| **Model** | **AP** | **Macro AP** | **AUROC** | **MCC .5 / val / oracle** |
|---|---|---|---|---|
| S0 | 0.6407 | – | 0.7902 | <u>0.3854</u> / **0.4229** / **0.4327** |
| S0+PairGeom | <u>0.6437</u> | – | <u>0.7954</u> | **0.3983** / <u>0.4068</u> / 0.4204 |
| S0+$\chi$ | **0.6448** | – | **0.7983** | 0.3787 / 0.4048 / <u>0.4263</u> |
| G0 | 0.6272 | – | 0.7854 | 0.3650 / 0.3868 / 0.4088 |
| G1 | 0.6305 | – | 0.7833 | 0.3830 / 0.4034 / 0.4187 |
| Scratch-N | 0.5970 | – | 0.7708 | 0.3594 / 0.3655 / 0.3736 |

This objective-removal panel varies paired pretraining/downstream seeds, whereas the main head/adaptation panel fixes pretraining seed 0. Removing $\chi$ or PairGeom changes AP by +0.0165 [0.0020, 0.0332] or +0.0176 [0.0037, 0.0315] relative to G0. Cross-panel differences do not isolate head effects.

**Table S30. RNA–protein interface prediction in published and local protocols.**

| Method | AP | AUROC | Accuracy | MCC reported | MCC .5 | MCC oracle |
|---|---|---|---|---|---|---|
| **Published protocols** | | | | | | |
| RGCN [11] | – | 0.6654 | – | 0.2469 | – | – |
| DualEquiNet [12] | – | – | 0.749 ± 0.015 | – | – | – |
| **Local CLEAN test: 191 graphs** | | | | | | |
| RGCN / native | 0.4944 | 0.7241 | 0.7003 | – | 0.3160 | 0.3323 |
| DualEquiNet / native | 0.4288 | 0.6771 | 0.7229 | – | 0.0239 | 0.2665 |
| G0 / frozen / GNN | 0.6471 | 0.8003 | 0.7455 | – | 0.4102 | 0.4216 |
| G0 / LoRA / GNN | **0.6550** | **0.8039** | **0.7484** | – | **0.4239** | **0.4402** |
| G0 / full / GNN | 0.6415 | 0.7868 | 0.7294 | – | 0.3893 | 0.4162 |

Published RGCN values are from Table 7 of the RNAglib benchmark [11]. Published DualEquiNet accuracy is mean±SD from Table 6 of [12]. Their original partitions and aggregation differ from CLEAN. The local block compares identical test membership and labels using three-seed probability ensembles. Native RGCN/DualEquiNet rows are our 40-epoch reruns. G0 rows repeat the GNN panel from Table S26, with accuracy added. Native inputs, architectures and optimization are not matched to BRIDGE. Local MCC .5 and oracle use a fixed 0.5 threshold and a test-label oracle threshold, respectively (a nondeployable diagnostic ceiling), whereas published MCC uses an unspecified threshold. Local accuracy is pooled residue accuracy at threshold 0.5, computed from the same three-seed mean probabilities used for AP, AUROC and MCC. Dashes indicate unavailable or undisplayed metrics. Bold and underline mark the best and second-best distinct displayed estimates within each protocol block and metric, respectively; ties share emphasis. Columns with fewer than two estimates are not ranked. No ranking across protocol blocks or statistical superiority is implied. Validation-threshold MCC and all seed results are supplementary. AP, average precision. AUROC, area under the receiver operating characteristic curve; MCC, Matthews correlation coefficient.

#### 5.3 Published RNAProt metrics

**Supplementary Data D038.** Published RNAProt benchmark metrics.

Wyss et al. [11] Table 7 reports RGCN AUROC 0.6654, balanced accuracy 0.6254 and MCC 0.2469, whereas Table 1 reports balanced accuracy 0.553±0.007. Their aggregation/run relationship is unreconciled. Both are retained. The published 881/189/189 split differs from local 835/171/191. Its MCC threshold is unspecified and cannot be inserted as fixed/validation/oracle MCC.

Xu et al. [12] Appendix D.1 Table 6 reports accuracy for SchNet, EGNN, GVP-GNN, TFN, FastEGNN, HEGNN, GotenNet and DualEquiNet. The pinned released runner thresholds at 0.5 and averages batch accuracies equally.

The loader defaults to random 70/15/15 splits without our PDB exclusion. These code observations do not prove identical training settings for every published baseline. AP is not reported by that table. The same-name sequence-window RNAProt method is a different endpoint and is not included.

#### 5.4 RNAProt GAT readout

**Table S31.** RNA–protein interface prediction with the GAT readout on RNAProt CLEAN (191 test graphs, 177 PDB-connected units). This panel complements the GNN results in main-text Table 5. BRIDGE uses pretraining epoch 100/seed 0; released public checkpoints are fixed. Values use three downstream-seed probability ensembles after 40 epochs with validation-selected checkpoints. Bold and underline indicate the best and second-best distinct estimates within each metric in this GAT panel. Emphasis is descriptive. MCC .5 uses threshold 0.5; MCC val uses a validation-selected threshold; MCC oracle maximizes MCC using test labels and is a nondeployable diagnostic ceiling. Strict macro AP is undefined because 55 units contain no positives. Input representations and pretraining exposure differ across released encoders. Random-frozen trains only the readout; Scratch-N full trains both encoder and readout from random initialization.

| Encoder | Mode | AP | AUROC | MCC .5 | MCC val | MCC oracle |
|---|---|---|---|---|---|---|
| S0 | frozen | 0.6500 | <u>0.8084</u> | <u>0.4270</u> | 0.4209 | 0.4296 |
| S0 | LoRA | 0.6416 | 0.7981 | 0.4200 | 0.4182 | 0.4237 |
| S0 | full | 0.6525 | 0.7995 | **0.4284** | **0.4429** | **0.4463** |
| G0 | frozen | 0.6384 | 0.8006 | 0.4041 | 0.4090 | 0.4147 |
| G0 | LoRA | **0.6574** | 0.7955 | 0.4255 | 0.4284 | <u>0.4437</u> |
| G0 | full | 0.6397 | 0.7930 | 0.4025 | 0.4150 | 0.4238 |
| G1 | frozen | 0.6523 | 0.8017 | 0.4096 | 0.4005 | 0.4183 |
| G1 | LoRA | 0.6477 | 0.8019 | 0.4075 | <u>0.4397</u> | 0.4434 |
| G1 | full | 0.6412 | 0.7875 | 0.3804 | 0.4140 | 0.4221 |
| Scratch-N | full | 0.6153 | 0.7766 | 0.3737 | 0.3912 | 0.4029 |
| Random-frozen | frozen | 0.5036 | 0.6871 | 0.2184 | 0.2594 | 0.2666 |
| gRNAde | frozen | 0.5812 | 0.7697 | 0.3383 | 0.3372 | 0.3433 |
| gRNAde | LoRA | 0.6292 | 0.7893 | 0.3530 | 0.3851 | 0.4060 |
| gRNAde | full | 0.6268 | 0.7975 | 0.3693 | 0.3769 | 0.3875 |
| NA-MPNN | frozen | 0.6432 | 0.7897 | 0.3969 | 0.3990 | 0.4197 |
| NA-MPNN | LoRA | <u>0.6532</u> | **0.8088** | 0.4067 | 0.4067 | 0.4267 |
| NA-MPNN | full | 0.6436 | 0.8002 | 0.3996 | 0.3898 | 0.4060 |

### 6 ATOM3D RNA structure ranking

Tables S32 and S33 give the MLP and width32 GNN counterparts to Figure 2E/F. Table S34 reports the larger MLP83k/residual-GNN83k readouts; Table S70 includes both small GNN widths. All adaptation conditions are shown.

**Table S32. ATOM3D RNA structure ranking (RSR) with the MLP readout at 20 epochs.**

| Encoder | Mode | Target $\rho$ | Seed SD | Pooled $\rho$ |
|---|---|---|---|---|
| **MLP readout** | | | | |
| S0 | frozen | 0.3106 | 0.0436 | 0.5059 |
| S0 | LoRA | 0.4333 | 0.0209 | 0.5094 |
| S0 | full | 0.3694 | 0.0787 | 0.3836 |
| G0 | frozen | 0.3107 | 0.0146 | <u>0.6046</u> |
| G0 | LoRA | <u>0.5107</u> | 0.0670 | 0.5898 |
| G0 | full | 0.4270 | 0.0452 | 0.5726 |
| G1 | frozen | 0.3470 | 0.0218 | **0.6173** |
| G1 | LoRA | **0.5377** | 0.0109 | 0.5539 |
| G1 | full | 0.3975 | 0.0977 | 0.4300 |
| Scratch-N | full | 0.2737 | 0.0090 | 0.3397 |
| Random-frozen | frozen | 0.2249 | 0.0445 | 0.5134 |
| gRNAde | frozen | 0.1953 | 0.0332 | 0.3604 |
| gRNAde | LoRA | 0.2369 | 0.0514 | 0.0483 |
| gRNAde | full | 0.2413 | 0.1142 | 0.2289 |
| NA-MPNN | frozen | 0.2646 | 0.0492 | 0.4629 |
| NA-MPNN | LoRA | 0.3201 | 0.0778 | 0.4522 |
| NA-MPNN | full | 0.3170 | 0.0418 | 0.4821 |
| **Released atomic reference** | | | | |
| ARES [9] | published | 0.3739 | – | 0.4145 |

The MLP readout has 16,641 parameters and includes S0/G0/G1 in three adaptation modes, random-frozen and scratch-full controls, and gRNAde/NA-MPNN in three modes. The table contains 17 fitted conditions and one released ARES-score reference. Values average three seed-specific metrics, with native pretraining seed 0 and public checkpoints fixed. Mean-target correlation weights four puzzles equally. Bold and underline mark the best and second-best distinct values among all displayed rows in each correlation column. Emphasis is descriptive, and SD is not ranked. Released-encoder inputs and pretraining exposure differ from BRIDGE. The width32 GNN panel is reported in Supplementary Table S33. Larger heads, per-seed and per-target results, and uncertainty are also in the supplement. Ties at the displayed precision share emphasis. $\rho$, Spearman correlation; SD, standard deviation.

#### 6.1 RSR width32 GNN readout

**Table S33.** ATOM3D RNA structure ranking with the residual width32 GNN readout at 20 epochs. This panel complements the MLP results in Supplementary Table S32 and main Figure 2F. Values average three seed-specific metrics, with native pretraining seed 0 and public checkpoints fixed. Mean-target Spearman correlation weights the four test puzzles equally. The GNN has 8,513 parameters. Bold and underline indicate the best and second-best distinct estimates within each correlation column; SD is not ranked. Ranking is descriptive and does not establish statistical superiority. Released-encoder inputs and pretraining exposure differ from BRIDGE. Figure 2 retains both readouts for the adaptation comparison.

| Encoder | Mode | Target $\rho$ | Seed SD | Pooled $\rho$ |
|---|---|---|---|---|
| S0 | frozen | <u>0.4189</u> | 0.0370 | 0.5045 |
| S0 | LoRA | 0.3203 | 0.0370 | 0.4296 |
| S0 | full | 0.3373 | 0.2332 | 0.4360 |
| G0 | frozen | 0.3712 | 0.0649 | 0.4542 |
| G0 | LoRA | 0.3520 | 0.0412 | 0.5329 |
| G0 | full | 0.3998 | 0.0465 | 0.5447 |
| G1 | frozen | 0.3947 | 0.0835 | 0.5298 |
| G1 | LoRA | **0.4729** | 0.0505 | **0.5772** |
| G1 | full | 0.3564 | 0.0391 | 0.5283 |
| Scratch-N | full | 0.1855 | 0.1331 | 0.4862 |
| Random-frozen | frozen | 0.4003 | 0.0237 | <u>0.5528</u> |
| gRNAde | frozen | 0.2617 | 0.1506 | 0.2921 |
| gRNAde | LoRA | 0.2097 | 0.0549 | -0.0768 |
| gRNAde | full | -0.0651 | 0.1691 | -0.0356 |
| NA-MPNN | frozen | 0.2305 | 0.0439 | 0.3865 |
| NA-MPNN | LoRA | 0.3523 | 0.0929 | 0.5249 |
| NA-MPNN | full | 0.3130 | 0.1269 | 0.4499 |

**Table S34. ATOM3D RSR: the retained larger MLP and residual-GNN readouts.**

| Encoder | Mode | Head | Target $\rho$ | Seed SD | Pooled $\rho$ |
|---|---|---|---|---|---|
| S0 | frozen | MLP83k | 0.3882 | 0.0294 | 0.5536 |
| S0 | LoRA | MLP83k | **0.4799** | 0.0184 | 0.6169 |
| S0 | full | MLP83k | 0.4335 | 0.1267 | 0.4567 |
| G0 | frozen | MLP83k | 0.3499 | 0.0126 | 0.5954 |
| G0 | LoRA | MLP83k | 0.4706 | 0.0605 | 0.5798 |
| G0 | full | MLP83k | 0.3687 | 0.0482 | 0.4824 |
| G1 | frozen | MLP83k | 0.3919 | 0.0095 | 0.6310 |
| G1 | LoRA | MLP83k | 0.4654 | 0.0315 | 0.6151 |
| G1 | full | MLP83k | 0.4406 | 0.0542 | 0.5765 |
| Scratch-N | full | MLP83k | 0.2703 | 0.0659 | 0.4791 |
| Random-frozen | frozen | MLP83k | 0.3256 | 0.0280 | **0.6316** |
| S0 | frozen | ResidualGNN83k | 0.3789 | 0.0130 | 0.5176 |
| S0 | LoRA | ResidualGNN83k | 0.3145 | 0.0417 | 0.3829 |
| S0 | full | ResidualGNN83k | 0.3622 | 0.0086 | 0.5155 |
| G0 | frozen | ResidualGNN83k | 0.3760 | 0.0332 | 0.4686 |
| G0 | LoRA | ResidualGNN83k | 0.3510 | 0.0481 | 0.5146 |
| G0 | full | ResidualGNN83k | 0.3243 | 0.0486 | 0.4599 |
| G1 | frozen | ResidualGNN83k | 0.3141 | 0.0324 | 0.4797 |
| G1 | LoRA | ResidualGNN83k | **0.4026** | 0.0467 | **0.5323** |
| G1 | full | ResidualGNN83k | 0.3608 | 0.0798 | 0.4598 |
| Scratch-N | full | ResidualGNN83k | 0.0955 | 0.0383 | 0.4497 |
| Random-frozen | frozen | ResidualGNN83k | 0.3808 | 0.0247 | 0.4822 |

All 22 original larger-head conditions are retained. Values average three separately scored downstream fits; SD is fitting variability, not target-sampling uncertainty. Ranks compare all displayed modes within each head and correlation column. No condition was selected for this table by its test score.

Readout and endpoint choice affect model ordering. Random-frozen MLP83k reaches pooled correlation 0.6316 but target-macro correlation 0.3256; random-frozen residual GNN83k reaches target-macro correlation 0.3808 versus G0/frozen 0.3760. S0 LoRA reaches 0.4333 with MLP and 0.4799 with MLP83k. G1 LoRA reaches 0.4729/0.3708 with GNN width32/64, compared with 0.5377 with MLP. These four-puzzle results support readout-dependent performance rather than a universally superior encoder or head.

#### 6.2 Per-puzzle adaptation effects

**Table S35. Per-puzzle frozen-to-LoRA effects on ATOM3D RNA structure ranking.**

| Encoder | Puzzle | Frozen $\rho$ | LoRA $\rho$ | $\Delta\,\rho$ |
|---|---|---|---|---|
| G0 | 18 | -0.1378 | 0.2962 | 0.4340 |
| G0 | 19 | 0.5437 | 0.6229 | 0.0792 |
| G0 | 20 | 0.5538 | 0.6629 | 0.1091 |
| G0 | 21 | 0.2832 | 0.4607 | 0.1776 |
| G1 | 18 | -0.0779 | 0.3824 | 0.4603 |
| G1 | 19 | 0.5389 | 0.6440 | 0.1050 |
| G1 | 20 | 0.6141 | 0.6042 | -0.0099 |
| G1 | 21 | 0.3128 | 0.5202 | 0.2074 |

Per-puzzle differences average seed-specific correlations before equal-puzzle aggregation. G1's Puzzle-20 effect is negative despite a positive aggregate conditional interval. Random-frozen MLP83k leads pooled correlation (0.6316) and RMSE (4.4250), illustrating the distinction between pooled and target-macro endpoints.

**Supplementary Data D043.** ATOM3D readout and adaptation results, including RMSD error.

Aggregate, individual-seed and per-puzzle results are reported separately; seed-specific metrics are averaged without ensembling predicted RMSD.

**Supplementary Data D044.** Individual-seed ATOM3D ranking results and selected epochs.

**Supplementary Data D045.** Per-puzzle ATOM3D correlations and top-1 regret.

**Table S36. Paired ATOM3D readout and adaptation contrasts.** Archived labels mlp17k, mlp and resgnn denote the 16,641-parameter MLP, MLP83k and ResidualGNN83k, respectively; Scratch-N / frozen denotes the random-frozen control.

| **Candidate − reference** | **Δmean-target $\rho$ [95% CI]** |
|---|---|
| G0-N_epoch100__frozen__mlp17k − Scratch-N__frozen__mlp17k | 0.0858 [-0.0116, 0.1617] |
| G0-N_epoch100__lora__mlp17k − G0-N_epoch100__frozen__mlp17k | 0.2000 [0.0941, 0.3528] |
| G0-N_epoch100__full__mlp17k − G0-N_epoch100__frozen__mlp17k | 0.1162 [-0.0077, 0.2402] |
| G0-N_epoch100__full__mlp17k − Scratch-N__full__mlp17k | 0.1533 [-0.0611, 0.3935] |
| G0-N_epoch100__frozen__mlp − Scratch-N__frozen__mlp | 0.0243 [-0.0880, 0.1366] |
| G0-N_epoch100__frozen__mlp − G0-N_epoch100__frozen__mlp17k | 0.0392 [0.0198, 0.0585] |
| G0-N_epoch100__lora__mlp − G0-N_epoch100__frozen__mlp | 0.1207 [0.0466, 0.2297] |
| G0-N_epoch100__lora__mlp − G0-N_epoch100__lora__mlp17k | -0.0401 [-0.0898, -0.0127] |
| G0-N_epoch100__full__mlp − G0-N_epoch100__frozen__mlp | 0.0188 [-0.0488, 0.1004] |
| G0-N_epoch100__full__mlp − Scratch-N__full__mlp | 0.0984 [-0.1613, 0.4282] |
| G0-N_epoch100__full__mlp − G0-N_epoch100__full__mlp17k | -0.0583 [-0.1135, 0.0015] |
| G0-N_epoch100__frozen__resgnn − Scratch-N__frozen__resgnn | -0.0048 [-0.1051, 0.1137] |
| G0-N_epoch100__frozen__resgnn − G0-N_epoch100__frozen__mlp | 0.0261 [-0.1014, 0.1977] |
| G0-N_epoch100__lora__resgnn − G0-N_epoch100__frozen__resgnn | -0.0249 [-0.1329, 0.0822] |
| G0-N_epoch100__lora__resgnn − G0-N_epoch100__lora__mlp | -0.1195 [-0.1887, -0.0702] |
| G0-N_epoch100__full__resgnn − G0-N_epoch100__frozen__resgnn | -0.0517 [-0.2486, 0.1859] |
| G0-N_epoch100__full__resgnn − Scratch-N__full__resgnn | 0.2288 [0.1030, 0.4124] |
| G0-N_epoch100__full__resgnn − G0-N_epoch100__full__mlp | -0.0444 [-0.1872, 0.0984] |
| G1-N_epoch100__frozen__mlp17k − Scratch-N__frozen__mlp17k | 0.1221 [0.0372, 0.1741] |
| G1-N_epoch100__lora__mlp17k − G1-N_epoch100__frozen__mlp17k | 0.1907 [0.0444, 0.3715] |
| G1-N_epoch100__full__mlp17k − G1-N_epoch100__frozen__mlp17k | 0.0506 [-0.0916, 0.1781] |
| G1-N_epoch100__full__mlp17k − Scratch-N__full__mlp17k | 0.1238 [-0.0784, 0.4372] |
| G1-N_epoch100__frozen__mlp − Scratch-N__frozen__mlp | 0.0663 [-0.0193, 0.1519] |
| G1-N_epoch100__frozen__mlp − G1-N_epoch100__frozen__mlp17k | 0.0449 [-0.0014, 0.0912] |
| G1-N_epoch100__lora__mlp − G1-N_epoch100__frozen__mlp | 0.0735 [-0.0068, 0.1538] |
| G1-N_epoch100__lora__mlp − G1-N_epoch100__lora__mlp17k | -0.0723 [-0.1577, 0.0131] |
| G1-N_epoch100__full__mlp − G1-N_epoch100__frozen__mlp | 0.0487 [-0.0381, 0.1353] |
| G1-N_epoch100__full__mlp − Scratch-N__full__mlp | 0.1703 [-0.0190, 0.4598] |
| G1-N_epoch100__full__mlp − G1-N_epoch100__full__mlp17k | 0.0431 [-0.0600, 0.1409] |
| G1-N_epoch100__frozen__resgnn − Scratch-N__frozen__resgnn | -0.0667 [-0.1914, 0.0580] |
| G1-N_epoch100__frozen__resgnn − G1-N_epoch100__frozen__mlp | -0.0778 [-0.2017, 0.0367] |
| G1-N_epoch100__lora__resgnn − G1-N_epoch100__frozen__resgnn | 0.0885 [-0.0225, 0.2146] |
| G1-N_epoch100__lora__resgnn − G1-N_epoch100__lora__mlp | -0.0628 [-0.1494, 0.0057] |
| G1-N_epoch100__full__resgnn − G1-N_epoch100__frozen__resgnn | 0.0468 [-0.1373, 0.2441] |
| G1-N_epoch100__full__resgnn − Scratch-N__full__resgnn | 0.2653 [0.1565, 0.4198] |
| G1-N_epoch100__full__resgnn − G1-N_epoch100__full__mlp | -0.0798 [-0.1418, -0.0264] |
| Scratch-N__frozen__mlp − Scratch-N__frozen__mlp17k | 0.1007 [0.0481, 0.1822] |
| Scratch-N__full__mlp − Scratch-N__full__mlp17k | -0.0034 [-0.0958, 0.0891] |
| Scratch-N__frozen__resgnn − Scratch-N__frozen__mlp | 0.0552 [0.0006, 0.1097] |
| Scratch-N__full__resgnn − Scratch-N__full__mlp | -0.1748 [-0.4914, 0.1683] |

#### 6.3 Separate frozen-readout loss ablation

The frozen-readout ablation uses the test-excluded source epoch-100 encoders and the same 12,479/4,000/4,000 decoy split (test Puzzles 18–21), but is not part of the preceding 20-epoch adaptation experiment. Encoders are frozen. Three independently pretrained source seeds (0/1/2) are each paired with the same readout seed 0. A mean-pooled 128-dimensional embedding feeds a 128–128–1 ReLU MLP with dropout 0.1. Training-only embedding/RMSD standardization, MSE, Adam (learning rate 0.001, weight decay 0.0001), batch size 256 and 100 head epochs are used. The earliest validation target-macro Spearman maximum selects the checkpoint. Values average separately scored runs, not a score ensemble. Source SD is the sample SD across source seeds, not downstream fitting variability. Correlation ranks below are confined to these five same-contract models.

**Table S37. Frozen-readout ATOM3D objective-ablation results.**

| **Encoder** | **Mean-target $\rho$** | **Source SD** | **Pooled $\rho$** |
|---|---|---|---|
| S0 | 0.1979 | 0.0446 | 0.3972 |
| S0+PairGeom | <u>0.2691</u> | 0.1206 | <u>0.4568</u> |
| S0+$\chi$ | 0.2426 | 0.0668 | 0.4477 |
| G0 | 0.2648 | 0.1099 | 0.4566 |
| G1 | **0.2977** | 0.0609 | **0.5143** |

These five frozen-readout models share the 100-epoch head-fitting protocol. Removing $\chi$ gives a similar target-macro estimate, while removing both auxiliary objectives lowers it descriptively. This panel does not measure effects under LoRA/full adaptation. Paired intervals resample four puzzles and condition on fitted models; they do not include pretraining variability.

**Supplementary Data D048.** Individual-pretraining-seed results for the frozen ATOM3D objective ablation.

**Supplementary Data D049.** Per-puzzle correlations for the frozen ATOM3D objective ablation.

**Table S38. Paired contrasts for the frozen ATOM3D objective ablation.**

| Contrast | Δmean-target $\rho$ | Conditional 95% CI |
|---|---|---|
| S0 minus G0 | -0.0669 | [-0.1340, -0.0007] |
| S0+PairGeom minus G0 | 0.0043 | [-0.0919, 0.1005] |
| S0+$\chi$ minus G0 | -0.0223 | [-0.0610, 0.0068] |
| G1 minus G0 | 0.0329 | [0.0142, 0.0515] |

##### 6.4 ARES released-score comparison on the current RSR test

This panel uses released ARES [9] benchmark1.csv predictions (Zenodo record 5090157) without rerunning ARES or changing BRIDGE models. Puzzle/tag matching aligns all 4,000 ordinary test decoys (1,000 per Puzzle 18–21), with RMSD and Rosetta-energy agreement within $10^{-6}$; native and near-native examples outside this test are excluded. ARES mean-target/pooled Spearman is 0.3739/0.4145. Main Figure 2 instead uses a separate CPU replay of the released checkpoint and native transform (mean-target Spearman 0.373846); Table S32 retains the archived-score reference. ARES supplies one fitted model, whereas BRIDGE averages three downstream-fit correlations. Evaluation examples and RMSD endpoints are aligned, but atomic inputs and training histories are not matched.

**Table S39. Per-puzzle MLP and released ARES correlations.** Scratch-N / frozen denotes the random-frozen control.

| Encoder | Mode | Puzzle 18 | 19 | 20 | 21 |
|---|---|---|---|---|---|
| G0 | frozen | -0.1378 | 0.5437 | 0.5538 | 0.2832 |
| G0 | LoRA | 0.2962 | 0.6229 | 0.6629 | 0.4607 |
| G0 | full | 0.1458 | 0.4953 | 0.5869 | 0.4799 |
| G1 | frozen | -0.0779 | 0.5389 | 0.6141 | 0.3128 |
| G1 | LoRA | 0.3824 | 0.6440 | 0.6042 | 0.5202 |
| G1 | full | 0.1001 | 0.3575 | 0.6415 | 0.4911 |
| Scratch-N | frozen | -0.0719 | 0.3720 | 0.4679 | 0.1316 |
| Scratch-N | full | 0.2182 | 0.3012 | 0.0456 | 0.5297 |
| ARES | published | 0.2033 | 0.4526 | 0.6899 | 0.1497 |

BRIDGE-minus-ARES contrasts use equal-puzzle Spearman and 10,000 paired four-puzzle bootstrap draws (seed 1729). BRIDGE seed-specific correlations are averaged before resampling. Supplementary Data D052 retains all 24 conditions; intervals are conditional, unadjusted and limited by four independent targets.

**Supplementary Data D052.** All ATOM3D readout/adaptation contrasts against released ARES scores.

##### 6.5 Published RSR context

**Supplementary Data D053.** Published ATOM3D RNA structure-ranking results.

Published 3DCNN/GNN/Rosetta values follow the revised ATOM3D Tables 6 and 8 [10]; GVP-GNN and EQGAT follow their original reports [5, 7]. Inputs and training histories differ. Published replicate SDs are distinct from paired-target confidence intervals.

#### 7 RNA3D-Decoy adaptation and CASP16 evaluation

##### 7.1 Cohorts, fitting and quality labels

RNA3D Level-I inputs are single-chain predictions from AF3, Boltz, Chai, ESMFold2 and Protenix [2]. Eight CASP-overlapping targets were excluded from RNA3D training before applying a 500-nt maximum length without cropping. AF3 decoys were excluded from training and training-derived normalization, while validation/test membership was retained. Training/validation contain 329/40 targets and 40,138/6,120 decoys. The original test contains 42 targets/6,426 decoys; CASP16 contains 20 targets/3,789 submissions.

Each BRIDGE model in the original MLP panel starts from the retained source-epoch-100/seed-0 encoder, or the corresponding random encoder for Scratch-N, rather than ATOM3D-adapted weights. Mean-pooled 128-dimensional embeddings enter a 128–128–1 ReLU MLP with dropout 0.1 (16,641 parameters). Embeddings and RMSD are standardized using initial training-only sample means/SDs, held fixed thereafter. The objective is decoy-weighted MSE. FP32 Adam uses head rate 0.001, full encoder rate 0.0001, weight decay 0.0001, clipping at 1, batch size 16 and node caps 500/sample and 1,500/microbatch. LoRA has rank 4, alpha 8 and adapter/head rate 0.001. Frozen fits train only the head. The 21 fits all complete 20 epochs. Earliest defined validation target-macro Spearman maxima select checkpoints. Downstream seeds vary the head, ordering and dropout. Source/random-encoder initialization remains fixed. All three original MLP scratch-full runs selected epoch 1. Equal-budget performance is not a scratch-optimal hyperparameter search.

We recomputed labels rather than mixing provider RMSD conventions. Canonical sequence positions register standard backbone and base heavy atoms. Hydrogen/deuterium are excluded. Alternate conformers use occupancy-based deterministic admission. The atom intersection across admitted native conformers is fixed before scoring decoys. Proper rigid-body least-squares alignment is performed on these common atoms, and the smallest RMSD across native conformers is the label. This common-heavy endpoint can omit experimentally unresolved atoms and is not an all-possible-atom or native-completeness claim. Predicted coordinates are the only structural input to inference. Native structures are used solely for labels.

#### 7.2 Post-hoc exposure sensitivity and metric definitions

The raw 42-target results were inspected before these exclusions. The exposure-only 40-target analysis removes 1SCL (exact source-training sequence) and 2Y95 (PDB/exact source-training sequence). RNA3D38 additionally removes 8VCI (known Rfam RF00507 link) and 2IXZ (8 nt), retaining all 153 decoys per remaining target. Thirty-two retained targets have unknown family annotation. This is a named, post-hoc sensitivity, not a general family/length filter, new untouched test or homology-independence certification. The three memberships and all seven conditions are reported below. Checkpoints were not reselected. CASP16 membership is unchanged and its registered sequence–structure clusters do not certify family independence.

For seed $s$, predicted RMSD $\hat{r}_{sti}$ and reference RMSD $r_{ti}$, the two correlation estimands are

$$\rho_{\text{target}} = \frac{1}{3}\sum_{s=1}^{3}\frac{1}{T}\sum_{t=1}^{T}\text{Spearman}(\hat{r}_{st\cdot}, r_{t\cdot}), \tag{7}$$

$$\rho_{\text{pooled}} = \frac{1}{3}\sum_{s=1}^{3}\text{Spearman}(\text{concat}_t\,\hat{r}_{st\cdot}, \text{concat}_t\,r_{t\cdot}). \tag{8}$$

Ties use average ranks. Any undefined target correlation makes the strict macro undefined. No target is dropped or assigned zero. All selected held-out scores have defined correlations. Top-1 regret is the true RMSD of the lowest-scored model minus the lowest true RMSD available for that target, in Å. Score ties choose the first immutable input row. Seed SD uses the sample denominator 2 and is not a confidence interval. Paired 10,000-draw intervals resample 38 RNA3D targets or 10 archived CASP clusters, preserving equal-target point weighting and averaging seed metrics within draws. They are unadjusted, conditional on fitted models and do not include source-training or retrospective-selection uncertainty.

**Table S40.** RNA3D-trained quality predictors on RNA3D38 and CASP16. Mean-target Spearman is mean±sample SD of three downstream-seed metrics. Pooled Spearman also averages seed-specific metrics, not predictions. BRIDGE source epoch 100/seed 0 and each public released checkpoint are fixed. All conditions use MLP and 20 epochs. RNA3D38 is post-hoc. CASP16 comprises 20 targets in 10 registered clusters. Bold/underline rank all displayed models and adaptation modes per cohort and metric, including gRNAde and NA-MPNN. Inputs and pretraining exposure still differ. Seed SD is not ranked. Emphasis is descriptive, not significance or selected deployment models.

| **Model/mode** | **RNA3D38** $\rho_{\text{target}}$ | $\rho_{\text{pooled}}$ | **CASP16** $\rho_{\text{target}}$ | $\rho_{\text{pooled}}$ |
|---|---|---|---|---|
| S0/frozen | 0.2424 ± 0.0520 | 0.5537 | 0.2139 ± 0.0084 | 0.4997 |
| S0/LoRA | 0.3639 ± 0.0362 | **0.6638** | 0.2025 ± 0.0572 | 0.5093 |
| S0/full | 0.2932 ± 0.0568 | 0.5959 | 0.2524 ± 0.0271 | 0.5727 |
| G0/Frozen | 0.2902 ± 0.0405 | 0.5665 | 0.1185 ± 0.0148 | 0.4017 |
| G0/LoRA | 0.3772 ± 0.0297 | 0.6255 | 0.2046 ± 0.0015 | 0.5363 |
| G0/Full | 0.3120 ± 0.0535 | 0.6148 | <u>0.2857</u> ± 0.0158 | 0.5450 |
| G1/Frozen | 0.2646 ± 0.0294 | 0.5531 | 0.1207 ± 0.0088 | 0.3927 |
| G1/LoRA | <u>0.3892</u> ± 0.0858 | 0.6366 | 0.1785 ± 0.0220 | 0.5156 |
| G1/Full | 0.2973 ± 0.0167 | 0.5949 | 0.2420 ± 0.0720 | 0.5019 |
| Scratch-N/Full | 0.2103 ± 0.0236 | 0.5839 | 0.0310 ± 0.0294 | 0.3395 |
| gRNAde/frozen | 0.2662 ± 0.0250 | 0.5961 | 0.2806 ± 0.0188 | **0.6000** |
| gRNAde/LoRA | 0.3533 ± 0.0243 | 0.6339 | 0.1745 ± 0.1081 | 0.5846 |
| gRNAde/full | 0.3109 ± 0.0678 | 0.6168 | 0.1604 ± 0.0434 | 0.5870 |
| NA-MPNN/frozen | 0.3381 ± 0.0606 | 0.5190 | 0.2638 ± 0.0026 | <u>0.5928</u> |
| NA-MPNN/LoRA | **0.4063** ± 0.0434 | 0.6558 | **0.2897** ± 0.0140 | 0.5686 |
| NA-MPNN/full | 0.3771 ± 0.0043 | <u>0.6625</u> | 0.2699 ± 0.0712 | 0.4824 |

**Table S41.** RNA3D target-macro Spearman across the original and post-hoc cohorts.

| Model/mode | Raw42 | Exposure40 | RNA3D38 | 38-target 95% CI |
|---|---|---|---|---|
| G0/Frozen | 0.2751 | 0.2861 | 0.2902 | [0.1973, 0.3860] |
| G0/LoRA | 0.3460 | 0.3714 | 0.3772 | [0.2682, 0.4856] |
| G0/Full | 0.2986 | 0.3104 | 0.3120 | [0.2135, 0.4074] |
| G1/Frozen | 0.2534 | 0.2665 | 0.2646 | [0.1557, 0.3721] |
| G1/LoRA | 0.3682 | 0.3844 | 0.3892 | [0.2885, 0.4868] |
| G1/Full | 0.2787 | 0.3019 | 0.2973 | [0.2034, 0.3893] |
| Scratch-N/Full | 0.1935 | 0.2098 | 0.2103 | [0.0621, 0.3557] |

**Supplementary Data D056.** All 21 validation-selected fits: seed-specific mean-target Spearman.

**Table S42.** All registered BRIDGE ranking contrasts. RNA3D targets and CASP clusters are resampled separately.

| Paired contrast | $\Delta\rho_{\mathbf{target}}$ | 95% CI |
|---|---|---|
| RNA3D38: G0/Full – Scratch-N/Full | 0.1018 | [-0.0473, 0.2598] |
| RNA3D38: G0/LoRA – G0/Frozen | 0.0871 | [-0.0111, 0.1812] |
| RNA3D38: G0/Full – G0/Frozen | 0.0219 | [-0.0838, 0.1243] |
| RNA3D38: G0/LoRA – G0/Full | 0.0652 | [0.0038, 0.1260] |
| RNA3D38: G1/Full – Scratch-N/Full | 0.0870 | [-0.0622, 0.2409] |
| RNA3D38: G1/LoRA – G1/Frozen | 0.1245 | [0.0344, 0.2148] |
| RNA3D38: G1/Full – G1/Frozen | 0.0326 | [-0.0716, 0.1359] |
| RNA3D38: G1/LoRA – G1/Full | 0.0919 | [0.0324, 0.1535] |
| RNA3D38: G1/Frozen – G0/Frozen | -0.0255 | [-0.0680, 0.0164] |
| RNA3D38: G1/LoRA – G0/LoRA | 0.0120 | [-0.0324, 0.0567] |
| RNA3D38: G1/Full – G0/Full | -0.0148 | [-0.0568, 0.0250] |
| CASP16: G0/Full – Scratch-N/Full | 0.2547 | [0.1644, 0.3714] |
| CASP16: G0/LoRA – G0/Frozen | 0.0861 | [-0.0056, 0.1638] |
| CASP16: G0/Full – G0/Frozen | 0.1673 | [0.0707, 0.2750] |
| CASP16: G0/LoRA – G0/Full | -0.0812 | [-0.2088, -0.0027] |
| CASP16: G1/Full – Scratch-N/Full | 0.2109 | [0.0722, 0.3497] |
| CASP16: G1/LoRA – G1/Frozen | 0.0578 | [-0.0618, 0.1353] |
| CASP16: G1/Full – G1/Frozen | 0.1212 | [0.0188, 0.2271] |
| CASP16: G1/LoRA – G1/Full | -0.0635 | [-0.2109, 0.0508] |
| CASP16: G1/Frozen – G0/Frozen | 0.0023 | [-0.0303, 0.0371] |
| CASP16: G1/LoRA – G0/LoRA | -0.0261 | [-0.0672, 0.0017] |
| CASP16: G1/Full – G0/Full | -0.0438 | [-0.1115, 0.0145] |

**Supplementary Data D058.** RNA3D38: per-target Spearman, BRIDGE averaged across three runs.

**Supplementary Data D059.** CASP16: per-target Spearman, BRIDGE averaged across three runs.

#### 7.3 Released inverse-folding encoders on the quality-prediction cohorts

Released gRNAde and NA-MPNN encoders use the same 20-epoch MLP fitting protocol, RNA3D training/validation sets and common-heavy RMSD labels as BRIDGE. Each validation-selected model is evaluated unchanged on RNA3D38 (5,814 decoys/38 targets) and CASP16 (3,789 submissions/20 targets). Three downstream seeds vary the readout, order and dropout while the released checkpoint remains fixed. Table S40 averages seed-specific target correlations, and Section 9 gives paired cross-model contrasts. These are RNA3D-adapted quality predictors, not zero-shot inverse-folding likelihoods or ATOM3D-trained models.

The released-model ranking depends on adaptation and cohort. RNA3D38 LoRA mean-target correlations are 0.3533 (gRNAde) and 0.4063 (NA-MPNN), compared with G0/G1 0.3772/0.3892. CASP16 frozen/LoRA/full values are 0.2806/0.1745/0.1604 for gRNAde and 0.2638/0.2897/0.2699 for NA-MPNN, versus G0 0.1185/0.2046/0.2857. Thus neither BRIDGE nor full adaptation is uniformly best. These are contextual deployed-model comparisons, not controlled architecture or pretraining-data ablations.

**Table S43. Released-encoder pretraining-inventory matches on RNA3D38 and CASP16.**

| Encoder | Cohort | Targets | PDB matches | Exact-sequence matches |
|---|---|---|---|---|
| gRNAde | RNA3D38 | 38 | 2 | 2 |
| gRNAde | CASP16 | 20 | 0 | 2 |
| NA-MPNN | RNA3D38 | 38 | 32 | – |
| NA-MPNN | CASP16 | 20 | 3 | – |

The exposure counts use the archived provenance inventories and the exact scored target lists. gRNAde has two RNA3D38 PDB/exact-sequence matches and two CASP16 exact-sequence matches (R1263/R1264), despite zero CASP16 PDB matches. NA-MPNN has 32/38 and 3/20 PDB-inventory matches. Complete source-sequence metadata is unavailable, so dashes denote unknown exact-sequence overlap rather than zero. gRNAde's inventory includes train/validation conformers and is not a checkpoint-specific training receipt. Neither inventory establishes homology/family independence. RNA3D38 screening was defined against BRIDGE and does not make it unexposed for every comparator. Inputs and native encoder dropout also remain unmatched as specified in Section 8.

**Supplementary Data D061.** Individual released-encoder quality-prediction seed metrics.

**Supplementary Data D062.** Per-target Spearman averaged over three seeds.

#### 7.4 Released ARES evaluation

ARES [9] was evaluated on RNA3D38 with its released checkpoint and native preprocessing, without RNA3D retraining. Consequently, this comparison measures zero-shot transfer of ARES against RNA3D-adapted predictors rather than matched-domain training.

CPU inference uses Python 3.8, PyTorch 1.5.0, ARES e3nn 0.1.0, Atom3D 0.2.6 and PyTorch Lightning 1.1.5. Gemmi converts mmCIF to PDB, and Reduce 4.16 (`-NOADJust -OH -Quiet`) adds hydrogen coordinates without changing heavy atoms. Conversion error is bounded by 0.000501 Å per coordinate component. Native hydrogen-inclusive normalization and admission are retained for all 5,814 inputs; selected/all-atom ratios are 0.62956–0.64112. Eight reference ATOM3D scores agree with published values within $1.38 \times 10^{-6}$. No PDB/canonical-sequence match to RNA3D38 was found in the known 14-training/4-validation ARES target inventory, although complete construct and homology exposure remain unknown.

**Table S44.** RNA3D38 released ARES comparison. Regret is lower-is-better. Inputs and training domains are not matched.

| Model/mode | Training | $\rho_{\text{target}}$ | Seed SD | $\rho_{\text{pooled}}$ | Regret (Å) |
|---|---|---|---|---|---|
| G0/Frozen | RNA3D fitted | 0.2902 | 0.0405 | 0.5665 | 1.5794 |
| G0/LoRA | RNA3D fitted | 0.3772 | 0.0297 | 0.6255 | 1.5911 |
| G0/Full | RNA3D fitted | 0.3120 | 0.0535 | 0.6148 | 1.8268 |
| G1/Frozen | RNA3D fitted | 0.2646 | 0.0294 | 0.5531 | 1.6226 |
| G1/LoRA | RNA3D fitted | 0.3892 | 0.0858 | 0.6366 | 1.5587 |
| G1/Full | RNA3D fitted | 0.2973 | 0.0167 | 0.5949 | 1.8093 |
| Scratch-N/Full | RNA3D fitted | 0.2103 | 0.0236 | 0.5839 | 2.2221 |
| ARES released | No RNA3D refit | -0.0186 | – | 0.0084 | 2.1659 |

ARES target-macro 95% CI: [-0.1387, 0.1065]. No target has undefined correlation.

**Table S45.** Conditional same-target system contrasts, not matched-training method superiority.

| BRIDGE minus released ARES | $\Delta\rho_{\text{target}}$ | 95% CI |
|---|---|---|
| G0/Frozen | 0.3088 | [0.1760, 0.4414] |
| G0/LoRA | 0.3958 | [0.2346, 0.5580] |
| G0/Full | 0.3306 | [0.1898, 0.4745] |
| G1/Frozen | 0.2832 | [0.1363, 0.4344] |
| G1/LoRA | 0.4078 | [0.2677, 0.5479] |
| G1/Full | 0.3159 | [0.1712, 0.4622] |
| Scratch-N/Full | 0.2289 | [0.0782, 0.3745] |

ARES uses one released model; BRIDGE values average three RNA3D-adapted fits. Paired differences compare these fitted systems rather than architectures under equal data or tuning. ARES was not evaluated on CASP16.

#### 7.5 RMSD distributions and decoy-domain diagnostic

**Table S46.** Measured RMSD distributions on the evaluated cohorts. Quantiles and fractions pool all models. The final column is the median of separate within-target Q75–Q25 ranges, not the pooled IQR. RNA3D38/CASP16 use common-heavy labels. ATOM3D retains its published RMSD convention.

| Evaluation cohort | Targets / models | Median [Q25,Q75], Å | P95, Å | $<$ 4Å (%) | Target IQR, Å |
|---|---|---|---|---|---|
| RNA3D38 | 38 / 5,814 | 2.98 [1.97, 4.32] | 18.98 | 71.52 | 0.97 |
| CASP16 | 20 / 3,789 | 11.76 [7.43, 20.42] | 38.08 | 9.40 | 7.68 |
| ATOM3D RSR | 4 / 4,000 | 8.91 [6.77, 13.36] | 16.73 | 1.23 | 3.15 |

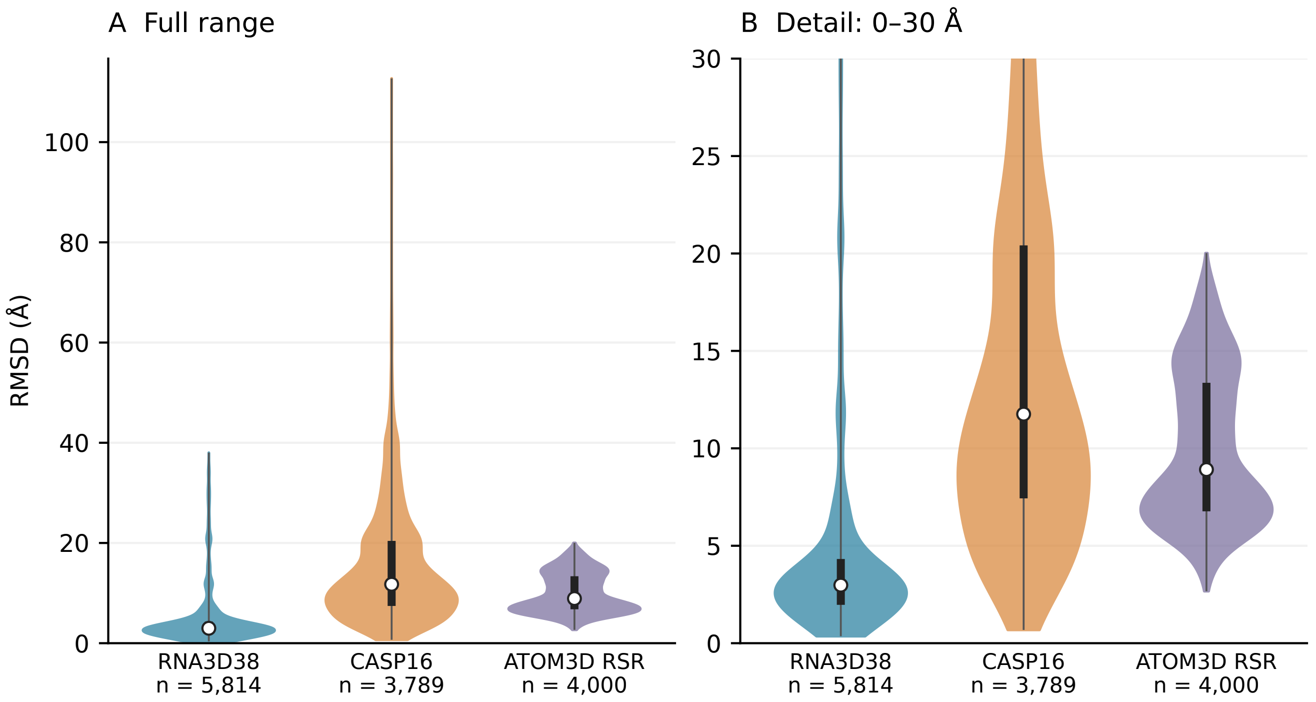


Figure S1: RMSD distributions of the evaluated RNA3D38, CASP16 and ATOM3D RSR models. (A) Full observed range. (B) A 0–30Å view of the same distributions. Both panels use every model, including the tails outside panel B. No decoy is excluded or winsorized. Violin widths show Gaussian kernel-density estimates with Scott bandwidths, normalized to equal maximum width within each cohort. Width does not encode cohort size. Black bars and white points denote pooled Q25–Q75 and medians. Thin lines span observed minima/maxima. These summaries weight decoys equally, not targets. RNA3D38 and CASP16 use the same common-heavy-atom alignment protocol. ATOM3D uses its published RMSD labels. Target composition, lengths and native support differ, so the plot does not isolate a generator effect.

CASP16 has a pooled median RMSD of 11.76 Å versus 2.98 Å for RNA3D38; median within-target IQRs are 7.68 and 0.97 Å, respectively. RNA3D38/CASP16 share the common-heavy-atom alignment convention, whereas ATOM3D retains published RMSD labels. RNA identities, lengths, reference masks and generation methods also differ, so the distributions do not isolate a generator effect or establish which cohort is easier to rank. These are descriptive distributions, not confidence intervals; a monotone score calibration cannot improve Spearman correlation.

**Supplementary Data D066.** Exploratory target-bootstrap diagnostics.

Generator-specific and alternative-label diagnostics show limited zero-shot ARES transfer to this neural-predictor bank. Separating RMSD-range restriction from generator-specific effects would require matched RNA targets and RMSD support across generators; that experiment and RNA3D-matched ARES retraining were not performed.

### 8 Released inverse-folding encoder comparisons

#### 8.1 Models, inputs and exposure

We use gRNAde v1 (official commit 2453b18778a1, released gRNAde drop3d@0.75 maxlen@500.h5) and NA-MPNN design (commit 9fabc2482092, s 19137.pt), not its specificity model. gRNAde supplies the scalar width-128 state after conformer pooling, before decoding. NA-MPNN uses its encoder output. Neither downstream adapter observes native nucleotide identities, a ligand, a protein partner or a quality label as an input feature. Structural information differs: gRNAde sees observed N1/N9 and base-derived secondary structure, whereas NA-MPNN uses twelve backbone atoms plus deterministic virtual N. BRIDGE uses P and five ribose atoms plus virtual N.

The gRNAde adapter respects chain and author-number segments for local/sequence features while retaining spatial and secondary-structure edges. Secondary structure uses the same observed base pairs as the official featurizer, without unnecessary dot-bracket coloring. NA-MPNN encodes residues with valid O4′/C1′/C2’ frames and scatters zero embeddings to frame-invalid original rows. Labels, rows, graph pooling denominators and readout edges are not dropped to improve compatibility. These are released weights plus benchmark-preserving adapters, not exact native-preprocessor reproductions. Native encoder dropout remains 0.5 for gRNAde and 0 for NA-MPNN. No coordinate augmentation is added.

**Table S47. Released-encoder pretraining-inventory matches across downstream benchmarks.**

| Cohort | gRNAde pool | NA-MPNN train list |
|---|---|---|
| Pretraining-test | 1/68 | 42/68 |
| TR60 test | 28/43 | 36/43 |
| RB9 | 1/9 | 7/9 |
| JL10 | 8/10 | 9/10 |
| TL12 | 9/12 | 8/12 |
| TE18 | 9/17 | 14/17 |
| RNAProt test | 160/191 | 115/191 |
| RSR test | 4/4 | 3/4 |

Counts flag provenance-PDB matches, not downstream-label leakage. gRNAde's inventory is the conservative official Das train-plus-validation pool and associated conformers, not checkpoint-specific training receipts. NA-MPNN's published design-training PDB list does not provide complete chain sequences here: exact-sequence and homology independence remain unknown. BRIDGE uses the documented test-excluded pretraining panel, although shared 3GS5 provenance remains in its pretraining-test diagnostic. Zero inventory matches cannot establish unseen-family transfer.

#### 8.2 Sequence recovery and decoding

All 68 pretraining-test records and 5568 residues are retained, with 11 dependence components. Unique experimental-mmCIF mapping recovers original row identities and atoms without replacing coordinates. Maximum discrepancy is below 0.000867Å. BRIDGE is evaluated using fixed epoch-100 archived predictions from three pretraining seeds. Public models each use one released checkpoint and three decoding seeds, one sequence draw per record per seed, temperature 0.1, and no native sequence context. gRNAde decodes sequentially. NA-MPNN uses a seeded random order and only A/G/C/U under the official shared-token convention (indices 21/23/22/24). These are not three independently pretrained public models.

BRIDGE's archived soft predicted-prefix rollout emits argmax sequences. Primary public values instead score categorical sampled sequences. As a secondary diagnostic, argmax conditional on generated hard prefixes is reported below. It is neither teacher-forced recovery nor the same decoder contract as BRIDGE. Public models have no comparable trained $\chi$ head, so no angular result is invented. The common 26-record/2148-residue subset excludes provenance PDB matches to either public inventory and was fixed before the new inference. It is a sensitivity, not a sequence/family-unseen certification. Recovery does not select downstream checkpoints or objectives.

**Table S48. Pretrained-model sequence recovery (%).**

| | All 68 records | | Common 26-record subset | |
|---|---|---|---|---|
| Model | Pooled (%) | Macro (%) | Pooled (%) | Macro (%) |
| Sequence-only | 60.88 | 58.61 | 58.30 | 56.88 |
| S0 | 62.19 | 59.83 | 59.75 | 58.75 |
| S0+PairGeom | 62.45 | 60.39 | 60.40 | 59.17 |
| S0+$\chi$ | <u>62.76</u> | <u>60.43</u> | 60.27 | 58.75 |
| G0 | 62.38 | 60.05 | <u>61.30</u> | <u>60.14</u> |
| G1 | 61.81 | 59.47 | 59.71 | 58.41 |
| gRNAde | 56.75 | 54.28 | 52.76 | 49.92 |
| NA-MPNN | **70.62** | **71.61** | **64.71** | **66.01** |

All 68 records contain 5,568 residues. The common 26-record subset contains 2,148. Macro values weight records, not PDBs, equally. All six BRIDGE variants, including sequence-only, average three pretraining runs. Public models average three decoding draws from one checkpoint. BRIDGE uses soft-prefix argmax decoding, whereas public entries are sequences sampled at temperature 0.1. Inputs and exposure differ. Bold and underline mark the best and second-best distinct values among all displayed models in each column. Emphasis is descriptive. In this table, released models are evaluated for sequence recovery only. Ties at the displayed precision share emphasis.

**Table S49. Generated-hard-prefix argmax sequence recovery.**

| Subset | Model | Pooled (%) | Record macro (%) |
|---|---|---|---|
| All 68 | gRNAde | 56.94 | 54.39 |
| Inventory 26 | gRNAde | 53.07 | 50.11 |
| All 68 | NA-MPNN | 70.79 | 71.82 |
| Inventory 26 | NA-MPNN | 64.91 | 66.36 |

The second table reports the alternative generated-hard-prefix argmax diagnostic.

**Supplementary Data D070.** Sequence recovery by seed.

#### 8.3 Downstream fitting and aggregation

Released-encoder classification retains the CoBRA and RNAProt memberships and fitting rules in Sections 1.9–1.10: 120/40 epochs, respectively, with an 82,689-parameter GNN, dropout 0.3, raw embeddings, C4′-kNN16 geometry and a 500-node sample cap. Only encoder-prefix parameters are adapted; sequence decoders are excluded. External CoBRA retains 48 set-qualified records from 44 PDBs, including duplicate chains and 8F0N, with the exclusion sensitivity reported separately.

RSR retains the 12,479/4,000/4,000 decoy split and published labels. Its 16,641-parameter mean-pooling MLP follows the 20-epoch recipe in Section 1.10, with gradient clipping at 1 and the earliest finite validation equal-target Spearman maximum selecting the checkpoint.

All released encoders use frozen, LoRA and full adaptation with three downstream seeds; the checkpoint is fixed. BRIDGE CoBRA full/LoRA pairs source and downstream seeds, whereas its RNAProt/RSR panels fix source seed 0. Classification scores mean probabilities; ranking averages seed-specific correlations. The 18 public CoBRA fits also supply apo/holo predictions without refitting. Section 9 provides the designated paired contrasts; other comparisons are descriptive. RNAProt full-unit macro AP remains undefined because 55 of 177 units lack positives.

**Supplementary Data D071.** CoBRA AP by encoder, adaptation mode and evaluation set.

Supplementary Data D071 gives pooled AP per set and external PDB-macro AP with a GNN readout. For BRIDGE full/LoRA, the paired pretraining/downstream seed convention differs from the fixed-encoder frozen panel. The following public-model metrics and seed results preserve those distinctions.

**Table S50. Released-encoder probability-ensemble metrics on CoBRA and RNAProt.**

| Task | Model | Mode | AP | Macro AP | AUROC | MCC .5 / val / oracle |
|---|---|---|---|---|---|---|
| CoBRA | gRNAde | frozen | 0.6729 | 0.7003 | 0.7897 | 0.4333 / 0.4406 / 0.4497 |
| CoBRA | gRNAde | LoRA | 0.6574 | 0.7151 | 0.7841 | 0.4336 / 0.4384 / 0.4592 |
| CoBRA | gRNAde | full | 0.6416 | 0.7051 | 0.7703 | 0.4290 / 0.4256 / 0.4400 |
| CoBRA | NA-MPNN | frozen | 0.6334 | 0.6987 | 0.7537 | 0.3960 / 0.4420 / 0.4525 |
| CoBRA | NA-MPNN | LoRA | 0.6796 | 0.7091 | 0.8068 | 0.4836 / 0.4852 / 0.4925 |
| CoBRA | NA-MPNN | full | 0.6829 | 0.7103 | 0.8102 | 0.4810 / 0.4759 / 0.4930 |
| RNAProt | gRNAde | frozen | 0.5992 | – | 0.7738 | 0.3473 / 0.3408 / 0.3567 |
| RNAProt | gRNAde | LoRA | 0.6220 | – | 0.7879 | 0.3504 / 0.3663 / 0.4052 |
| RNAProt | gRNAde | full | 0.6118 | – | 0.7803 | 0.3645 / 0.3700 / 0.3836 |
| RNAProt | NA-MPNN | frozen | 0.6335 | – | 0.7794 | 0.3753 / 0.3994 / 0.4238 |
| RNAProt | NA-MPNN | LoRA | 0.6383 | – | 0.7931 | 0.3895 / 0.3959 / 0.4090 |
| RNAProt | NA-MPNN | full | 0.6517 | – | 0.8028 | 0.3955 / 0.4045 / 0.4096 |

**Supplementary Data D073.** Individual downstream-seed results for released encoders.

The MLP and width32 GNN summaries are in Tables S32 and S33; Supplementary Data D074 provides the combined comparison.

**Supplementary Data D075.** Individual downstream-seed results for released-encoder RSR models, covering all six encoder/adaptation conditions.

**Supplementary Data D076.** Released-encoder per-puzzle ATOM3D correlations.

Per-puzzle correlations average three seed metrics. The independent test units remain the four puzzles.

#### 8.4 Released-encoder apo/holo inference without refitting

The 18 released-encoder CoBRA fits are applied unchanged to six apo/holo pairs, using their TR60 validation-selected checkpoints and thresholds. Seven graphs per state represent six targets, with chains of 6XJZ/6XJQ combined. The apo/holo supports are 605/607 labelled residues and 64 positives each. Apo labels are projected holo contacts.

Main Table 6 reports target-macro BA, validation-threshold MCC and AUROC under full adaptation. Table S19 retains full-adaptation AP and Table S51 gives frozen/LoRA results. Pooled metrics combine residues; macro AP weights six targets equally. The metrics use three-seed mean probabilities. This conformation comparison is descriptive and is not covered by the paired public-model contrasts in Section 9.

**Table S51. Public encoders on apo/holo: frozen and LoRA conditions.**

| Model | Mode | State | AP | Macro AP | AUROC | MCC .5 | MCC val | MCC oracle |
|---|---|---|---|---|---|---|---|---|
| gRNAde | frozen | apo | 0.1881 | 0.3632 | 0.6865 | 0.1055 | 0.0938 | 0.2062 |
| gRNAde | LoRA | apo | 0.2624 | 0.3844 | 0.7033 | 0.2051 | 0.2420 | 0.2896 |
| NA-MPNN | frozen | apo | **0.2858** | **0.4311** | **0.7531** | **0.3056** | **0.2878** | **0.3129** |
| NA-MPNN | LoRA | apo | **0.3140** | **0.4796** | **0.7886** | **0.2819** | **0.3000** | **0.3247** |
| gRNAde | frozen | holo | **0.5776** | 0.5975 | 0.8729 | 0.3886 | 0.4354 | **0.5951** |
| gRNAde | LoRA | holo | 0.5085 | 0.5628 | 0.8539 | 0.3418 | 0.3545 | 0.4759 |
| NA-MPNN | frozen | holo | 0.5403 | **0.6435** | **0.9107** | **0.4362** | **0.4949** | 0.5380 |
| NA-MPNN | LoRA | holo | **0.5484** | **0.6200** | **0.8927** | **0.4091** | **0.4537** | **0.5154** |

Full-adaptation conditions are in Table S19. Macro AP weights six targets equally; AUROC and MCC pool residues within state. Ranks compare encoders within the same state and adaptation mode. Inputs and pretraining exposure differ across encoders.

Four holo identities (3SUX, 2YIE, 4LVX, 2L8H) occur in TR60 training for both BRIDGE and the released-encoder adapters. The table below reports additional pretraining-inventory matches. gRNAde uses a conservative train-plus-validation/conformer inventory; NA-MPNN uses its published design-training PDB list. Inventory matches describe structural exposure, not necessarily downstream-label leakage. Inputs and pretraining histories differ, and family independence is unverified.

**Table S52. Released-encoder pretraining-inventory matches for apo/holo targets.**

| State | gRNAde provenance pool | NA-MPNN design train list |
|---|---|---|
| apo | 5/6 | 3/6 |
| holo | 5/6 | 3/6 |

##### 8.4.1 Individual downstream-seed results

Rows report individual downstream seeds; these do not represent independently pretrained released encoders.

**Supplementary Data D079.** Released-encoder individual-seed conformation AP.

##### 8.4.2 Per-target AP of the three-model probability ensembles

Column headings give apo/holo PDB pairs. The first table uses apo coordinates, the second holo coordinates.

The six-column mean gives target-macro AP. **Apo coordinates.**

**Supplementary Data D080.** Released-encoder per-target AP from apo coordinates.

**Holo coordinates.**

**Supplementary Data D081.** Released-encoder per-target AP from holo coordinates.

#### 8.5 Frozen binding-site readouts and controls

The frozen-head panel fixes source epoch 100 and encoder initialization seed 0 while varying three downstream head seeds. G0, G1 and Random-frozen use the same 82,689-parameter GNN, 331 TR60 training and 41 validation chains, 120 head epochs, Adam at $10^{-3}$, weight decay $10^{-4}$ and dropout 0.3. Earliest maximum validation AP every five epochs selects the checkpoint. Three probabilities are averaged before scoring. Random-frozen is a random encoder with a trained head, not an unfitted predictor. It is not Scratch-N/full. These seed and optimization contracts are not interchangeable with the source/downstream-paired full-adaptation panel.

The same frozen readouts are evaluated on 43 TR60 test chains, the external 48-record/44-PDB cohort, its 8F0N-excluded sensitivity, and the six apo/holo pairs.

**Table S53. Frozen-readout ligand-binding and conformation metrics.**

| Cohort | Frozen encoder | AP | Macro AP | AUROC | MCC .5 / val / oracle |
|---|---|---|---|---|---|
| TR60 test | G0 | 0.8031 | 0.8031 | 0.8696 | 0.5912 / 0.6277 / 0.6404 |
| TR60 test | G1 | 0.7812 | 0.7729 | 0.8564 | 0.5471 / 0.5688 / 0.5885 |
| TR60 test | Random-frozen | 0.6547 | 0.6990 | 0.7804 | 0.3818 / 0.3818 / 0.4546 |
| External 48 | G0 | 0.6274 | 0.6840 | 0.7551 | 0.3899 / 0.4227 / 0.4227 |
| External 48 | G1 | 0.6150 | 0.6744 | 0.7372 | 0.4030 / 0.3927 / 0.4136 |
| External 48 | Random-frozen | 0.5033 | 0.5918 | 0.6788 | 0.2600 / 0.2600 / 0.2855 |
| External without 8F0N | G0 | 0.6128 | 0.6768 | 0.7484 | 0.3793 / 0.4121 / 0.4121 |
| External without 8F0N | G1 | 0.5979 | 0.6671 | 0.7294 | 0.3927 / 0.3802 / 0.4045 |
| External without 8F0N | Random-frozen | 0.4876 | 0.5837 | 0.6687 | 0.2470 / 0.2470 / 0.2732 |
| Apo | G0 | 0.1844 | 0.2683 | 0.6711 | 0.1970 / 0.1084 / 0.2213 |
| Apo | G1 | 0.2639 | 0.3819 | 0.7021 | 0.1713 / 0.1753 / 0.2490 |
| Apo | Random-frozen | 0.1452 | 0.2128 | 0.5987 | 0.0655 / 0.0655 / 0.1347 |
| Holo | G0 | 0.5221 | 0.6617 | 0.8576 | 0.3996 / 0.4670 / 0.5260 |
| Holo | G1 | 0.5195 | 0.6710 | 0.8925 | 0.4248 / 0.4774 / 0.4946 |
| Holo | Random-frozen | 0.2891 | 0.4091 | 0.7232 | 0.1918 / 0.1918 / 0.2981 |

Paired 10,000-draw PDB-bootstrap intervals compare pretrained and random frozen encoders under the same readout protocol. They condition on the fitted models, are unadjusted and do not measure pretraining-seed variability.

**Table S54. Frozen pretrained-minus-random CoBRA AP contrasts.**

| Cohort | Contrast | Pooled Δ[95% CI] | Macro Δ[95% CI] |
|---|---|---|---|
| TR60 test | G0 minus Random-frozen | 0.1484 [0.0942, 0.1987] | 0.1041 [0.0502, 0.1554] |
| TR60 test | G1 minus Random-frozen | 0.1265 [0.0696, 0.1802] | 0.0738 [0.0214, 0.1262] |
| External 48 | G0 minus Random-frozen | 0.1241 [0.0644, 0.1818] | 0.0921 [0.0507, 0.1356] |
| External 48 | G1 minus Random-frozen | 0.1117 [0.0509, 0.1710] | 0.0826 [0.0428, 0.1253] |
| External without 8F0N | G0 minus Random-frozen | 0.1251 [0.0626, 0.1871] | 0.0931 [0.0518, 0.1387] |
| External without 8F0N | G1 minus Random-frozen | 0.1102 [0.0479, 0.1750] | 0.0834 [0.0436, 0.1272] |

### 9 Sensitivity analyses

These analyses were specified after initial test inspection and are exploratory. Paired percentile intervals use 10,000 shared draws (seed 20261016), condition on fitted models and are unadjusted for multiple comparisons and prior model search. Contrasts use full-precision values and do not alter checkpoint selection or the primary cohorts.

#### 9.1 Matched RNAProt learning-rate development

G0 and scratch were compared at three encoder learning rates with three downstream seeds per arm and a fixed head rate of $10^{-3}$. Both selected $10^{-5}$ by mean best validation AP. The subsequent held-out sensitivity gives G0/scratch AP 0.6478/0.6120 (Section 16.3), while the primary comparison retains $10^{-4}$. This limited development grid does not establish optimally tuned performance.

**Table S55. RNAProt validation AP by encoder learning rate.**

| Encoder LR | G0 | Scratch-N |
|---|---|---|
| $10^{-5}$ | 0.7625 | 0.7329 |
| $10^{-4}$ | 0.7547 | 0.6319 |
| $3 \times 10^{-4}$ | 0.7176 | 0.3730 |

#### 9.2 Exact-chain deduplication of external CoBRA

All three seed vectors and labels agree exactly between RB9/TE18 copies of 1NEM A, 1Q8N A and 2TOB A in every reported structural full-adaptation arm. Deduplication keeps the RB9 copy and removes 81 duplicate residues. Both distinct 8HB1 chains remain. The resulting cohorts are 45 records/44 PDBs/2288 residues and, after also excluding 8F0N, 44 records/43 PDBs/2239 residues. The original 48/47-record estimands remain primary and sensitivity references, respectively. Exclusions do not establish family independence. The archived Chi-off label in Tables S56 and S57 denotes S0+PairGeom.

**Table S56. External CoBRA exact-chain deduplication sensitivity.**

| Cohort | Model (full) | Records | PDBs | Pooled AP | Macro AP |
|---|---|---|---|---|---|
| Original 48 | S0 | 48 | 44 | 0.6883 | 0.7294 |
| Original 48 | Chi-off | 48 | 44 | 0.6602 | 0.7193 |
| Original 48 | S0+$\chi$ | 48 | 44 | 0.6803 | 0.7245 |
| Original 48 | G0 | 48 | 44 | 0.7028 | 0.7369 |
| Original 48 | G1 | 48 | 44 | 0.6934 | 0.7169 |
| Original 48 | Scratch-N | 48 | 44 | 0.6347 | 0.7003 |
| Exclude 8F0N | S0 | 47 | 43 | 0.6762 | 0.7238 |
| Exclude 8F0N | Chi-off | 47 | 43 | 0.6505 | 0.7147 |
| Exclude 8F0N | S0+$\chi$ | 47 | 43 | 0.6674 | 0.7183 |
| Exclude 8F0N | G0 | 47 | 43 | 0.6919 | 0.7313 |
| Exclude 8F0N | G1 | 47 | 43 | 0.6822 | 0.7116 |
| Exclude 8F0N | Scratch-N | 47 | 43 | 0.6219 | 0.6937 |
| Deduplicated | S0 | 45 | 44 | 0.6831 | 0.7294 |
| Deduplicated | Chi-off | 45 | 44 | 0.6509 | 0.7193 |
| Deduplicated | S0+$\chi$ | 45 | 44 | 0.6712 | 0.7245 |
| Deduplicated | G0 | 45 | 44 | 0.6945 | 0.7369 |
| Deduplicated | G1 | 45 | 44 | 0.6835 | 0.7169 |
| Deduplicated | Scratch-N | 45 | 44 | 0.6265 | 0.7003 |
| Dedup + exclude 8F0N | S0 | 44 | 43 | 0.6703 | 0.7238 |
| Dedup + exclude 8F0N | Chi-off | 44 | 43 | 0.6405 | 0.7147 |
| Dedup + exclude 8F0N | S0+$\chi$ | 44 | 43 | 0.6573 | 0.7183 |
| Dedup + exclude 8F0N | G0 | 44 | 43 | 0.6827 | 0.7313 |
| Dedup + exclude 8F0N | G1 | 44 | 43 | 0.6713 | 0.7116 |
| Dedup + exclude 8F0N | Scratch-N | 44 | 43 | 0.6128 | 0.6937 |

Pooled G0–Scratch AP differences are 0.0680 after deduplication and 0.0699 after both exclusions, versus 0.0681 in the original pool. Equal-PDB AP is unchanged by exact same-chain duplication within the affected single-chain PDBs. This is verified from scores, not assumed for arbitrary multichain duplicates. The paired effect intervals below resample complete PDBs, preserving the selected membership policy.

**Table S57. Paired CoBRA AP contrasts after deduplication.**

| Cohort | Contrast | Pooled Δ[95% CI] | Macro Δ[95% CI] |
|---|---|---|---|
| Original 48 | S0 minus G0 | -0.0146 [-0.0365, 0.0078] | -0.0075 [-0.0288, 0.0120] |
| Original 48 | S0 minus Scratch-N | 0.0535 [0.0205, 0.0826] | 0.0291 [0.0056, 0.0521] |
| Original 48 | G0 minus Chi-off | 0.0426 [0.0179, 0.0611] | 0.0176 [-0.0041, 0.0381] |
| Original 48 | G0 minus S0+$\chi$ | 0.0225 [0.0012, 0.0406] | 0.0124 [-0.0071, 0.0308] |
| Original 48 | G0 minus Scratch-N | 0.0681 [0.0320, 0.0985] | 0.0366 [0.0136, 0.0608] |
| Original 48 | G1 minus Scratch-N | 0.0587 [0.0226, 0.0904] | 0.0166 [-0.0084, 0.0424] |
| Exclude 8F0N | S0 minus G0 | -0.0156 [-0.0381, 0.0071] | -0.0075 [-0.0285, 0.0124] |
| Exclude 8F0N | S0 minus Scratch-N | 0.0544 [0.0210, 0.0839] | 0.0301 [0.0061, 0.0538] |
| Exclude 8F0N | G0 minus Chi-off | 0.0414 [0.0152, 0.0594] | 0.0166 [-0.0053, 0.0377] |
| Exclude 8F0N | G0 minus S0+$\chi$ | 0.0245 [0.0011, 0.0429] | 0.0130 [-0.0073, 0.0316] |
| Exclude 8F0N | G0 minus Scratch-N | 0.0700 [0.0331, 0.1020] | 0.0376 [0.0143, 0.0624] |
| Exclude 8F0N | G1 minus Scratch-N | 0.0603 [0.0236, 0.0939] | 0.0179 [-0.0082, 0.0437] |
| Deduplicated | S0 minus G0 | -0.0114 [-0.0327, 0.0098] | -0.0075 [-0.0288, 0.0120] |
| Deduplicated | S0 minus Scratch-N | 0.0566 [0.0233, 0.0853] | 0.0291 [0.0056, 0.0521] |
| Deduplicated | G0 minus Chi-off | 0.0436 [0.0172, 0.0620] | 0.0176 [-0.0041, 0.0381] |
| Deduplicated | G0 minus S0+$\chi$ | 0.0233 [0.0016, 0.0418] | 0.0124 [-0.0071, 0.0308] |
| Deduplicated | G0 minus Scratch-N | 0.0680 [0.0326, 0.0965] | 0.0366 [0.0136, 0.0608] |
| Deduplicated | G1 minus Scratch-N | 0.0570 [0.0234, 0.0848] | 0.0166 [-0.0084, 0.0424] |
| Dedup + exclude 8F0N | S0 minus G0 | -0.0124 [-0.0338, 0.0093] | -0.0075 [-0.0285, 0.0124] |
| Dedup + exclude 8F0N | S0 minus Scratch-N | 0.0575 [0.0242, 0.0864] | 0.0301 [0.0061, 0.0538] |
| Dedup + exclude 8F0N | G0 minus Chi-off | 0.0422 [0.0146, 0.0602] | 0.0166 [-0.0053, 0.0377] |
| Dedup + exclude 8F0N | G0 minus S0+$\chi$ | 0.0254 [0.0012, 0.0441] | 0.0130 [-0.0073, 0.0316] |
| Dedup + exclude 8F0N | G0 minus Scratch-N | 0.0699 [0.0338, 0.1001] | 0.0376 [0.0143, 0.0624] |
| Dedup + exclude 8F0N | G1 minus Scratch-N | 0.0585 [0.0242, 0.0881] | 0.0179 [-0.0082, 0.0437] |

#### 9.3 Symmetric seed influence and conditional protein-interface metrics

Leave-one-seed-out analyses recompute two-seed probability ensembles for all 16 conditions. These overlapping ensembles are influence diagnostics, not independent replicates. Full-GNN scratch AP is 0.6225 when seed 2 is omitted, versus the original 0.6225; omitting seed 0 or 1 gives 0.5852 or 0.6093. A weak individual seed need not proportionally reduce ensemble AP because AP depends on score ordering.

**Table S58. RNAProt leave-one-seed-out probability-ensemble AP.**

| Model/mode/head | All 3 | Omit 0 | Omit 1 | Omit 2 |
|---|---|---|---|---|
| G0/Frozen/GAT | 0.6384 | 0.6310 | 0.6334 | 0.6346 |
| G0/Frozen/GNN | 0.6471 | 0.6458 | 0.6548 | 0.6313 |
| G0/Full/GAT | 0.6397 | 0.6313 | 0.6430 | 0.6268 |
| G0/Full/GNN | 0.6415 | 0.6400 | 0.6287 | 0.6437 |
| G0/LoRA/GAT | 0.6574 | 0.6534 | 0.6496 | 0.6500 |
| G0/LoRA/GNN | 0.6550 | 0.6406 | 0.6572 | 0.6507 |
| G1/Frozen/GAT | 0.6523 | 0.6527 | 0.6429 | 0.6484 |
| G1/Frozen/GNN | 0.6328 | 0.6293 | 0.6322 | 0.6300 |
| G1/Full/GAT | 0.6412 | 0.6381 | 0.6380 | 0.6386 |
| G1/Full/GNN | 0.6506 | 0.6455 | 0.6522 | 0.6419 |
| G1/LoRA/GAT | 0.6477 | 0.6415 | 0.6442 | 0.6392 |
| G1/LoRA/GNN | 0.6402 | 0.6406 | 0.6346 | 0.6343 |
| Scratch-N/Full/GAT | 0.6153 | 0.6134 | 0.6140 | 0.6039 |
| Scratch-N/Full/GNN | 0.6225 | 0.5852 | 0.6093 | 0.6225 |
| Random-frozen/GAT | 0.5036 | 0.4982 | 0.5015 | 0.4975 |
| Random-frozen/GNN | 0.5160 | 0.5105 | 0.5192 | 0.5129 |

Strict macro AP over all 177 RNAProt PDB-connected units is undefined because 55 units lack positives. We separately report equal-unit AP on the 122 positive-support units and macro/pooled false-positive rates on the 55 all-negative units, using each ensemble's fixed validation threshold. This conditional AP is a different estimand from full-unit macro AP.

**Table S59. RNAProt positive-unit AP and negative-unit false-positive rates.**

| Model/mode/head | Positive macro AP | Negative macro FPR | Negative pooled FPR | Val. cut |
|---|---|---|---|---|
| G0/Frozen/GNN | 0.6197 | 0.0142 | 0.0140 | 0.64 |
| G0/LoRA/GNN | 0.6335 | 0.0069 | 0.0085 | 0.66 |
| G0/Full/GNN | 0.6214 | 0.0338 | 0.0339 | 0.58 |
| G0/Frozen/GAT | 0.6168 | 0.0217 | 0.0181 | 0.66 |
| G0/LoRA/GAT | 0.6194 | 0.0194 | 0.0205 | 0.52 |
| G0/Full/GAT | 0.6292 | 0.0230 | 0.0227 | 0.66 |
| G1/Frozen/GNN | 0.6198 | 0.0544 | 0.0435 | 0.66 |
| G1/LoRA/GNN | 0.6102 | 0.0117 | 0.0153 | 0.64 |
| G1/Full/GNN | 0.6208 | 0.0109 | 0.0098 | 0.54 |
| G1/Frozen/GAT | 0.6206 | 0.0353 | 0.0312 | 0.68 |
| G1/LoRA/GAT | 0.6172 | 0.0084 | 0.0104 | 0.60 |
| G1/Full/GAT | 0.6225 | 0.0113 | 0.0167 | 0.70 |
| Scratch-N/Full/GNN | 0.5896 | 0.0255 | 0.0208 | 0.66 |
| Scratch-N/Full/GAT | 0.6004 | 0.0384 | 0.0380 | 0.66 |
| Random-frozen/GNN | 0.5712 | 0.0342 | 0.0432 | 0.68 |
| Random-frozen/GAT | 0.5553 | 0.0461 | 0.0544 | 0.76 |

#### 9.4 Paired released-encoder contrasts and common-support inventories

Binary contrasts fix G0/full versus each released encoder/full with the GNN head. Ranking retains frozen, LoRA and full, all with MLP. No test winner is selected for these contrasts. Binary arrays are checked for identical record order, label arrays and PDB identities before pairing. RSR uses four puzzle units, RNA3D38 uses 38 targets and CASP16 uses the ten registered clusters. Ranking contrasts first average each target's correlation across the three seeds, then average targets. Cluster resampling retains whole clusters and recomputes equal-target weighting, not equal-cluster weighting.

**Full-cohort and inventory-unmatched binary results.** The common subset for a comparison excludes any PDB-connected unit with a known PDB or available exact-sequence match to that public encoder inventory. CoBRA additionally excludes downstream-training PDB 8F0N. Both models are evaluated on exactly that subset. Inventory matching is asymmetric in information: gRNAde has sequence metadata, whereas NA-MPNN's exact-sequence inventory remains unavailable. "Unmatched" therefore means no match under available audited keys, not family-unseen or absence of predictor/label exposure. Different pairwise subsets must not be combined into one cross-model ranking.

**Table S60. Inventory-unmatched support for binary prediction comparisons.**

| Task | Comparator | Records | Units | Residues | Positives |
|---|---|---|---|---|---|
| CoBRA | gRNAde | 21 | 18 | 846 | 308 |
| CoBRA | NA-MPNN | 10 | 9 | 394 | 151 |
| RNAProt | gRNAde | 30 | 27 | 1658 | 642 |
| RNAProt | NA-MPNN | 76 | 67 | 4746 | 1443 |

**Table S61. G0-versus-released-encoder binary prediction contrasts.**

| Task | Support | Contrast (full) | Pooled Δ[95% CI] | Macro Δ[95% CI] |
|---|---|---|---|---|
| CoBRA | Full cohort | G0 minus NA-MPNN | 0.0199 [-0.0229, 0.0607] | 0.0266 [-0.0061, 0.0596] |
| CoBRA | Full cohort | G0 minus gRNAde | 0.0612 [0.0240, 0.0987] | 0.0318 [0.0054, 0.0603] |
| CoBRA | Common unmatched | G0 minus NA-MPNN | -0.0030 [-0.0434, 0.0509] | 0.0028 [-0.0533, 0.0501] |
| CoBRA | Common unmatched | G0 minus gRNAde | 0.0500 [0.0035, 0.0948] | 0.0173 [-0.0156, 0.0518] |
| RNAProt | Full cohort | G0 minus NA-MPNN | -0.0102 [-0.0311, 0.0104] | – |
| RNAProt | Full cohort | G0 minus gRNAde | 0.0297 [-0.0003, 0.0652] | – |
| RNAProt | Common unmatched | G0 minus NA-MPNN | -0.0100 [-0.0558, 0.0254] | – |
| RNAProt | Common unmatched | G0 minus gRNAde | -0.0119 [-0.0607, 0.0261] | – |

The G0–NA-MPNN full-cohort difference is unresolved for both CoBRA and RNAProt. A point advantage on CoBRA is not a demonstrated general superiority. Even a resolved interval cannot equalize atom inputs, encoder dropout, pretraining exposure or training histories. The RNAProt all-unit macro estimand remains undefined in these contrasts. Empty cells are not zero effects.

**Ranking support and intervals.** Inventory exclusions are closed over the registered resampling clusters. Consequently CASP16 common supports can contain fewer targets than a simple count of unmatched target identities. RSR has no gRNAde-inventory-unmatched target, and only one NA-MPNN-unmatched puzzle: the latter is descriptive only, with no bootstrap interval. The six RNA3D38 targets outside the NA-MPNN PDB inventory are not a new adequately powered family-independent benchmark.

**Table S62. Inventory-unmatched target and cluster support for ranking comparisons.**

| Cohort | Comparator | Targets | Clusters |
|---|---|---|---|
| CASP16 | gRNAde | 16 | 9 |
| CASP16 | NA-MPNN | 15 | 8 |
| RNA3D38 | gRNAde | 36 | 36 |
| RNA3D38 | NA-MPNN | 6 | 6 |
| RSR | gRNAde | 0 | 0 |
| RSR | NA-MPNN | 1 | 1 |

**Table S63. G0-versus-released-encoder structure-ranking contrasts.**

| Cohort | Mode | Support | Contrast | Mean-target $\rho$ difference [95% CI] |
|---|---|---|---|---|
| CASP16 | Frozen | Full | G0 minus NA-MPNN | -0.1453 [-0.3155, 0.0191] |
| CASP16 | Frozen | Full | G0 minus gRNAde | -0.1622 [-0.2954, -0.0346] |
| CASP16 | Frozen | Unmatched | G0 minus NA-MPNN | -0.0496 [-0.1943, 0.0590] |
| CASP16 | Frozen | Unmatched | G0 minus gRNAde | -0.1462 [-0.3196, -0.0024] |
| CASP16 | LoRA | Full | G0 minus NA-MPNN | -0.0851 [-0.1976, 0.0072] |
| CASP16 | LoRA | Full | G0 minus gRNAde | 0.0301 [-0.1041, 0.1889] |
| CASP16 | LoRA | Unmatched | G0 minus NA-MPNN | -0.0683 [-0.2244, 0.0277] |
| CASP16 | LoRA | Unmatched | G0 minus gRNAde | 0.0643 [-0.0984, 0.2290] |
| CASP16 | Full | Full | G0 minus NA-MPNN | 0.0159 [-0.0187, 0.0560] |
| CASP16 | Full | Full | G0 minus gRNAde | 0.1254 [-0.0899, 0.3652] |
| CASP16 | Full | Unmatched | G0 minus NA-MPNN | 0.0392 [0.0185, 0.0701] |
| CASP16 | Full | Unmatched | G0 minus gRNAde | 0.2408 [0.0707, 0.4112] |
| RNA3D38 | Frozen | Full | G0 minus NA-MPNN | -0.0479 [-0.1489, 0.0570] |
| RNA3D38 | Frozen | Full | G0 minus gRNAde | 0.0240 [-0.1080, 0.1581] |
| RNA3D38 | Frozen | Unmatched | G0 minus NA-MPNN | -0.1224 [-0.2675, 0.0330] |
| RNA3D38 | Frozen | Unmatched | G0 minus gRNAde | 0.0340 [-0.1065, 0.1738] |
| RNA3D38 | LoRA | Full | G0 minus NA-MPNN | -0.0291 [-0.1080, 0.0511] |
| RNA3D38 | LoRA | Full | G0 minus gRNAde | 0.0240 [-0.0743, 0.1301] |
| RNA3D38 | LoRA | Unmatched | G0 minus NA-MPNN | 0.0199 [-0.1463, 0.1796] |
| RNA3D38 | LoRA | Unmatched | G0 minus gRNAde | 0.0065 [-0.0869, 0.1083] |
| RNA3D38 | Full | Full | G0 minus NA-MPNN | -0.0650 [-0.1317, 0.0022] |
| RNA3D38 | Full | Full | G0 minus gRNAde | 0.0012 [-0.0890, 0.0935] |
| RNA3D38 | Full | Unmatched | G0 minus NA-MPNN | 0.0133 [-0.1586, 0.2092] |
| RNA3D38 | Full | Unmatched | G0 minus gRNAde | -0.0147 [-0.0992, 0.0706] |
| RSR | Frozen | Full | G0 minus NA-MPNN | 0.0461 [-0.1330, 0.2086] |
| RSR | Frozen | Full | G0 minus gRNAde | 0.1154 [-0.0625, 0.2770] |
| RSR | Frozen | Unmatched | G0 minus NA-MPNN | -0.2435 (descriptive) |
| RSR | LoRA | Full | G0 minus NA-MPNN | 0.1906 [0.1164, 0.2649] |
| RSR | LoRA | Full | G0 minus gRNAde | 0.2737 [0.1790, 0.4063] |
| RSR | LoRA | Unmatched | G0 minus NA-MPNN | 0.2424 (descriptive) |
| RSR | Full | Full | G0 minus NA-MPNN | 0.1100 [-0.0172, 0.2371] |
| RSR | Full | Full | G0 minus gRNAde | 0.1857 [0.0364, 0.3317] |
| RSR | Full | Unmatched | G0 minus NA-MPNN | -0.0096 (descriptive) |

#### 9.5 Conformation exposure tiers

The six-target primary descriptive panel is retained. The four holo-seen pairs are 1ANR/2L8H, 3SUY/3SUX, 6WJR/2YIE and 7KD1/4LVX. The two with no direct audited TR60 match are 3D0X/3D0U and 6XJZ/6XJQ. The latter tier has not been cleared for family homology. Each row averages target-level metrics, calculated after averaging seed probabilities, rather than pooling different-length RNAs. Oracle MCC is optimized separately within each target and is nondeployable. Four-target support comprises 328 apo/330 holo labelled residues and 42 positives per state. Two-target support comprises 277 labelled residues and 22 positives per state. No model or threshold was reselected.

**Table S64. Conformation metrics by downstream-training exposure tier.**

| State | Tier | Model | Macro AP | Macro AUROC | MCC .5/val/oracle |
|---|---|---|---|---|---|
| apo | Seen (4) | S0/Full | 0.6424 | 0.8305 | 0.4127/0.3651/0.6680 |
| apo | Seen (4) | G0/Full | 0.6781 | 0.8565 | 0.4077/0.4460/0.6188 |
| apo | Seen (4) | G1/Full | 0.6564 | 0.8250 | 0.4712/0.4963/0.6084 |
| apo | Seen (4) | Scratch-N | 0.4136 | 0.7624 | 0.2724/0.2885/0.4840 |
| apo | No direct (2) | S0/Full | 0.2902 | 0.8498 | 0.3715/0.3749/0.4380 |
| apo | No direct (2) | G0/Full | 0.3201 | 0.8546 | 0.2425/0.2286/0.4206 |
| apo | No direct (2) | G1/Full | 0.3922 | 0.8430 | 0.3327/0.3300/0.4507 |
| apo | No direct (2) | Scratch-N | 0.2339 | 0.7687 | 0.1238/0.1770/0.3597 |
| holo | Seen (4) | S0/Full | 0.9235 | 0.9740 | 0.7417/0.7921/0.8667 |
| holo | Seen (4) | G0/Full | 0.9046 | 0.9673 | 0.7473/0.7789/0.8950 |
| holo | Seen (4) | G1/Full | 0.8862 | 0.9695 | 0.7580/0.7580/0.8551 |
| holo | Seen (4) | Scratch-N | 0.6495 | 0.9185 | 0.6237/0.5868/0.7057 |
| holo | No direct (2) | S0/Full | 0.3747 | 0.8714 | 0.2978/0.4039/0.4236 |
| holo | No direct (2) | G0/Full | 0.3970 | 0.8943 | 0.3619/0.3184/0.4978 |
| holo | No direct (2) | G1/Full | 0.5086 | 0.8746 | 0.3352/0.3018/0.5657 |
| holo | No direct (2) | Scratch-N | 0.2848 | 0.8203 | 0.2330/0.1854/0.4190 |

These small exposure tiers cannot identify causal effects of conformation or training exposure. Apo labels are projected holo annotations, and complete homology and historical checkpoint-exposure clearance remain unavailable.

### 10 RNA3D frozen controls and larger readouts

#### 10.1 Readout designs and fitting protocol

The MLP random-frozen control uses encoder initialization seed 0 and the same 20-epoch fitting and validation-selection protocol as frozen G0/G1. Each encoder has its own fixed training-derived normalization. The exploratory larger-head comparison includes MLP83k and ResidualGNN83k under frozen, LoRA and full adaptation for S0/G0/G1, together with random-frozen and scratch-full controls. Three downstream seeds vary initialization, ordering and dropout while the source/random encoder seed is fixed. Head comparisons were introduced after earlier test inspection.

MLP83k has 83,031 parameters: normalized 128-dimensional graph-mean embeddings enter widths 230–230–1 with ReLU and dropout 0.1. ResidualGNN83k has 83,201 parameters: normalized residue embeddings enter a width-128 projection and two residual self/mean-neighbour layers with LayerNorm, ReLU and dropout 0.1, followed by a scalar node readout and graph mean. Its fixed C4′ nearest-neighbour graph has up to 16 neighbours per residue, with padding masked on short chains. The encoder graph is unchanged. Parameter counts are approximately matched, but this comparison does not separately isolate topology, pooling, normalization and message passing.

Fitting follows Section 7.1, with the earliest finite validation target-macro Spearman maximum selecting the checkpoint; undefined correlations are ineligible. Scratch MLP83k selects epochs 1/20/1 and ResidualGNN83k selects 19/16/10. Aggregate results include all 22 conditions. The 48-fit seed and paired-contrast panels cover G0/G1 and controls; S0 seeds are reported separately without paired intervals.

#### 10.2 All-condition and selection-set results

Tables report means of seed-specific metrics and sample SD. Correlation and regret rankings compare all displayed conditions within each cohort; validation, individual seeds and contrasts are unranked.

**Table S65.** MLP frozen-control comparison. Values are means and sample SDs across three head seeds; ranks compare the three encoders within each cohort.

| Cohort | Frozen encoder | Target $\rho$ | Pooled $\rho$ | Regret (Å) |
|---|---|---|---|---|
| RNA3D38 | G0 | 0.2902 ± 0.0405 | **0.5665** ± 0.0310 | **1.5794** ± 0.0953 |
| RNA3D38 | G1 | 0.2646 ± 0.0294 | 0.5531 ± 0.0247 | 1.6226 ± 0.1395 |
| RNA3D38 | Random | **0.3097** ± 0.0875 | 0.5423 ± 0.0235 | 1.6604 ± 0.1214 |
| CASP16 | G0 | 0.1185 ± 0.0148 | **0.4017** ± 0.0167 | 15.0761 ± 1.9649 |
| CASP16 | G1 | **0.1207** ± 0.0088 | 0.3927 ± 0.0118 | 12.8689 ± 1.4896 |
| CASP16 | Random | 0.1087 ± 0.0413 | 0.3169 ± 0.0388 | **9.9912** ± 2.6701 |

**Table S66.** RNA3D38: all 22 readout/adaptation conditions. Mean and sample SD across three downstream seeds. Correlation is higher-is-better. Regret is lower-is-better. Ranks compare all 22 displayed conditions per metric. SDs are unranked.

| Head | Encoder | Mode | Target $\rho$ | Pooled $\rho$ | Regret (Å) |
|---|---|---|---|---|---|
| MLP83k | S0 | frozen | 0.2884 ± 0.0090 | 0.6181 ± 0.0366 | 1.5763 ± 0.1394 |
| MLP83k | S0 | LoRA | **0.3903** ± 0.0308 | **0.6608** ± 0.0267 | 1.6573 ± 0.2643 |
| MLP83k | S0 | full | 0.2987 ± 0.0083 | 0.6160 ± 0.0227 | 1.4981 ± 0.0307 |
| MLP83k | G0 | Frozen | 0.2639 ± 0.0563 | 0.6202 ± 0.0222 | 1.7081 ± 0.0769 |
| MLP83k | G0 | LoRA | 0.3413 ± 0.0365 | 0.6315 ± 0.0379 | 1.6026 ± 0.0821 |
| MLP83k | G0 | Full | 0.3428 ± 0.0151 | 0.6207 ± 0.0171 | 1.6890 ± 0.1672 |
| MLP83k | G1 | Frozen | 0.2576 ± 0.0040 | 0.6285 ± 0.0118 | 1.7247 ± 0.0431 |
| MLP83k | G1 | LoRA | 0.3493 ± 0.0356 | 0.6319 ± 0.0120 | 1.4775 ± 0.0398 |
| MLP83k | G1 | Full | 0.3122 ± 0.0169 | 0.6109 ± 0.0073 | 1.6409 ± 0.2619 |
| MLP83k | Random | Frozen | 0.3315 ± 0.0294 | 0.5918 ± 0.0163 | 1.4614 ± 0.1756 |
| MLP83k | Scratch | Full | 0.2372 ± 0.0433 | 0.5990 ± 0.0112 | 1.9142 ± 0.2926 |
| ResidualGNN83k | S0 | frozen | 0.2285 ± 0.0499 | 0.4948 ± 0.0787 | 1.7836 ± 0.1503 |
| ResidualGNN83k | S0 | LoRA | 0.2774 ± 0.0166 | 0.6375 ± 0.0036 | 1.5114 ± 0.0932 |
| ResidualGNN83k | S0 | full | 0.3223 ± 0.0187 | 0.6523 ± 0.0119 | 1.4583 ± 0.2050 |
| ResidualGNN83k | G0 | Frozen | 0.3444 ± 0.0247 | 0.5472 ± 0.0932 | **1.3866** ± 0.1161 |
| ResidualGNN83k | G0 | LoRA | 0.2905 ± 0.0101 | 0.6086 ± 0.0157 | 1.5699 ± 0.0674 |
| ResidualGNN83k | G0 | Full | 0.2776 ± 0.0356 | 0.5957 ± 0.0073 | 1.5950 ± 0.1921 |
| ResidualGNN83k | G1 | Frozen | 0.3183 ± 0.0294 | 0.5170 ± 0.0401 | 1.5871 ± 0.1031 |
| ResidualGNN83k | G1 | LoRA | 0.3601 ± 0.0452 | 0.6425 ± 0.0267 | 1.6409 ± 0.1436 |
| ResidualGNN83k | G1 | Full | 0.2959 ± 0.0700 | 0.5988 ± 0.0351 | 1.6589 ± 0.1188 |
| ResidualGNN83k | Random | Frozen | 0.2483 ± 0.0354 | 0.5269 ± 0.0558 | 1.5783 ± 0.0480 |
| ResidualGNN83k | Scratch | Full | 0.2723 ± 0.0321 | 0.5221 ± 0.0081 | 2.1495 ± 0.4064 |

**Table S67.** CASP16: all 22 readout/adaptation conditions. Mean and sample SD across three downstream seeds. Correlation is higher-is-better. Regret is lower-is-better. Ranks compare all 22 displayed conditions per metric. SDs are unranked.

| Head | Encoder | Mode | Target $\rho$ | Pooled $\rho$ | Regret (Å) |
|---|---|---|---|---|---|
| MLP83k | S0 | frozen | 0.1284 ± 0.0216 | 0.3963 ± 0.0283 | **7.9256** ± 0.9118 |
| MLP83k | S0 | LoRA | 0.0772 ± 0.0129 | 0.4707 ± 0.0050 | 16.0452 ± 1.1857 |
| MLP83k | S0 | full | 0.2376 ± 0.0517 | 0.5307 ± 0.0475 | 10.0940 ± 3.4209 |
| MLP83k | G0 | Frozen | 0.0580 ± 0.0342 | 0.2466 ± 0.0240 | 11.4201 ± 0.6863 |
| MLP83k | G0 | LoRA | 0.1162 ± 0.0416 | 0.4713 ± 0.0658 | 10.5309 ± 1.7129 |
| MLP83k | G0 | Full | 0.1928 ± 0.0329 | 0.4895 ± 0.0213 | 11.8862 ± 2.6961 |
| MLP83k | G1 | Frozen | 0.0029 ± 0.0798 | 0.2022 ± 0.0437 | 9.8614 ± 0.1512 |
| MLP83k | G1 | LoRA | 0.1352 ± 0.0328 | 0.4483 ± 0.0097 | 9.3015 ± 1.2746 |
| MLP83k | G1 | Full | 0.1758 ± 0.0672 | 0.5178 ± 0.0670 | 12.2386 ± 2.6919 |
| MLP83k | Random | Frozen | 0.0356 ± 0.0359 | 0.2618 ± 0.0243 | 15.8634 ± 3.9240 |
| MLP83k | Scratch | Full | 0.1276 ± 0.1183 | 0.3903 ± 0.0707 | 21.6746 ± 4.8199 |
| ResidualGNN83k | S0 | frozen | 0.2027 ± 0.0257 | 0.5233 ± 0.0164 | 10.1245 ± 0.7205 |
| ResidualGNN83k | S0 | LoRA | 0.1641 ± 0.0870 | 0.4780 ± 0.0356 | 13.6411 ± 2.7024 |
| ResidualGNN83k | S0 | full | 0.2785 ± 0.0535 | 0.5307 ± 0.0120 | 11.3265 ± 2.4523 |
| ResidualGNN83k | G0 | Frozen | 0.1851 ± 0.0235 | 0.5057 ± 0.0334 | 9.2920 ± 1.9842 |
| ResidualGNN83k | G0 | LoRA | 0.1772 ± 0.0529 | 0.5192 ± 0.0667 | 11.3653 ± 2.6068 |
| ResidualGNN83k | G0 | Full | **0.2901** ± 0.0249 | 0.5258 ± 0.0158 | 10.4840 ± 2.8104 |
| ResidualGNN83k | G1 | Frozen | 0.1417 ± 0.0153 | 0.4919 ± 0.0202 | 10.6234 ± 1.1984 |
| ResidualGNN83k | G1 | LoRA | 0.2094 ± 0.0107 | 0.5064 ± 0.0531 | 9.7173 ± 2.6848 |
| ResidualGNN83k | G1 | Full | 0.2695 ± 0.0595 | **0.5535** ± 0.0286 | 11.7001 ± 1.3571 |
| ResidualGNN83k | Random | Frozen | 0.0654 ± 0.0746 | 0.3340 ± 0.0811 | 9.4852 ± 3.3619 |
| ResidualGNN83k | Scratch | Full | 0.1330 ± 0.0592 | 0.4517 ± 0.0610 | 11.8629 ± 2.1854 |

**Supplementary Data D097.** Validation results and selected epochs for all 66 larger-head fits.

The original MLP G0/G1 frozen contrasts against random remain unresolved on both cohorts. The larger MLP83k does not resolve these frozen benefits either. With ResidualGNN83k, frozen G0–random effects are 0.0961 on RNA3D38 and 0.1198 on CASP16, with conditional intervals above zero. Corresponding G1 effects remain unresolved. Full G0–scratch is resolved on CASP16 for ResidualGNN83k (0.1571), but not MLP83k (0.0652). On RNA3D38 neither new-head full G0–scratch contrast is resolved. These conditional findings support readout-dependent transfer, not a uniformly superior encoder, head or adaptation mode. An unresolved contrast is not evidence of equivalence.

#### 10.3 Paired inference and complete seed reporting

Paired intervals resample 38 RNA3D targets or 10 CASP sequence–structure clusters, retaining equal-target point weighting and averaging seed metrics within each draw. They condition on fitted seeds, are unadjusted and exclude pretraining and retrospective-selection uncertainty. All four MLP-control and 32 larger-head Spearman contrasts are reported, including unresolved and negative estimates.

**Table S68.** All four MLP control contrasts and all 32 larger-readout contrasts. Intervals resample RNA3D targets or registered CASP clusters, conditional on the fitted seeds. These contrasts are unadjusted and exploratory, not confirmatory.

| Cohort | Candidate minus reference | $\Delta\,\rho_{\text{target}}$ [95% CI] |
|---|---|---|
| RNA3D38 | MLP: G0–Random (Frozen) | -0.0195 [-0.1344, 0.0940] |
| RNA3D38 | MLP: G1–Random (Frozen) | -0.0451 [-0.1698, 0.0755] |
| RNA3D38 | MLP83k: G0–Random (Frozen) | -0.0676 [-0.1897, 0.0449] |
| RNA3D38 | MLP83k: G0–Scratch (Full) | 0.1056 [-0.0366, 0.2425] |
| RNA3D38 | MLP83k: G1–Random (Frozen) | -0.0739 [-0.2023, 0.0461] |
| RNA3D38 | MLP83k: G1–Scratch (Full) | 0.0751 [-0.0679, 0.2151] |
| RNA3D38 | ResidualGNN83k: G0–Random (Frozen) | 0.0961 [0.0120, 0.1731] |
| RNA3D38 | ResidualGNN83k: G0–Scratch (Full) | 0.0053 [-0.1014, 0.1102] |
| RNA3D38 | ResidualGNN83k: G1–Random (Frozen) | 0.0700 [-0.0246, 0.1602] |
| RNA3D38 | ResidualGNN83k: G1–Scratch (Full) | 0.0236 [-0.0696, 0.1142] |

**Table S68. (continued)**

| Cohort | Candidate minus reference | $\Delta\,\rho_{\mathbf{target}}$ [95% CI] |
|---|---|---|
| RNA3D38 | ResGNN–MLP: G0/Frozen | 0.0805 [-0.0085, 0.1777] |
| RNA3D38 | ResGNN–MLP: G0/LoRA | -0.0508 [-0.1405, 0.0282] |
| RNA3D38 | ResGNN–MLP: G0/Full | -0.0652 [-0.1389, 0.0050] |
| RNA3D38 | ResGNN–MLP: G1/Frozen | 0.0607 [-0.0619, 0.1865] |
| RNA3D38 | ResGNN–MLP: G1/LoRA | 0.0108 [-0.0550, 0.0745] |
| RNA3D38 | ResGNN–MLP: G1/Full | -0.0163 [-0.0962, 0.0647] |
| RNA3D38 | ResGNN–MLP: Random/Frozen | -0.0832 [-0.1889, 0.0195] |
| RNA3D38 | ResGNN–MLP: Scratch/Full | 0.0352 [-0.0819, 0.1519] |
| CASP16 | MLP: G0–Random (Frozen) | 0.0097 [-0.1375, 0.1682] |
| CASP16 | MLP: G1–Random (Frozen) | 0.0120 [-0.1206, 0.1579] |
| CASP16 | MLP83k: G0–Random (Frozen) | 0.0224 [-0.1080, 0.1727] |
| CASP16 | MLP83k: G0–Scratch (Full) | 0.0652 [-0.0577, 0.1927] |
| CASP16 | MLP83k: G1–Random (Frozen) | -0.0327 [-0.1643, 0.1143] |
| CASP16 | MLP83k: G1–Scratch (Full) | 0.0482 [-0.0696, 0.1742] |
| CASP16 | ResidualGNN83k: G0–Random (Frozen) | 0.1198 [0.0065, 0.2988] |
| CASP16 | ResidualGNN83k: G0–Scratch (Full) | 0.1571 [0.0023, 0.3323] |
| CASP16 | ResidualGNN83k: G1–Random (Frozen) | 0.0763 [-0.0401, 0.2512] |
| CASP16 | ResidualGNN83k: G1–Scratch (Full) | 0.1366 [-0.0060, 0.2981] |
| CASP16 | ResGNN–MLP: G0/Frozen | 0.1272 [0.0604, 0.2158] |
| CASP16 | ResGNN–MLP: G0/LoRA | 0.0610 [-0.0058, 0.1378] |
| CASP16 | ResGNN–MLP: G0/Full | 0.0973 [0.0483, 0.1579] |
| CASP16 | ResGNN–MLP: G1/Frozen | 0.1388 [0.0745, 0.2140] |
| CASP16 | ResGNN–MLP: G1/LoRA | 0.0742 [0.0069, 0.1281] |
| CASP16 | ResGNN–MLP: G1/Full | 0.0937 [0.0499, 0.1428] |
| CASP16 | ResGNN–MLP: Random/Frozen | 0.0298 [-0.0517, 0.1028] |
| CASP16 | ResGNN–MLP: Scratch/Full | 0.0053 [-0.1350, 0.1231] |

**Supplementary Data D099.** All 48 selected-fit target-macro correlations.

### 11 Auxiliary-objective and small-readout comparisons

#### 11.1 Scope, aggregation and selection

The exploratory panels evaluate small CoBRA/RNAProt GNNs, released-encoder RNAProt GAT readouts, S0, and residual RSR GNN widths 32/64. Native epoch-100 pretraining seed 0 is fixed and three downstream seeds are fitted per condition. Classification reports metrics of mean probabilities; ranking reports means of seed-specific metrics. Training lasts 120 epochs for CoBRA, 40 for RNAProt and 20 for ranking. Original-width frozen S0 CoBRA also uses the matched 120-epoch protocol described in Section 14. All checkpoints are validation-selected; the panels were introduced after initial benchmark inspection.

Small residue-level GNNs have width32/64 and 8,385/24,961 parameters (two non-residual message-passing layers, dropout 0.3, C4′-kNN16). Residual RSR heads have 8,513/25,217 parameters. These are not exact MLP capacity matches. Random-N denotes a randomly initialized encoder kept frozen, whereas Scratch-N full trains the same architecture from initialization. Public gRNAde/NA-MPNN inputs and pretraining exposure remain unmatched.

Small-head rankings are within head/width and cohort, with displayed ties sharing emphasis. SDs, seeds, validation results and contrasts are unranked. Oracle MCC uses test labels. RNAProt strict macro AP remains undefined because 55 of 177 units have no positives.

**Table S69. Small residue-level GNN readouts across tasks.** All 34 encoder/mode/width conditions are retained. Native pretraining seed 0 is fixed across three downstream fits per condition. Values are pooled AP of three-fit probability ensembles: external CoBRA has 48 records/44 PDBs, RNAProt has 191 graphs/177 PDB units, and apo/holo reuse the six-pair inference protocol. These columns are distinct tasks, not interchangeable estimates. Widths 32/64 use 8,385/24,961 parameters. Within each width and task, bold and underline indicate the best and second-best distinct estimates descriptively. Complete metrics and external-set results are in Supplementary Data D100–D104. Original-width released-encoder GAT-minus-GNN contrasts are reported separately in D108.

| Width | Encoder | Mode | External AP | RNAProt AP | Apo AP | Holo AP |
|---|---|---|---|---|---|---|
| 32 | S0 | frozen | 0.6102 | 0.6506 | 0.2122 | 0.4217 |
| 32 | S0 | LoRA | 0.6377 | 0.6408 | 0.3484 | 0.6880 |
| 32 | S0 | full | 0.6436 | 0.6323 | 0.3952 | 0.6862 |
| 32 | G0 | frozen | 0.6150 | 0.6412 | 0.2347 | 0.4062 |
| 32 | G0 | LoRA | 0.6589 | 0.6333 | 0.3434 | 0.6542 |
| 32 | G0 | full | 0.6633 | 0.6365 | 0.3711 | **0.7352** |
| 32 | G1 | frozen | 0.5993 | 0.6492 | 0.2497 | 0.3689 |
| 32 | G1 | LoRA | 0.6548 | 0.6471 | 0.3425 | 0.6853 |
| 32 | G1 | full | **0.6729** | 0.6496 | <u>0.4130</u> | <u>0.7147</u> |
| 32 | Scratch-N | full | 0.5906 | 0.5849 | 0.2239 | 0.4547 |
| 32 | Random-N | frozen | 0.4727 | 0.4900 | 0.1706 | 0.3019 |
| 32 | gRNAde | frozen | 0.6572 | 0.6042 | 0.1768 | 0.4685 |
| 32 | gRNAde | LoRA | 0.6420 | 0.6344 | 0.3170 | 0.5198 |
| 32 | gRNAde | full | 0.6420 | 0.6250 | 0.2893 | 0.5616 |
| 32 | NA-MPNN | frozen | 0.6349 | 0.6425 | 0.3365 | 0.4902 |
| 32 | NA-MPNN | LoRA | <u>0.6684</u> | **0.6591** | 0.2855 | 0.5738 |
| 32 | NA-MPNN | full | <u>0.6684</u> | <u>0.6520</u> | **0.4248** | 0.6487 |
| 64 | S0 | frozen | 0.6232 | 0.6397 | 0.2587 | 0.4869 |
| 64 | S0 | LoRA | 0.6434 | 0.6458 | 0.3009 | 0.5897 |
| 64 | S0 | full | 0.6675 | 0.6402 | <u>0.3748</u> | <u>0.7249</u> |
| 64 | G0 | frozen | 0.6347 | 0.6423 | 0.1989 | 0.4653 |
| 64 | G0 | LoRA | 0.6536 | **0.6570** | 0.3573 | 0.6729 |
| 64 | G0 | full | 0.6542 | 0.6382 | 0.3645 | 0.6916 |
| 64 | G1 | frozen | 0.6088 | 0.6463 | 0.2562 | 0.4560 |
| 64 | G1 | LoRA | 0.6532 | 0.6484 | 0.3080 | 0.7019 |
| 64 | G1 | full | 0.6759 | 0.6437 | **0.4181** | **0.7610** |
| 64 | Scratch-N | full | 0.5923 | 0.6120 | 0.2026 | 0.4675 |
| 64 | Random-N | frozen | 0.4869 | 0.4820 | 0.1393 | 0.2398 |
| 64 | gRNAde | frozen | 0.6726 | 0.6005 | 0.2106 | 0.6041 |
| 64 | gRNAde | LoRA | <u>0.6766</u> | 0.6145 | 0.2763 | 0.5375 |
| 64 | gRNAde | full | 0.6550 | 0.6104 | 0.3026 | 0.6040 |
| 64 | NA-MPNN | frozen | 0.6328 | 0.6425 | 0.2522 | 0.4374 |
| 64 | NA-MPNN | LoRA | 0.6581 | 0.6472 | 0.3646 | 0.6675 |
| 64 | NA-MPNN | full | **0.6796** | <u>0.6495</u> | 0.3594 | 0.6045 |

#### 11.2 Classification and conformation results

The summary above retains all small-GNN encoder, mode and width conditions. Complete CoBRA metrics and named external-set AP are in Supplementary Data D100–D101; RNAProt metrics are in D102 and apo/holo metrics in D103–D104. The external pool retains 48 set-qualified records across 44 PDBs, with a separate no-8F0N sensitivity. The six-pair conformation analysis retains the Section 4 mappings and exposure qualifications, with 605/607 labelled apo/holo residues and 64 positives per state.

##### 11.3 Small residual RSR heads

**Table S70.** RSR residual GNN width32/64: all 22 conditions. Mean of three seed-specific metrics across four puzzles. Ranks are within head. SD is unranked.

| Head | Encoder | Mode | Target $\rho$ | Seed SD | Pooled $\rho$ |
|---|---|---|---|---|---|
| GNN | S0 | frozen | 0.4189 | 0.0370 | 0.5045 |
| GNN | S0 | LoRA | 0.3203 | 0.0370 | 0.4296 |
| GNN | S0 | full | 0.3373 | 0.2332 | 0.4360 |
| GNN | G0 | frozen | 0.3712 | 0.0649 | 0.4542 |
| GNN | G0 | LoRA | 0.3520 | 0.0412 | 0.5329 |
| GNN | G0 | full | 0.3998 | 0.0465 | 0.5447 |
| GNN | G1 | frozen | 0.3947 | 0.0835 | 0.5298 |
| GNN | G1 | LoRA | **0.4729** | 0.0505 | **0.5772** |
| GNN | G1 | full | 0.3564 | 0.0391 | 0.5283 |
| GNN | Scratch-N | full | 0.1855 | 0.1331 | 0.4862 |
| GNN | Random-N | frozen | 0.4003 | 0.0237 | 0.5528 |
| ResidualGNN64 | S0 | frozen | 0.3792 | 0.0462 | 0.5303 |
| ResidualGNN64 | S0 | LoRA | 0.3183 | 0.0367 | 0.4696 |
| ResidualGNN64 | S0 | full | 0.3101 | 0.0833 | 0.4952 |
| ResidualGNN64 | G0 | frozen | 0.3660 | 0.0478 | 0.5385 |
| ResidualGNN64 | G0 | LoRA | 0.3523 | 0.0126 | 0.5155 |
| ResidualGNN64 | G0 | full | 0.2790 | 0.1089 | 0.4690 |
| ResidualGNN64 | G1 | frozen | 0.3897 | 0.0628 | 0.5181 |
| ResidualGNN64 | G1 | LoRA | 0.3708 | 0.0368 | 0.5091 |
| ResidualGNN64 | G1 | full | 0.3236 | 0.0614 | 0.4884 |
| ResidualGNN64 | Scratch-N | full | 0.1852 | 0.0537 | 0.5378 |
| ResidualGNN64 | Random-N | frozen | **0.4374** | 0.0097 | **0.5840** |

The larger point estimate is not a validation-selected deployment claim. G1 LoRA width32 minus its original MLP target correlation is −0.0648. The conditional interval is [−0.0892, −0.0440]. Random-frozen width64 reaches 0.4374. A smaller graph readout therefore does not uniformly improve transfer or replace a matched unpretrained control.

##### 11.4 S0 results across tasks

S0 results cover ligand binding, RNAProt, RSR and RNA3D/CASP16. Figure 2F compares MLP and width32 GNN RSR results (Tables S32 and S33). Supplementary Data D106–D107 include every evaluated cohort; validation values are selection diagnostics.

**Supplementary Data D106.** S0 binary completion: all cohorts.

**Supplementary Data D107.** S0 structure-ranking completion, 20 epochs.

##### 11.5 Paired uncertainty

Small classification-GNN and S0 extension results are point estimates without paired intervals, except the matched frozen120 S0 comparison in Section 14. Released-GAT and small-RSR intervals condition on fitted seeds, are unadjusted and exclude pretraining variability. RSR resampling has only four independent puzzles.

**Supplementary Data D108.** Released-encoder GAT minus GNN, archived paired 95% intervals using 177 PDB-connected units and 10,000 draws.

**Supplementary Data D109.** All archived small-RSR-head paired target-macro contrasts, conditional unadjusted 95% intervals.

**Supplementary Data D110.** RSR width64-minus-width32 contrasts for all 11 matched conditions.

#### 12 Exploratory observed-backbone atom ablation

##### 12.1 Input definitions and matched fitting contract

G0 remains the main six-atom model. This extension tests whether additional observed backbone atoms improve transfer under the same G0 objectives. G0 uses P, C1′, C2′, C3′, C4′ and O4′. G0–8-atom adds the non-bridging phosphate oxygens OP1 and OP2. G0–12-atom additionally includes O2′, O3′, O5′ and C5′. All three use the same deterministic virtual nitrogen, sugar-based frame and edge anchors. Observed base atoms are excluded.

Inactive slots are zeroed in the same feature layout, preserving parameter count. Missing additional atoms remain masked without removing records. The six-atom coordinates, graph, residue order and labels are retained exactly.

The six-atom source is the existing G0 epoch-100 checkpoint with pretraining seed 0. The eight- and twelve-atom variants were separately pretrained to epoch 100 with seed 0 under the same G0 objectives. The atom intervention therefore applies during both pretraining and downstream fitting. Downstream fitting fixes the source checkpoint and varies three head/training seeds. Head initialization uses $91000 + s$ and training randomness uses $92000 + s$ for $s = 0, 1, 2$. Random-frozen and scratch-full controls use a fixed untrained encoder initialization with the corresponding atom input. The panel comprises three inputs, five pretrained/control regimes and four fitted tasks, giving 180 fits. SMARTPocket and CASP16 apply those fitted weights without further training. This is an exploratory extension after earlier benchmark inspection. It does not replace the main reference or establish a prospectively selected optimal atom subset.

Classification uses the 82,689-parameter width-128 GNN for 120 CoBRA or 40 RNAProt epochs; ranking uses the 16,641-parameter 128–128–1 ReLU MLP with dropout 0.1 for 20 epochs. Rates, BCE weighting, normalization, batch sizes and checkpoint selection follow Sections 1.6–1.10, with ranking gradient clipping at 1 and a 500-node sample cap. The CoBRA 331/41, RNAProt CLEAN 835/171, RSR 12,479/4,000 and RNA3D 40,138/6,120 training/validation supports are unchanged; RNA3D excludes AF3 from training. Ranking validation is evaluated every epoch.

Classification reports three-seed probability-ensemble metrics. Ranking reports the mean of seedwise correlations. Pooled AP, equal-PDB AP and mean-target Spearman remain distinct estimands. Existing memberships and exposure qualifications are unchanged. External evaluation contains 48 records from 44 PDBs, with a separate 8F0N exclusion sensitivity. Neither benchmark inspection nor downstream holo exposure is removed by changing the input atoms. This newly paired downstream panel is not an exact replay of every historical G0 fitting run, so its six-atom results must not be silently substituted into earlier tables with different source/seed contracts. In particular, LoRA adapters were initialized at a different random-number-generator state: the historical RSR pipeline inserted them after an intermediate MLP initialization under seed 91000, whereas this panel inserted them after encoder construction under seed 0. Archived records identify the same source checkpoint and matching per-seed head initialization, but different initial LoRA encoder states; LoRA rank 4, $\alpha = 8$ and head/training seeds are unchanged.

#### 12.2 Results and interpretation

The six-atom LoRA reference has RSR mean-target Spearman 0.4547, compared with 0.5107 for the historical fit in the main panel (Table S32). Both average seed-specific correlations. These are different adapter-initialization realizations, not exact replays. The isolated contribution of this initialization difference to the score gap was not tested.

Under full adaptation, twelve-atom external pooled AP is 0.6782 versus 0.6686 for G0, with TE18 AP 0.5856 versus 0.5472. Frozen external AP instead decreases from 0.6274 to 0.6110. Full RNAProt AP is similar at 0.6406 versus 0.6415. Twelve-atom LoRA improves RNA3D38 mean-target Spearman from 0.3438 to 0.4030, whereas full-adaptation CASP16 correlation decreases from 0.2857 to 0.2206 and apo AUROC from 0.8210 to 0.7802. RSR LoRA values are close at 0.4555 and 0.4547. Thus additional backbone atoms have task- and adaptation-dependent effects in these point estimates. They do not establish universal twelve-atom superiority or six-atom optimality. All unpretrained controls are retained, including the RNA3D frozen comparison where six-atom random features exceed pretrained features (0.3097 versus 0.2902).

No paired confidence intervals were computed for the atom-input panel. Seed SD measures downstream fitting variability conditional on one pretraining seed; it does not establish significance or equivalence. These exploratory tables are unranked.

**Table S71.** Observed-backbone input comparison. AP is pooled average precision of three-seed probability ensembles. $\rho$ is mean-target Spearman averaged over three individually scored seeds. G0 is the six-atom reference. All five pretrained/control regimes are shown for every input. RNA3D and CASP denote RNA3D38 and CASP16.

| Model | Mode | TR60 AP | Ext. AP | Prot. AP | RSR $\rho$ | RNA3D $\rho$ | CASP $\rho$ |
|---|---|---|---|---|---|---|---|
| Pretrained / Frozen | | | | | | | |
| G0 (6-atom) | Frozen | 0.8031 | 0.6274 | 0.6471 | 0.3107 | 0.2902 | 0.1185 |
| G0–8-atom | Frozen | 0.7894 | 0.6138 | 0.6170 | 0.3452 | 0.2891 | 0.1121 |
| G0–12-atom | Frozen | 0.7873 | 0.6110 | 0.6263 | 0.4043 | 0.3363 | 0.1913 |
| Pretrained / LoRA | | | | | | | |
| G0 (6-atom) | LoRA | 0.8270 | 0.6534 | 0.6402 | 0.4547 | 0.3438 | 0.1794 |
| G0–8-atom | LoRA | 0.8261 | 0.6553 | 0.6380 | 0.4115 | 0.3917 | 0.2349 |
| G0–12-atom | LoRA | 0.8329 | 0.6701 | 0.6463 | 0.4555 | 0.4030 | 0.2223 |
| Pretrained / Full | | | | | | | |
| G0 (6-atom) | Full | 0.8338 | 0.6686 | 0.6415 | 0.4270 | 0.3120 | 0.2857 |
| G0–8-atom | Full | 0.8264 | 0.6557 | 0.6442 | 0.3890 | 0.3396 | 0.2545 |
| G0–12-atom | Full | 0.8358 | 0.6782 | 0.6406 | 0.4536 | 0.3296 | 0.2206 |
| Unpretrained control / Frozen | | | | | | | |
| Random (6-atom) | Frozen | 0.6547 | 0.5033 | 0.5160 | 0.2249 | 0.3097 | 0.1087 |
| Random (8-atom) | Frozen | 0.6377 | 0.4834 | 0.4996 | 0.2219 | 0.3228 | 0.0715 |
| Random (12-atom) | Frozen | 0.6064 | 0.4869 | 0.4919 | 0.2598 | 0.3455 | 0.0951 |
| Unpretrained control / Full | | | | | | | |
| Scratch (6-atom) | Full | 0.8066 | 0.6346 | 0.6225 | 0.2737 | 0.2103 | 0.0310 |
| Scratch (8-atom) | Full | 0.8103 | 0.6378 | 0.6077 | 0.2793 | 0.2227 | 0.1209 |
| Scratch (12-atom) | Full | 0.7903 | 0.6179 | 0.6075 | 0.3029 | 0.2946 | 0.2013 |

##### 12.3 Detailed results

Supplementary Data D112–D113 give all classification metrics and named external-set AP. D114 retains apo/holo metrics using the same TR60-fitted weights and validation thresholds. D115 reports RSR puzzle-level correlations; D116 reports RNA3D38 and CASP16 ranking metrics. These files include all five pretrained/control regimes for each atom input.

#### 13 Released-encoder RSR width32 comparison

##### 13.1 Fitting protocol

Released gRNAde and NA-MPNN use frozen, LoRA and full adaptation with fixed checkpoints and three downstream seeds. Their adapters, native dropout and exposure qualifications follow Section 8. The shared readout is the 8,513-parameter two-block residual width32 GNN with LayerNorm, dropout 0.1, C4′-kNN16 edges and scalar mean pooling.

The RSR recipe in Section 1.10 is retained for 20 epochs, with gradient clipping at 1, a 500-node sample cap and earliest maximum validation mean-target Spearman selection. The split remains 12,479/4,000/4,000 decoys, with 1,000 per test Puzzle 18–21. Head initialization follows the native downstream-seed design. Metrics average three seed-specific correlations; the four puzzles remain the independent test units. Inputs and pretraining histories remain unmatched, and this comparison is exploratory.

**Supplementary Data D117.** Released-encoder RSR width32 results.

**Supplementary Data D118.** All 18 public width32 evaluations and their selected-validation metrics.

##### 13.2 Puzzle-specific behavior and controls

**Supplementary Data D119.** All width32 conditions on the same four RSR puzzles, with 1,000 decoys per puzzle.

G1/LoRA has the largest GNN mean-target point estimate (0.4729), but the MLP G1/LoRA value remains higher at 0.5377. Random-frozen GNN correlation 0.4003 exceeds frozen G0/G1 values 0.3712/0.3947, so the graph head can extract ranking information without pretraining. G0/full exceeds scratch-full numerically, while gRNAde/full has negative mean correlation despite successful fitting. All conditions and negative values are retained. These observations do not identify the cause of the gRNAde adaptation result or establish a universal architecture ranking.

##### 13.3 Paired target-level uncertainty

Public-minus-native comparisons match adaptation mode and head. Paired bootstrap intervals use 10,000 draws of the same four puzzles after averaging the three seed correlations within each puzzle. These intervals are conditional and unadjusted; only 35 unordered resample-count patterns are possible. Seed SD describes fitting variability, a separate uncertainty axis.

G1 minus NA-MPNN under LoRA is 0.1206 with a conditional interval of [0.0715, 0.1573]. G0 minus NA-MPNN under full adaptation is 0.0868 with interval [-0.0923, 0.2659], leaving the difference unresolved. G0 minus gRNAde under full adaptation is 0.4649 with interval [0.2307, 0.6442]. These illustrative differences do not turn this post-inspection four-target panel into a confirmatory comparison.

**Table S72.** Public minus reference under the same adaptation mode and width32 head. Values are paired target-bootstrap differences and 95% percentile intervals, with 10,000 draws and seed 20261022. They are conditional on the fitted seeds, descriptive and unadjusted for multiple comparisons.

| Public encoder | Mode | Reference | $\Delta\rho$ | Lower | Upper |
|---|---|---|---|---|---|
| gRNAde | Frozen | S0 | -0.1573 | -0.3434 | 0.0477 |
| gRNAde | Frozen | G0 | -0.1095 | -0.2240 | 0.0049 |
| gRNAde | Frozen | G1 | -0.1330 | -0.2874 | 0.0214 |
| gRNAde | Frozen | Random-frozen | -0.1386 | -0.2938 | 0.0345 |
| gRNAde | LoRA | S0 | -0.1106 | -0.2670 | -0.0120 |
| gRNAde | LoRA | G0 | -0.1423 | -0.2914 | 0.0067 |
| gRNAde | LoRA | G1 | -0.2632 | -0.3705 | -0.1559 |
| gRNAde | Full | S0 | -0.4025 | -0.5897 | -0.2780 |
| gRNAde | Full | G0 | -0.4649 | -0.6442 | -0.2307 |
| gRNAde | Full | G1 | -0.4216 | -0.6357 | -0.2663 |
| gRNAde | Full | Scratch-N | -0.2506 | -0.4845 | -0.0869 |
| NA-MPNN | Frozen | S0 | -0.1884 | -0.2786 | -0.1080 |
| NA-MPNN | Frozen | G0 | -0.1407 | -0.1942 | -0.0541 |
| NA-MPNN | Frozen | G1 | -0.1641 | -0.2443 | -0.0470 |
| NA-MPNN | Frozen | Random-frozen | -0.1698 | -0.3113 | 0.0279 |
| NA-MPNN | LoRA | S0 | 0.0320 | -0.1061 | 0.2208 |
| NA-MPNN | LoRA | G0 | 0.0003 | -0.0899 | 0.0904 |
| NA-MPNN | LoRA | G1 | -0.1206 | -0.1573 | -0.0715 |
| NA-MPNN | Full | S0 | -0.0243 | -0.0986 | 0.0872 |
| NA-MPNN | Full | G0 | -0.0868 | -0.2659 | 0.0923 |
| NA-MPNN | Full | G1 | -0.0434 | -0.1357 | 0.0452 |
| NA-MPNN | Full | Scratch-N | 0.1276 | 0.0370 | 0.1935 |

### 14 S0 frozen readout at a matched 120-epoch budget

#### 14.1 Matched fitting protocol

Frozen S0 uses the same 331/41 TR60 training/validation chains and 120-epoch protocol as G0/G1/random-frozen. The epoch-100/seed-0 encoder supplies raw embeddings to the 82,689-parameter GNN, with Adam at 0.001, weight decay 0.0001 and dropout 0.3. Three downstream seeds use matched head initialization. Earliest maximum validation AP, evaluated every five epochs, selects epochs 95/35/90. The 120-epoch budget was chosen to match the comparison protocol rather than by test performance.

#### 14.2 Individual fits and all-cohort metrics

**Supplementary Data D121.** Frozen 120-epoch fits.

Supplementary Data D106 reports this S0 ensemble on all ten cohorts. Its validation-selected threshold is 0.56. Apo/holo results use the same six target pairs, but the ligand-test uncertainty analysis below does not supply apo/holo intervals.

#### 14.3 Matched-duration paired analysis

Probabilities are averaged across the three downstream seeds before scoring. TR60 test, pooled external and no-8F0N external contain 43, 44 and 43 independent PDBs, respectively. The full external pool retains all 48 set-qualified records. Repeated PDBs are resampled together, not deduplicated by chain identifier. The paired percentile bootstrap uses 10,000 draws and seed 20261001 with identical multiplicities for all four models and both estimands. All draws are defined. No model, epoch, cohort or threshold is selected using these contrasts.

**Table S73.** Exploratory S0 frozen120 paired contrasts. The same 10,000 PDB bootstrap draws produce pooled-residue and equal-PDB-macro estimands. Intervals condition on these fitted models and are unadjusted for multiplicity. They do not quantify retraining variation or demonstrate equivalence when zero is included.

| Cohort | Contrast | Δpooled AP | 95% interval | Δmacro AP | 95% interval |
|---|---|---|---|---|---|
| TR60 test | S0 − G0 | -0.0163 | [-0.0398, +0.0024] | -0.0474 | [-0.0734, -0.0229] |
| TR60 test | S0 − G1 | +0.0056 | [-0.0194, +0.0265] | -0.0171 | [-0.0431, +0.0088] |
| TR60 test | S0 − Random-frozen | +0.1321 | [+0.0737, +0.1838] | +0.0567 | [+0.0045, +0.1098] |
| External | S0 − G0 | +0.0002 | [-0.0258, +0.0251] | -0.0116 | [-0.0368, +0.0134] |
| External | S0 − G1 | +0.0126 | [-0.0115, +0.0374] | -0.0021 | [-0.0218, +0.0178] |
| External | S0 − Random-frozen | +0.1243 | [+0.0680, +0.1802] | +0.0805 | [+0.0415, +0.1201] |
| No 8F0N | S0 − G0 | +0.0034 | [-0.0230, +0.0286] | -0.0104 | [-0.0367, +0.0159] |
| No 8F0N | S0 − G1 | +0.0183 | [-0.0059, +0.0418] | -0.0007 | [-0.0215, +0.0195] |
| No 8F0N | S0 − Random-frozen | +0.1286 | [+0.0714, +0.1869] | +0.0827 | [+0.0429, +0.1235] |

S0 improves on random-frozen under both AP estimands in all three reported cohorts. Relative to G0, both external estimands remain unresolved. TR60 test PDB-macro AP favors G0, whereas its pooled AP contrast remains unresolved. Thus the external proximity supports transfer without explicit geometric auxiliary losses but is not evidence of equivalence or uniformly preserved performance. These post-hoc, conditional intervals do not include source-seed or retraining uncertainty.

### 15 Native RNAProt baselines on the CLEAN split

#### 15.1 Cohort and native inputs

RGCN and DualEquiNet are fitted from random initialization on the same 835 training and 171 validation graphs used for the BRIDGE RNAProt head panel. The locked test set contains 191 graphs, 10,780 residues and 2,980 positive labels across 177 PDB-connected units. All original retained residue IDs, ordering, labels and the 500-node sample limit are preserved. This is an exploratory comparator addition after prior benchmark inspection, not a newly independent benchmark. Test results do not select the six checkpoints, architecture or thresholds.

RGCN receives native A/U/C/G one-hot features and the directed RNAglib graph with 20 backbone and Leontis–Westhof relation types. DualEquiNet receives the same nucleotide features and graph adjacency without edge types, together with phosphate coordinates. The original graph is induced on the retained residue support, with unsupported relation labels removed as in RNAglib. No BRIDGE kNN edges are substituted. Native base-pair relations can encode base geometry, so these methods are not base-blind input controls. Despite its implementation class name, the DualEquiNet RNAProt model operates on nucleotide nodes, not all-heavy-atom nodes.

The three native T annotations at 6VRD chain C residues 3, 7 and 15 use RNAglib's unknown-base zero vector; BRIDGE normalizes T to U. Residue support is unchanged. RNAProt macro AP remains undefined because 55 of 177 test units have no positives.

#### 15.2 Architecture, optimization and selection

The RGCN uses the installed RNAglib PygModel with four width128 relational layers, dropout 0.2 and 1,378,817 parameters. Its architecture follows the RNAglib benchmark Appendix F specification [11]. Training uses Adam with learning rate 0.001, batch size 8 and binary cross-entropy weighted by the square root of the training negative-to-positive ratio, approximately 1.69. This differs from alternative configurations in the released example scripts and is not claimed as an exact replay of the published result.

DualEquiNet uses the released DualEquiNet Bi CSIP Atom class at commit 154682846465 [12]. The model has hidden width32, two atom-level layers, four heads, maximum spherical-harmonic degree 2 and 60,336 parameters. Its spherical-harmonic cutoff is 0.99, maximum neighbor count is 32 and neighbor updates are disabled. Training uses Adam with learning rate 0.0005, batch size 32 and unweighted binary cross-entropy. These settings follow the pinned native runner defaults, not a recovered tuned RNAProt configuration. Its native default training duration is much longer than this fixed-budget comparison.

Both models use zero weight decay, FP32, no mixed precision, no scheduler and no gradient clipping. Three seeds complete all 40 epochs without early stopping. Validation pooled AP is evaluated every five epochs, selecting the earliest maximum among epochs 5, 10, . . ., 40. Epoch1 is diagnostic only. RGCN selects epochs 5/10/5 and DualEquiNet selects 40/40/30 for seeds 0/1/2. The latter two boundary selections limit conclusions about an optimal training duration. Selection does not use test labels. Native-model seeds vary initialization and fitting, unlike BRIDGE's fixed pretrained-encoder seed with varied downstream seeds.

The test probability ensemble averages the three selected models before scoring. Validation MCC selects a threshold from 0.02, 0.04, . . ., 0.98, with the smallest tied threshold. The ensemble thresholds are 0.60 for RGCN and 0.34 for DualEquiNet. Individual-seed thresholds are reported separately. At 0.5, DualEquiNet predicts only 51 positive residues, compared with 2,980 observed positives. At its validation-fixed threshold it predicts 3,233.

Consequently, its accuracy is close to the all-negative value while its fixed and validation-threshold MCC values differ substantially. Test-label oracle MCC remains a diagnostic only.

**Supplementary Data D123.** Native-model CLEAN test results.

**Supplementary Data D124.** All six native-model test evaluations.

##### 15.3 Conditional uncertainty

Paired percentile-bootstrap intervals use 10,000 shared draws of the 177 PDB-connected units, with seed 20261026. These describe test-cohort uncertainty conditional on the fitted models, not retraining variability. They are exploratory and unadjusted for multiple comparisons. RGCN exceeds DualEquiNet in the paired AP and AUROC contrasts under these fixed recipes. No native-model-versus-BRIDGE paired interval has been computed, and numerical cross-model differences do not establish an architecture-specific pretraining effect.

**Table S74.** Conditional intervals for the two native-model probability ensembles and their paired difference. The same 10,000 bootstrap draws resample 177 PDB-connected units, with seed 20261026. Intervals are exploratory and unadjusted for multiplicity. All draws are defined. No new paired native-model-versus-BRIDGE interval is computed.

| Model or contrast | Metric | Estimate | 95% interval |
|---|---|---|---|
| RGCN | AP | 0.4944 | [0.4243, 0.5646] |
| RGCN | AUROC | 0.7241 | [0.6863, 0.7633] |
| RGCN | MCC at 0.5 | 0.3160 | [0.2562, 0.3801] |
| RGCN | Oracle MCC | 0.3323 | [0.2697, 0.4060] |
| DualEquiNet | AP | 0.4288 | [0.3663, 0.4927] |
| DualEquiNet | AUROC | 0.6771 | [0.6409, 0.7141] |
| DualEquiNet | MCC at 0.5 | 0.0239 | [-0.0048, 0.0577] |
| DualEquiNet | Oracle MCC | 0.2665 | [0.2132, 0.3305] |
| RGCN − DualEquiNet | AP | +0.0657 | [+0.0405, +0.0929] |
| RGCN − DualEquiNet | AUROC | +0.0470 | [+0.0311, +0.0634] |
| RGCN − DualEquiNet | MCC at 0.5 | +0.2921 | [+0.2279, +0.3563] |
| RGCN − DualEquiNet | Oracle MCC | +0.0657 | [+0.0429, +0.0871] |

RGCN uses PyTorch 2.9.1/CUDA 12.8 and DualEquiNet uses PyTorch 2.9.0/CUDA 12.8 on RTX 5070 Ti GPUs.

#### 16 Sequence-only control and optimization sensitivity

##### 16.1 Matched removal of SASA supervision

Sequence-only retains S0's architecture, six observed backbone atoms, virtual N, paired initialization, membership and optimizer. Only sequence cross-entropy with label smoothing 0.05 is active; SASA, $\chi$ and PairGeom weights are zero. Auxiliary modules remain instantiated to preserve initialization and random-number behavior. The objective therefore uses no observed base-coordinate targets, although inputs come from experimental atomic structures.

Three pretraining seeds run for 100 epochs on 2,501 records, with fixed terminal checkpoints and target-free soft-prefix argmax decoding. Neither native sequence nor $\chi$ enters decoder context. Diagnostics cover all 68 records, the 67-record 3GS5 exclusion sensitivity and the existing 26-record public-inventory-unmatched subset. S0 and other objective-removal results use the same decoding and cohort definitions.

**Table S75.** Source recovery averages three pretraining-seed metrics. Macro weights RNA records equally. SD describes the three seed-specific pooled values, not cohort uncertainty.

| Subset | Model | Pooled (%) | Macro (%) | Pooled SD (pp) |
|---|---|---|---|---|
| All 68 | Sequence-only | 60.88 | 58.61 | 2.21 |
| All 68 | S0 | 62.19 | 59.83 | 0.12 |
| All 68 | S0+PairGeom | 62.45 | 60.39 | 0.93 |
| All 68 | S0+$\chi$ | 62.76 | 60.43 | 0.55 |
| All 68 | G0 | 62.38 | 60.05 | 0.74 |
| All 68 | G1 | 61.81 | 59.47 | 0.50 |
| No 3GS5 | Sequence-only | 60.92 | 58.67 | 2.19 |
| No 3GS5 | S0 | 62.22 | 59.86 | 0.17 |
| No 3GS5 | S0+PairGeom | 62.41 | 60.25 | 0.89 |
| No 3GS5 | S0+$\chi$ | 62.76 | 60.40 | 0.58 |
| No 3GS5 | G0 | 62.38 | 60.02 | 0.74 |
| No 3GS5 | G1 | 61.79 | 59.39 | 0.54 |
| Inventory 26 | Sequence-only | 58.30 | 56.88 | 1.63 |
| Inventory 26 | S0 | 59.75 | 58.75 | 0.48 |
| Inventory 26 | S0+PairGeom | 60.40 | 59.17 | 0.79 |
| Inventory 26 | S0+$\chi$ | 60.27 | 58.75 | 0.84 |
| Inventory 26 | G0 | 61.30 | 60.14 | 0.63 |
| Inventory 26 | G1 | 59.71 | 58.41 | 0.34 |

**Supplementary Data D127.** Every pretraining seed under the same target-free soft-prefix decoding and frozen source-test membership.

**Table S76.** Paired recovery contrasts from 20,000 draws of PDB/sequence/structure-cluster components. Full68 and no3GS5 contain 11 units, inventory26 contains six. The full-cohort dependence closure is retained after exclusions. Intervals condition on the fitted seeds and decoding orders, exclude training-seed uncertainty and are not multiplicity-adjusted.

| Subset | Contrast | Metric | Δ(pp) | 95% interval (pp) |
|---|---|---|---|---|
| All 68 | Seq. − S0 | Pooled | -1.31 | [-2.82, 0.57] |
| All 68 | Seq. − S0 | Macro | -1.22 | [-2.95, 0.33] |
| All 68 | Seq. − G0 | Pooled | -1.50 | [-3.09, 0.26] |
| All 68 | Seq. − G0 | Macro | -1.43 | [-3.39, 0.20] |
| No 3GS5 | Seq. − S0 | Pooled | -1.30 | [-2.81, 0.57] |
| No 3GS5 | Seq. − S0 | Macro | -1.18 | [-2.93, 0.33] |
| No 3GS5 | Seq. − G0 | Pooled | -1.46 | [-3.04, 0.28] |
| No 3GS5 | Seq. − G0 | Macro | -1.34 | [-3.30, 0.21] |
| Inventory 26 | Seq. − S0 | Pooled | -1.44 | [-2.74, -0.29] |
| Inventory 26 | Seq. − S0 | Macro | -1.87 | [-3.16, -0.54] |
| Inventory 26 | Seq. − G0 | Pooled | -3.00 | [-4.58, -1.14] |
| Inventory 26 | Seq. − G0 | Macro | -3.26 | [-4.61, -1.91] |

Sequence-only pooled recovery is 60.88%, compared with 62.19% for S0 and 62.38% for G0. Full-cohort paired differences remain unresolved. Individual sequence-only values are 59.73%, 59.48% and 63.43%, showing variability that the conditional cohort intervals do not capture. The inventory26 sensitivity favors S0 and G0 over sequence-only under the fixed-model component bootstrap, but contains only six dependence components. It cannot establish a general ranking across pretraining seeds or unseen RNA families. The inactive angle head is not scored.

#### 16.2 Frozen transfer without SASA targets

Frozen transfer fixes pretraining seed 0 and varies three head seeds with the 82,689-parameter GNN. CoBRA uses 120 epochs and RNAProt 40, matching S0's readout and optimization protocol. CoBRA evaluation includes TR60 test43, external48 and no-8F0N; RNAProt retains 191 graphs/177 PDB-connected units, with strict macro AP undefined because 55 units lack positives.

Sequence-only exceeds random-frozen in pooled AP on external and RNAProt test, supporting transfer without SASA supervision. S0-minus-sequence-only external pooled AP is 0.0268 and unresolved, whereas PDB-macro AP is 0.0288 with a positive paired interval. RNAProt pooled difference 0.0087 is unresolved. These comparisons do not establish equivalence or absence of SASA utility. Downstream fitting uses source seed0 only. The other two pretraining seeds support recovery replication, not downstream source-seed replication. Released gRNAde is a SASA-free contextual comparator, not this matched removal, because its inputs and training history differ.

**Table S77.** Frozen GNN transfer using three-head-seed probability ensembles. Sequence-only source seed0 was fixed before downstream fitting. CoBRA uses120 epochs and RNAProt40. Archived S0/G0/random controls use matched readout contracts. Undefined RNAProt macro AP is retained as a dash.

| Task | Cohort | Encoder | AP | Macro AP | AUROC | MCC .5 / val / oracle |
|---|---|---|---|---|---|---|
| CoBRA | JL10 | Sequence-only | 0.6105 | 0.6435 | 0.7382 | 0.3239 / 0.3111 / 0.3546 |
| CoBRA | JL10 | S0 | 0.5926 | 0.6264 | 0.7314 | 0.3684 / 0.3911 / 0.3989 |
| CoBRA | JL10 | G0 | 0.6131 | 0.6468 | 0.7486 | 0.3361 / 0.3908 / 0.3979 |
| CoBRA | JL10 | Random-frozen | 0.4658 | 0.5273 | 0.6600 | 0.2459 / 0.2459 / 0.2821 |
| CoBRA | RB9 | Sequence-only | 0.6963 | 0.7410 | 0.8091 | 0.4563 / 0.4653 / 0.5169 |
| CoBRA | RB9 | S0 | 0.7251 | 0.8005 | 0.8241 | 0.4596 / 0.4773 / 0.5372 |
| CoBRA | RB9 | G0 | 0.7568 | 0.8333 | 0.8440 | 0.5152 / 0.5755 / 0.5808 |
| CoBRA | RB9 | Random-frozen | 0.4822 | 0.6990 | 0.6613 | 0.1997 / 0.1997 / 0.3133 |
| CoBRA | TE18 | Sequence-only | 0.4691 | 0.5569 | 0.6284 | 0.1828 / 0.1961 / 0.2259 |
| CoBRA | TE18 | S0 | 0.5342 | 0.5976 | 0.7025 | 0.3202 / 0.3168 / 0.3393 |
| CoBRA | TE18 | G0 | 0.4927 | 0.6013 | 0.6634 | 0.2660 / 0.2698 / 0.2902 |
| CoBRA | TE18 | Random-frozen | 0.4335 | 0.5671 | 0.5966 | 0.1513 / 0.1513 / 0.1768 |
| CoBRA | TL12 | Sequence-only | 0.7197 | 0.7285 | 0.7811 | 0.4882 / 0.4800 / 0.5079 |
| CoBRA | TL12 | S0 | 0.7505 | 0.7842 | 0.8259 | 0.5590 / 0.5677 / 0.5856 |
| CoBRA | TL12 | G0 | 0.7702 | 0.7674 | 0.8023 | 0.5124 / 0.5118 / 0.5313 |
| CoBRA | TL12 | Random-frozen | 0.6825 | 0.6836 | 0.8044 | 0.4610 / 0.4610 / 0.4846 |
| CoBRA | TR60 test | Sequence-only | 0.7900 | 0.7703 | 0.8507 | 0.5428 / 0.5856 / 0.5997 |
| CoBRA | TR60 test | S0 | 0.7868 | 0.7558 | 0.8569 | 0.5561 / 0.5708 / 0.5900 |
| CoBRA | TR60 test | G0 | 0.8031 | 0.8031 | 0.8696 | 0.5912 / 0.6277 / 0.6404 |
| CoBRA | TR60 test | Random-frozen | 0.6547 | 0.6990 | 0.7804 | 0.3818 / 0.3818 / 0.4546 |
| CoBRA | No 8F0N | Sequence-only | 0.5885 | 0.6368 | 0.7251 | 0.3386 / 0.3358 / 0.3667 |
| CoBRA | No 8F0N | S0 | 0.6162 | 0.6664 | 0.7575 | 0.4047 / 0.4157 / 0.4209 |
| CoBRA | No 8F0N | G0 | 0.6128 | 0.6768 | 0.7484 | 0.3793 / 0.4121 / 0.4121 |
| CoBRA | No 8F0N | Random-frozen | 0.4876 | 0.5837 | 0.6687 | 0.2470 / 0.2470 / 0.2732 |
| CoBRA | External | Sequence-only | 0.6008 | 0.6435 | 0.7317 | 0.3480 / 0.3487 / 0.3765 |
| CoBRA | External | S0 | 0.6276 | 0.6724 | 0.7638 | 0.4139 / 0.4256 / 0.4301 |
| CoBRA | External | G0 | 0.6274 | 0.6840 | 0.7551 | 0.3899 / 0.4227 / 0.4227 |
| CoBRA | External | Random-frozen | 0.5033 | 0.5918 | 0.6788 | 0.2600 / 0.2600 / 0.2855 |
| RNAProt | Test | Sequence-only | 0.6329 | – | 0.7986 | 0.3943 / 0.4088 / 0.4183 |
| RNAProt | Test | S0 | 0.6415 | – | 0.7983 | 0.4101 / 0.4141 / 0.4202 |
| RNAProt | Test | G0 | 0.6471 | – | 0.8003 | 0.4102 / 0.4178 / 0.4216 |
| RNAProt | Test | Random-frozen | 0.5160 | – | 0.6728 | 0.2226 / 0.2701 / 0.2917 |

**Supplementary Data D130.** Individual sequence-only readout seeds, including validation and each scored held-out cohort.

**Table S78.** Paired 10,000-draw PDB bootstrap contrasts, conditional on fitted models and unadjusted. Differences are sequence-only minus the named control. No source-seed replication of downstream fitting is implied.

| Task | Cohort | Contrast | Metric | Estimate | 95% interval |
|---|---|---|---|---|---|
| CoBRA | TR60 test | Seq. − S0 | Pooled AP | 0.0032 | [-0.0181, 0.0258] |
| CoBRA | TR60 test | Seq. − S0 | Macro AP | 0.0146 | [-0.0160, 0.0465] |
| CoBRA | TR60 test | Seq. − Random | Pooled AP | 0.1352 | [0.0770, 0.1880] |
| CoBRA | TR60 test | Seq. − Random | Macro AP | 0.0713 | [0.0238, 0.1184] |
| CoBRA | TR60 test | Seq. − G0 | Pooled AP | -0.0132 | [-0.0398, 0.0099] |
| CoBRA | TR60 test | Seq. − G0 | Macro AP | -0.0328 | [-0.0587, -0.0075] |
| CoBRA | No 8F0N | Seq. − S0 | Pooled AP | -0.0277 | [-0.0745, 0.0139] |
| CoBRA | No 8F0N | Seq. − S0 | Macro AP | -0.0295 | [-0.0608, -0.0018] |
| CoBRA | No 8F0N | Seq. − Random | Pooled AP | 0.1009 | [0.0335, 0.1637] |
| CoBRA | No 8F0N | Seq. − Random | Macro AP | 0.0532 | [0.0135, 0.0937] |
| CoBRA | No 8F0N | Seq. − G0 | Pooled AP | -0.0242 | [-0.0691, 0.0182] |
| CoBRA | No 8F0N | Seq. − G0 | Macro AP | -0.0399 | [-0.0705, -0.0108] |
| CoBRA | External | Seq. − S0 | Pooled AP | -0.0268 | [-0.0703, 0.0143] |
| CoBRA | External | Seq. − S0 | Macro AP | -0.0288 | [-0.0578, -0.0018] |
| CoBRA | External | Seq. − Random | Pooled AP | 0.0975 | [0.0329, 0.1592] |
| CoBRA | External | Seq. − Random | Macro AP | 0.0517 | [0.0112, 0.0914] |
| CoBRA | External | Seq. − G0 | Pooled AP | -0.0265 | [-0.0688, 0.0150] |
| CoBRA | External | Seq. − G0 | Macro AP | -0.0404 | [-0.0699, -0.0122] |
| RNAProt | Test | Seq. − S0 | Pooled AP | -0.0087 | [-0.0238, 0.0060] |
| RNAProt | Test | Seq. − S0 | Macro AP | – | – |
| RNAProt | Test | Seq. − Random | Pooled AP | 0.1169 | [0.0547, 0.1898] |
| RNAProt | Test | Seq. − Random | Macro AP | – | – |
| RNAProt | Test | Seq. − G0 | Pooled AP | -0.0143 | [-0.0339, 0.0033] |
| RNAProt | Test | Seq. − G0 | Macro AP | – | – |

#### 16.3 Held-out RNAProt learning-rate sensitivity

The validation development analysis selected encoder rate $10^{-5}$ for both G0 and scratch by mean validation AP. We evaluated those six validation-selected checkpoints once, without further fitting or threshold selection on test. The retained primary rate is$10^{-4}$. At the lower rate, G0/scratch ensemble AP is 0.6478/0.6120, with paired difference 0.0358. The within-model learning-rate differences remain unresolved. Scratch individual-seed AP variability is smaller at the lower rate, but individual means and probability-ensemble AP are different estimands.

**Table S79.** RNAProt full-adaptation sensitivity on the unchanged CLEAN191 test. Head learning rate remains0.001, duration40 epochs and source seed0. Three-seed probabilities are averaged. The main encoder rate remains0.0001.

| Model | Encoder LR | AP | AUROC | MCC .5 / val / oracle |
|---|---|---|---|---|
| G0 | 1e-4 | 0.6415 | 0.7868 | 0.3893 / 0.3987 / 0.4162 |
| Scratch-N | 1e-4 | 0.6225 | 0.7762 | 0.3379 / 0.3851 / 0.4020 |
| G0 | 1e-5 | 0.6478 | 0.7956 | 0.4112 / 0.4188 / 0.4230 |
| Scratch-N | 1e-5 | 0.6120 | 0.7619 | 0.3483 / 0.3781 / 0.3912 |

**Supplementary Data D133.** Individual downstream-seed AP, its mean and sample SD.

**Table S80.** Paired AP intervals from10,000 draws of177 PDB-connected units. All results are conditional and exploratory. Test outcomes do not change the main learning rate.

| Contrast | Metric | Estimate | 95% interval |
|---|---|---|---|
| G0 − Scratch, $10^{-4}$ | AP | 0.0190 | [0.0028, 0.0363] |
| G0 − Scratch, $10^{-5}$ | AP | 0.0358 | [0.0145, 0.0614] |
| G0, $10^{-5} - 10^{-4}$ | AP | 0.0063 | [-0.0119, 0.0270] |
| Scratch, $10^{-5} - 10^{-4}$ | AP | -0.0105 | [-0.0294, 0.0078] |

These controls were introduced after initial benchmark inspection and remain exploratory. Paired intervals condition on fitted models and are unadjusted for multiplicity.

### 17 AI assistance

AI-assisted tools supported code development, manuscript drafting and revision, analysis presentation and internal critical review. Outputs were checked against archived computational evidence. The authors retain responsibility for the scientific claims, analyses, references and final text.
